# Inferring the Total-Evidence Timescale of Marattialean Fern Evolution in the Face of Model Sensitivity

**DOI:** 10.1101/2020.09.25.313643

**Authors:** Michael R. May, Dori L. Contreras, Michael A. Sundue, Nathalie S. Nagalingum, Cindy V. Looy, Carl J. Rothfels

## Abstract

Phylogenetic divergence-time estimation has been revolutionized by two recent developments: 1) total-evidence dating (or “tip-dating”) approaches that allow for the incorporation of fossils as tips in the analysis, with their phylogenetic and temporal relationships to the extant taxa inferred from the data, and 2) the fossilized birth-death (FBD) class of tree models that capture the processes that produce the tree (speciation, extinction, and fossilization), and thus provide a coherent and biologically interpretable tree prior. To explore the behaviour of these methods, we apply them to marattialean ferns, a group that was dominant in Carboniferous landscapes prior to declining to its modest extant diversity of slightly over 100 species. We show that tree models have a dramatic influence on estimates of both divergence times and topological relationships. This influence is driven by the strong, counter-intuitive informativeness of the uniform tree prior and the inherent nonidentifiability of divergence-time models. In contrast to the strong influence of the tree models, we find minor effects of differing the morphological transition model or the morphological clock model. We compare the performance of a large pool of candidate models using a combination of posterior-predictive simulation and Bayes factors. Notably, an FBD model with epoch-specific speciation and extinction rates was strongly favored by Bayes factors. Our best-fitting model infers stem and crown divergences for the Marattiales in the mid-Devonian and Late Cretaceous, respectively, with elevated speciation rates in the Mississippian and elevated extinction rates in the Cisuralian leading to a peak diversity of ∼2800 species at the end of the Carboniferous, representing the heyday of the Psaroniaceae. This peak is followed by the rapid decline and ultimate extinction of the Psaroniaceae, with their descendants, the Marattiaceae, persisting at approximately stable levels of diversity until the present. This general diversification pattern appears to be insensitive to potential biases in the fossil record; despite the preponderance of available fossils being from Pennsylvanian coal balls, incorporating fossilization-rate variation does not improve model fit. In addition, by incorporating temporal data directly within the model and allowing for the inference of the phylogenetic position of the fossils, our study makes the surprising inference that the clade of extant Marattiales is relatively young, younger than any of the fossils historically thought to be congeneric with extant species. This result is a dramatic demonstration of the dangers of node-based approaches to divergence-time estimation, where the assignment of fossils to particular clades are made *a priori* (earlier node-based studies that constrained the minimum ages of extant genera based on these fossils resulted in much older age estimates than in our study) and of the utility of explicit models of morphological evolution and lineage diversification.

---

The ability to infer phylogenies with branch lengths in units of time (“divergence-time estimation”) is an extremely powerful tool of evolutionary biology. Beyond simply allowing for the inference of the timing of evolutionary divergences, it enables studies of diversification rates and rates of molecular evolution, permits testing of the drivers of global patterns of biodiversity and biogeography (*e*.*g*., the roles of vicariance and dispersal), and allows us to examine the evolutionary impact of major events in the Earth’s history. Extensive research over the past two decades has dramatically improved our ability to infer time-scaled phylogenies (see Donoghue and Yang 2016), such that researchers today can choose from a wide variety of models that relax the molecular clock (Sanderson 1997; Thorne et al. 1998; Magallón 2004; Drummond et al. 2006; Drummond and Suchard 2010; Lartillot et al. 2016) and from sophisticated methods for associating fossil data with nodes on a phylogeny (Marshall 2008; Ho and Phillips 2009; Heath 2012; Rothfels et al. 2015a). Molecular dating techniques, however, are fraught with controversy, and their application remains contentious (*e*.*g*., Graur and Martin 2004; Wheat and Wahlberg 2013; Wilf and Escapa 2014; Cracraft et al. 2015; Mitchell et al. 2015; Wang and Mao 2015). Much of this controversy is due to the difficulties inherent in accurately associating data from the fossil record with particular nodes in a phylogenetic tree, as is required by the dominant method of divergence-time estimation, the “node-dating” approach. In a node-dating analysis, the investigator associates fossils (or other sources of temporal information, such as island ages, or age estimates from prior studies) with particular nodes in a phylogenetic tree, and provides a calibration density for each node that reflects the investigator’s belief about the temporal relationship between the calibrated node and its constraining fossil (Ho and Phillips 2009).

In a conceptual departure from node-dating approaches, the increasingly popular “tip-dating” or “total-evidence dating” (TED) methods treat fossils as their own terminals in the phylogenetic analysis and jointly infer the placement of the fossils, the patterns of morphological evolution, and a time-calibrated phylogeny. These methods use a dataset containing molecular characters for extant taxa and morphological characters for both extant and fossil taxa (Lee et al. 2009; Pyron 2011; Ronquist et al. 2012; Sterli et al. 2013). TED approaches thus overcome the main weaknesses of node-based methods: the phylogenetic affinities of calibrating fossils are inferred from the morphological data rather than depending on researchers’ implicit assumptions or separate cladistic analyses; the temporal connection between a fossil and a portion of the tree of extant species is not determined in advance; and many more fossils can be used, including fragmentary ones and ones that are members of wholly extinct clades. More generally, these approaches allow for the incorporation of a much greater proportion of the fossil record and shift divergence-time estimation toward a less subjective treatment of fossil data.

For these reasons, TED methods offer great promise for more transparent inferences of divergence time. However, inferring the topological and temporal position of fossils in a Bayesian framework requires the investigator to specify a model of discrete morphological evolution and a tree model that includes non-contemporaneous tips. The morphological model most frequently used— the Mk model (Lewis 2001)—assumes that rates of change between character states are the same for all characters, and has been criticized as inadequate for modeling morphological evolution (e.g., Sterli et al. 2013; Goloboff et al. 2019). Among the tree models, the fossilized birth-death models (Heath et al. 2014; Zhang et al. 2016; Gavryushkina et al. 2017) are an advance over the earlier uniform tree model (Ronquist et al. 2012) in the sense that they provide a coherent mechanistic description of diversification and preservation (Marshall 2019), but they are still limited in biologically important ways, and the relative impacts of these tree models on inference under TED have not been exhaustively explored.

In addition to the modeling concerns specific to total-evidence dating, TED methods share an unusual statistical pathology with other divergence-time estimation methods: nonidentifiability (dos Reis and Yang 2012; Zhu et al. 2015). Under the standard models of character evolution (continuous-time Markov chains), there is no information in molecular or morphological data about either rate or time individually; it is only their product (*e*.*g*., number of substitutions per site) that can be estimated. There are thus an infinite number of combinations of rate and time that have identical likelihoods for a given dataset, and the goal of divergence-time estimation—to isolate time from rate—depends on the priors on node ages and clock rates when performed in a Bayesian framework. This fundamental nonidentifiability of relaxed-clock models is apparent in node-dating analyses (Zhu et al. 2015), but is shared by TED analyses, too: the same data (character alignments) can be equivalently fit by very different combinations of model parameters, and those different model parameters could potentially lead to very different branch length inferences. There is thus a need for rigorous analyses of the behaviour of TED methods, for biologically meaningful priors, and for the development of general tools for evaluating model performance (Wilf and Escapa 2016).

Here we explore the sensitivity of TED analyses to these modeling choices, focusing on marattialean ferns (Marattiales: Polypodiopsida). These ferns have a deep fossil record that extends back more than 320 million years (see extensive review by Rothwell et al. 2018a); their relatively modest extant diversity (approximately 110 species; Murdock 2008a; Schuettpelz et al. 2016) belies their former dominance, especially during the Pennsylvanian when they were canopy dominants in both clastic and peat swamp communities (Cleal 2015; DiMichele and Phillips 1996, 2002; Phillips et al. 1985). Apparent diversity then declined in the Triassic through Cretaceous, and no unequivocal records exist from the Cenozoic (Lundgren et al. 2019; Rothwell et al. 2018a). This temporal pattern—clusters of extinct and extant diversity separated by a depauperate intermediate sample—as well as extensive morphological homoplasy among the extant genera (Murdock 2008b; Lehtonen et al. 2020) not only make traditional node-dating approaches effectively impossible, but may also challenge many of the assumptions of total-evidence dating models. In this study, we employ the statistical tools of sensitivity analysis, model adequacy, and model comparison to evaluate the impact of modeling choices, to improve estimates of marattialean phylogeny and divergence times, and to learn about the processes driving marattialean evolution and diversification.

## Marattiales

Marattialean ferns constitute one of the eusporangiate fern clades, and while their relationship to other ferns has historically been unclear (e.g., Schuettpelz et al. 2006; Qiu et al. 2007; Lehtonen 2011), an emerging consensus places them as the sister group to the largest group of extant ferns, the leptosporangiates (Rai and Graham 2010; Kuo et al. 2011; Grewe et al. 2013; Knie et al. 2015; Rothfels et al. 2015b; Kuo et al. 2018; Qi et al. 2018; Lehtonen et al. 2020; but see Wickett et al. 2014; Shen et al. 2018; One Thousand Plant Transcriptomes Initiative 2019). The approximately 110 extant species of Marattiales are divided among six genera (Murdock 2008a; Schuettpelz et al. 2016). These ferns are homosporous, tropical in distribution, and range in size from megaherbs (species of “king fern” in the genus *Angiopteris* can have leaves exceeding 3m in length) to the small dimorphic-leaved species of *Danaea* (Fig. S1).

Fossil taxa of the Marattiales are generally attributed to one of two families: the extinct Psaroniaceae and the still living Marattiaceae (see Rothwell et al. 2018a). The Marattiales fossil record includes spores, compressions, and impressions, but they are best known from Carboniferous taxa that were carbonate permineralized and described in anatomical detail, including whole-plant reconstructions (Fig S1E, M) The extensive record of the Marattiales, in its richness, extensiveness, and temporal extent (Millay 1997; Liu et al. 2000; Rothwell et al. 2018a), makes this lineage an ideal test case for divergence-time estimation.

In addition, the morphology of marattialeans is comparatively well-studied (e.g., Stidd 1974; Millay 1979; Millay and Taylor 1984; Hill and Camus 1986; Millay 1997; Liu et al. 2000; Murdock 2008b; Cleal 2015; Rothwell et al. 2018b), the phylogeny of the extant lineages is fairly well understood (Li and Lu 2007; Murdock 2008b; Senterre et al. 2014; Lehtonen et al. 2020), and there are pre-existing matrices of molecular and morphological (including strong fossil representation) characters available for a broad taxon sample Hill and Camus 1986; Murdock 2008b; Rothwell et al. 2018b; see also Lehtonen et al. 2020, which was published after our analyses were completed).

## Data

### Taxon samples

Our taxon sample includes fossil and extant members of the Marattiales and their sister clade, the leptosporangiate ferns (Fig S1; summarized in Table S.2). The ingroup sample comprises 45 fossil taxa representing either the Psaroniaceae or Marattiaceae, along with 26 extant Marattiaceae; the outgroup samples include nine extant species selected for phylogenetic breadth and six well-understood fossil reconstructions spanning the diversity of leptosporangiate ferns. We chose from among fossil taxa coded by Rothwell et al. (2018b), excluding taxa with large amounts of missing data that were primarily of interest for informing *Scolecopteris* classification. Additionally, we added three Marattiales species to expand our fossil age representation: *Floratheca apokalyptica* (early Permian), *Rothwellopteris pecopteroides* (late Permian), and an un-named species from the Lower Cretaceous assigned to the Marattiaceae by Vera and Césari (2016). Our extant-taxon sample is also based on Rothwell et al. (2018b) but altered to maximize the number of taxa that had both morphological and DNA data by adding five species of *Ptisana* and three species of *Danaea*.

In some of the analyses that follow, we used three additional plant fossils: 1) *Psilophyton crenulatum* (early Devonian; Doran 1980); 2) *Pertica quadrifaria* (early Devonian; Kasper and Andrews 1972), and; 3) *Rhacophyton ceratangium* (late Devonian; Cornet et al. 1976; Dittrich et al. 1983; see Supplemental Table S.2). We chose these taxa because they are among the most complete fossil vascular plants and are outside of the Marattiales + leptosporangiate clade.

### Morphological and molecular data

Our morphological dataset was largely derived from Rothwell et al. (2018b), which itself relied heavily on Hill and Camus (1986) and Murdock (2008b), and amended as necessary. Our final morphological matrix comprised 98 discrete characters describing anatomy and gross morphology; in total, there were 79 binary characters, 10 three-state characters, four four-state characters, three five-state characters, one six-state character, and one seven-state character. We provide the details of how we assembled and scored our morphological dataset in the Supplemental Material (S§1.1).

We used available chloroplast DNA sequences from Murdock (2008b) augmented with additional data for our outgroup taxa. The final dataset comprised 33 species with sequences from four chloroplast markers: *aptB, rbcL, rps4* + *rps4-trnS* spacer, *trnS-trnG* spacer + *trnG* intron (Table S.1).

The complete morphological and molecular matrices are available in the Data Dryad repository DOI:X and the GitHub repository https://github.com/mikeryanmay/marattiales_supplemental/releases/tag/1.0.

## Methods

### Models

The Bayesian total-evidence dating model consists of five main components: 1) the molecular substitution model; 2) the molecular clock model; 3) the morphological transition model; 4) the morphological clock model, and; 5) the tree model. There is a long history of studying the impact of substitution and molecular-clock models on phylogenetic inference (Huelsenbeck and Rannala 2004; Schenk and Hufford 2010; dos Reis and Yang 2012; Zhu et al. 2015). By contrast, the influence of the morphological-transition, morphological-clock, and tree models have received less attention, and earlier studies suggest that these model components can have strong effects on divergence-time estimates (Rothfels and Schuettpelz 2014; Condamine et al. 2015).

We performed analyses under a range of morphological-transition, morphological clock, and tree models to explore the relative impact of these three model components on phylogenetic estimates of the marattialean ferns. In total, these analyses comprised 72 model combinations: four morphological-transition models *×* two morphological-clock models *×* nine tree models, as described below (a graphical-model schematic is shown in Fig. 1; graphical model representations of specific model component are available in Supplemental Material S§2).

**Figure 1:**
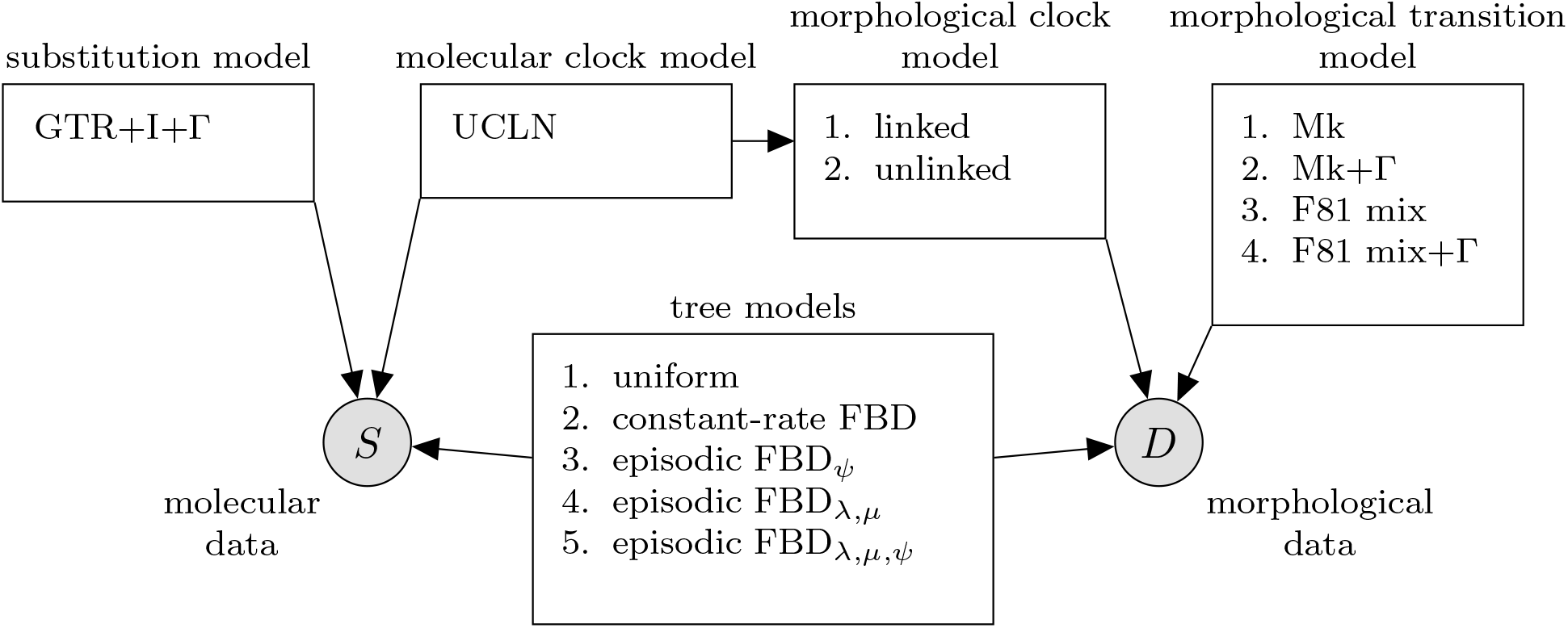
A graphical model representation of the total-evidence dating model. The total-evidence model includes five separate model components: 1) the substitution model; 2) the molecular-clock model; 3) the morphological-transition model; 4) the morphological-clock model, and; 5) the tree model. Each box shows the dependence between each module and the molecular (*S*) and morphological (*D*) datasets. GTR+I+Γ: General time-reversible substitution model with a proportion of invariant sites and gamma-distributed among-site rate variation. UCLN: Uncorrelated lognormal relaxed molecular clock. Mk: Markov transition model with *k* states. F81 mix: Markov transition model with a mixture of unequal stationary frequencies. FBD: Fossilized birth-death model.

#### Substitution model and molecular clock

In all analyses, we partitioned the molecular dataset by locus, and then by intronic and exonic regions, and for exonic regions, among codon positions. We assigned an independent GTR+I+Γ substitution model with four rate categories to each data subset. For the molecular clock, we assumed that branch-specific rates of molecular evolution were uncorrelated and drawn from a shared lognormal prior distribution (the UCLN model; Drummond et al. 2006).

#### Morphological transition models

As with substitution models for molecular evolution (*e*.*g*., GTR+I+Γ), the morphological transition model is composed of a model that describes how characters are partitioned, a model that describes how rates vary among characters within a partition (the among-character rate variation, or “ACRV”, model), and a model that describes the relative rates of change among character states (the morphological matrix model) within a partition.

For the morphological partition scheme, we partitioned characters based on the number of states—such that there was one subset for binary characters, one for three-state characters, etc.—for all analyses. Within each subset, we assumed the characters evolved according to one of the ACRV and morphological matrix models described below; for a given combination of ACRV and matrix model, we assumed each subset evolved under the same type of model, but with parameter values that differed among subsets.

We used two different morphological matrix models: an Mk model (Lewis 2001) that assumes that rates of change—and therefore stationary frequencies—are equal among states, and a model that allows the stationary frequencies to vary among character states; the latter model is equivalent to the F81 model commonly applied to molecular data (Felsenstein 1981), and we therefore refer to this model as an F81 model. The Mk model assumes that relative rates of transition are the same among character states, such that the stationary frequency of each state is the same among states *and* among characters; for example, for binary characters, state “0” in one character has the same frequency as state “0” in another character. Because the relative rates are the same among characters, and the overall rate of evolution is described by the morphological clock model (described below), the Mk model has no free parameters. For the F81 model, we further relaxed the assumption that stationary frequencies are shared across characters using a mixture model (Wright et al. 2016); we refer to this as the F81 mixture model. In this model, state “0” in one character is permitted to have a different stationary frequency than state “0” in another character. This feature is particularly important in morphological data because, unlike molecular data where, for example, a “T” indicates particular features regardless of which alignment site it occurs in, the naming of morphological character states is arbitrary, with no commonalities across characters. For binary characters, we discretized a Beta distribution into five mixture categories and defined an F81 transition matrix using the value of each category as *π*_0_. We assumed a symmetrical Beta distribution with shape parameter *γ*_*m*_; the symmetry of the distribution guarantees that the likelihood does not depend on the labeling of the binary states (*i*.*e*., which state is labeled “0” or “1”), and the five mixture categories guarantees that the middle category corresponds to a symmetric (Mk) model. For each multistate subset we drew five sets of stationary frequencies from a symmetrical Dirichlet distribution with parameter *γ*_*m*_, and also a vector of mixture weights, *ω*, which define the prior probability that a character evolves according to each of the stationary frequencies (Pagel and Meade 2004). We assumed the parameter *γ*_*m*_ was shared among the Beta distribution for the binary characters and the Dirichlet distributions for the multistate characters, and estimated *γ*_*m*_ and *ω* (one per number of states greater than two) from the data (Figure S6). For both models we corrected for the fact that only variable characters were included (Lewis 2001).

In addition to the two morphological transition models, we also modeled how rates of evolution varied among characters within each subset. We used two alternative among-character rate-variation models: a shared-rate model and a variable-rate model. The shared-rate model assumed that the rate of evolution is the same for all characters with the same number of states. For the variable-rate model, we assumed character-specific rates for characters with *i* states drawn from a discretized Gamma distribution with four categories and parameter *α*_*i*_, which we also estimated from the data.

In total, we used four different morphological transition models: Mk, Mk+Γ, F81 mixture, and F81 mixture+Γ.

#### Morphological clock models

The morphological clock model describes how rates of evolution vary among branches in the tree. We used two variants of this model: a linked model where the rate of morphological evolution on a given branch is proportional to the rate of molecular evolution on that branch, and an unlinked model where the morphological and molecular rates are independent. For the linked model, we included a single free parameter, *β*_*m*_, that defines the relative rate of morphological to molecular evolution across all of the branches (Figure S8). For the unlinked model, the free parameter *β*_*m*_ describes the relative *mean* rate of morphological to molecular evolution. We then drew each branch-specific rate, *r*_*m,i*_, independently from a lognormal distribution with standard deviation *σ*_*m*_, which we also estimated from the data (Figure S9).

#### Tree models

The tree model defines the probability of the tree topology and node ages. We used five different tree models: 1) a uniform model (Ronquist et al. 2012); 2) a constant-rate fossilized birth-death model (CRFBD; Heath et al. 2014); 3) an episodic fossilized birth-death model where fossilization rates were allowed to vary over time (EFBD_*ψ*_); 4) an episodic fossilized birth-death model where diversification rates (speciation and extinction rates, *λ* and *µ*) were allowed to vary over time (EFBD_*λ,µ*_), and; 5) an episodic fossilized birth-death model where all rates were allowed to vary over time (EFBD_*λ,µ,ψ*_). For the variable-rate episodic models, we divided time into 33 geological epochs (beginning with the Terreneuvian and ending with the Holocene) defined per the International Chronostratigraphic Chart (updated from Cohen et al. 2013), and allowed one or more parameters to vary among epochs according to a mixture model. Specifically, we assumed that there were three mixture categories, each with their own rate parameter and a mixture weight *κ*_*i*_. The assignments of epochs among mixture categories were treated as independent random variables (with values 1 through 3) such that each epoch was assigned to one of the three mixture categories with prior probability *κ*_*i*_. For a given assignment, each epoch was associated with a specific rate parameter; we then estimated the rate and mixture weight for each category, and the assignment of each epoch among categories. By averaging over the assignment of epochs to each mixture category, this model also provides an estimate of the epoch-specific parameters. For models where multiple rates varied, we assumed that they varied independently (*i*.*e*., that they were drawn from separate mixture models). This model differs from previous work where each time slice was allowed to have an independent rate parameter (rather than being drawn from a mixture model), but used a small number of time slices (see Zhang et al. 2016; Wright et al. 2020). Our approach balances model complexity (the number of rate parameters) and the temporal resolution over which rates vary (the number of time slices). For all models, we specified a uniform prior distribution between 410 and 550 Ma on the age of the origin of the tree. We conservatively based the maximum age on the age of the earliest plant fossil evidence (cryptospores from the Ordovician ∼470 Ma Rubinstein et al. 2010), and the minimum age on the age of the oldest sample in our extended “ancient plants” dataset (see below). Additionally, we accommodated uncertainty in the ages of each fossil, which has been demonstrated to be important for accurate divergence-time estimates (Barido-Sottani et al. 2019).

The final component of the FBD tree models is the mechanism for accounting for incomplete sampling of extant taxa. Our taxon sample includes a relatively well-sampled extant ingroup (27 of 111 extant taxa), and a very sparsely sampled outgroup (nine of ∼ 12000; Schuettpelz et al. 2016). To accommodate this heterogeneous taxon sampling we therefore conducted each fossilized birth-death analysis twice, once with a sampling fraction, *ρ*, corresponding to the ingroup portion of the tree (*ρ* = 27 ÷ 111), and once corresponding to the full tree (*ρ* = 36 ÷ 12000).

In total, we used nine different tree models: the uniform tree model, plus four fossilized birth-death models *×* two taxon-sampling fractions.

### Analyses

#### MCMC analyses

For each combination of the above model components, we estimated the posterior distribution using four replicate Metropolis-coupled MCMC with five coupled chains in RevBayes (Höhna et al. 2016). We performed all of our analyses on the University of California, Berkeley HPC cluster, savio (run times and additional computation details are described in Supple-mental Material S 3.2). We assessed whether each chain failed to converge to, or sample adequately from, the joint posterior distribution using protocols described in the Supplemental Material (S§3.1). All of the scripts used for analysis and post-processing are available in the Data Dryad repository DOI:X and the GitHub repository https://github.com/mikeryanmay/marattiales_supplemental/releases/tag/1.0.

#### Comparing estimates of topology and branch lengths

We employed two techniques to summarize differences in phylogenetic estimates under the model combinations that we explored. First, we used multidimensional scaling (MDS) of tree-distance metrics (following Hillis et al. 2005) to compare the distributions of tree topologies and branch lengths inferred under these models. MDS projects the pairwise distance between each tree in a sample of trees into a lower dimensional—and therefore easier to visualize—representation of tree space. To facilitate automated MDS for a large number of comparisons, we implemented these MDS analyses using the R packages phangorn and smacof (de Leeuw and Mair 2009; Schliep et al. 2017; R Core Team 2019). We compared the distribution of phylogenies using MDS with three different metrics: 1) the Robinson-Foulds distance (RF, a measure of topological distance; Robinson and Foulds 1981); 2) the Kü hner-Felsenstein distance (KF, a distance metric that incorporates both topology and branch lengths; Kü hner and Felsenstein 1994) between time-scaled phylogenies (chronograms), and; 3) the KF distance between phylogenies with branch lengths proportional to the expected amount of morphological evolution (“morphograms”). Computing distances among phylogenies with sampled ancestors is difficult, since the number of tips and branches depends on the inferred number of sampled ancestors. We therefore resolved all sampled ancestors onto zero-length branches before computing distances among trees. We included 100 trees from the posterior distribution of each of the models in our MDS plots, and colored each point in the resulting tree space according to one of the TED model components to visually compare the relative impact of each model component on the posterior distribution of trees.

Second, we used lineage-through-time (LTT) curves to summarize divergence-time estimates under each model. An LTT curve displays the number of branches in the inferred tree at any given time, and therefore provides a less abstract summary of divergence-time estimates than MDS; however, in contrast to MDS plots, the LTT approach is unable to capture differences in topology. We compute the average LTT curve under a given model by calculating the average number of branches present at each time point for each tree in the posterior distribution; similarly, we compute the 95% credible interval (CI) of the number of branches at a given time. To compare the relative impact of the three model components, we compute the average LTT curve for each model, then compute the average of these curves among all model combinations that share the focal model component.

#### Stochastic character mapping

We used stochastic character mapping (Huelsenbeck et al. 2003) to visualize patterns of morphological character evolution across the phylogeny. In these analyses, we conditioned on the maximum-clade-credibility (MCC) tree inferred for each model, as well as posterior mean estimates of all relevant model parameters, and simulated stochastic maps using the R package phytools (Revell 2012).

#### Implied total diversity over time

We simulated lineages under each of the fossilized birth-death models, without fossilization, to generate the implied total number of lineages present at each point in time. For each model, we sampled the origin time and diversification parameters from the corresponding posterior distribution, then simulated lineages forward in time until the present to generate the posterior-predictive distribution of the full diversification process. We computed the median and 95% CI of the number of lineages at a large number of evenly spaced time points to summarize the distribution of the implied total diversity over time. These simulations do not condition on the sampled tree, and therefore should not be interpreted as the posterior distribution of the number of missing taxa in our inferred tree. Rather, these simulations represent the distribution of the number of lineages if we were to repeat the inferred lineage-diversification process many times.

#### Assessing absolute model adequacy

We assessed the adequacy of each model using posterior-predictive simulation (PPS; Bollback 2002; Höhna et al. 2018). The premise of PPS is that, if a given model provides an adequate description of the true process that generated the observed data, then datasets simulated by the model should resemble the observed data. The degree of resemblance for a given simulated dataset is described by a summary statistic that is designed to capture relevant aspects of the data-generating process. If the distribution of this statistic computed across simulated datasets (the posterior-predictive distribution, PPD) contains the statistic for the observed data with high probability, then the model is deemed adequate, *i*.*e*., the model provides a reasonable description of the true data-generating process.

We assessed the adequacy of our models from the perspective of the morphological data. For a given model, we drew random samples from the posterior distribution of model parameters (including the phylogeny and parameters of morphological evolution). For each sample, we simulated a morphological dataset, the same size as the original and with the same patterns of missing data, given the model parameters and computed two summary statistics: 1) *S*, the total parsimony score (number of steps computed on the sampled tree) for the simulated characters minus the total parsimony score for the observed characters, and; 2) *V*, the variance in parsimony scores among simulated characters minus the variance in parsimony scores among observed characters. The *S* statistic is intended to assess whether the model adequately characterized the average rate of evolution, while the *V* statistic is intended to assess if the model captures how the rate and process of evolution vary among characters. If the 95% interval of the posterior-predictive distribution of these statistics for a given model did not include 0, we deemed that model inadequate. We provide further details of these simulations in the Supplemental Material (S§4).

The structure of the TED model provides some expectations about the sensitivity of PPS to the different model components we explored. Specifically, the PPD will be sensitive to model components that have a strong impact on the likelihood function, but insensitive to non-identifiable model components. We therefore expect the morphological-transition model to influence the PPD because this model component is identifiable. By contrast, for a given morphological-clock model, the tree model may have little influence on the PPD because rate and time are nonidentifiable: tree models that prefer different node ages may nonetheless have similar likelihoods because the clock model can compensate for different node ages. However, for a given tree model, the two clock models we used can in principle influence the PPD because the models with separate relaxed clocks can achieve sets of branch rates that models with one relaxed clock cannot (Zhu et al. 2015). For example, if the unlinked model is true (*i*.*e*., if rates of morphological and molecular evolution are not proportional), then estimates of branch-specific rates of morphological evolution under the linked model will be influenced by the molecular data and therefore be unable to attain values that fit the morphological data.

#### Assessing relative model fit

We compared the relative fit of competing models using Bayes factors, which are the ratio of the marginal likelihood of each model (Kass and Raftery 1995). We estimated the marginal likelihood for a given model using four replicate power-posterior analyses in RevBayes, and computing the path-sampling (Lartillot and Philippe 2006) and stepping-stone estimators (Xie et al. 2011). Given the computational expense of marginal-likelihood estimation, we only calculated Bayes factors among the tree models (conditional on the preferred morphological-transition and morphological-clock models).

#### Assessing the effect of the taxon sample and rooting

Our fossil dataset is dominated by a cluster of late Carboniferous taxa whose relationships to each other and to surviving lineages are apt to be highly uncertain, which may limit our ability to infer ancient divergences. Additionally, the inclusion of a sparsely sampled out-group makes it difficult to specify an appropriate taxon-sampling fraction for the fossilized birth-death models. We performed additional analyses to understand the robustness of divergence-time estimates within the Marattiales to different taxon samples and rooting strategies (described in more detail in the Supplemental Material S§5). In particular, in addition to our “standard” taxon set (the ingroup Marattiales plus an outgroup of leptosporangiate ferns) we performed analyses with only ingroup taxa, with the addition of the ancient land-plant fossils described above, and by polarizing a subset of characters for which we are confident in their ancestral states *a priori* (our “ingroup-only”, “ancient plants”, and “polarized” analyses, respectively). In each of these analyses, we assumed the best-performing morphological transition model, morphological clock model, and tree model, as determined in the core analyses.

## Results

### Modeling results

Here, for the fossilized birth-death models, we present the results for the analyses that assume the ingroup sampling fraction, for a total of 40 distinct model combinations (four morphological transition models × two morphological clock models × five tree models); results using the overall sampling fraction are qualitatively similar, and are presented in the Supplemental Material S§6.2. LTT curves for each individual model combination are available in the Supplemental Material. We also compared the ages of individual clades for each pair of models (Figs. S20, S21, S22), and between ingroup and overall sampling fractions (Fig. S59).

### Topological distance

The greatest differentiation in topological space is between models with and without the uniform tree model (Fig. 2, top row); the individual fossilized birth-death tree models each have weak effects on topology (Fig. 2, top right). The morphological transition model has a strong influence on topologies (Fig. 2, top left), whereas the morphological clock models have a mild effect (Fig. 2, top middle).

**Figure 2:**
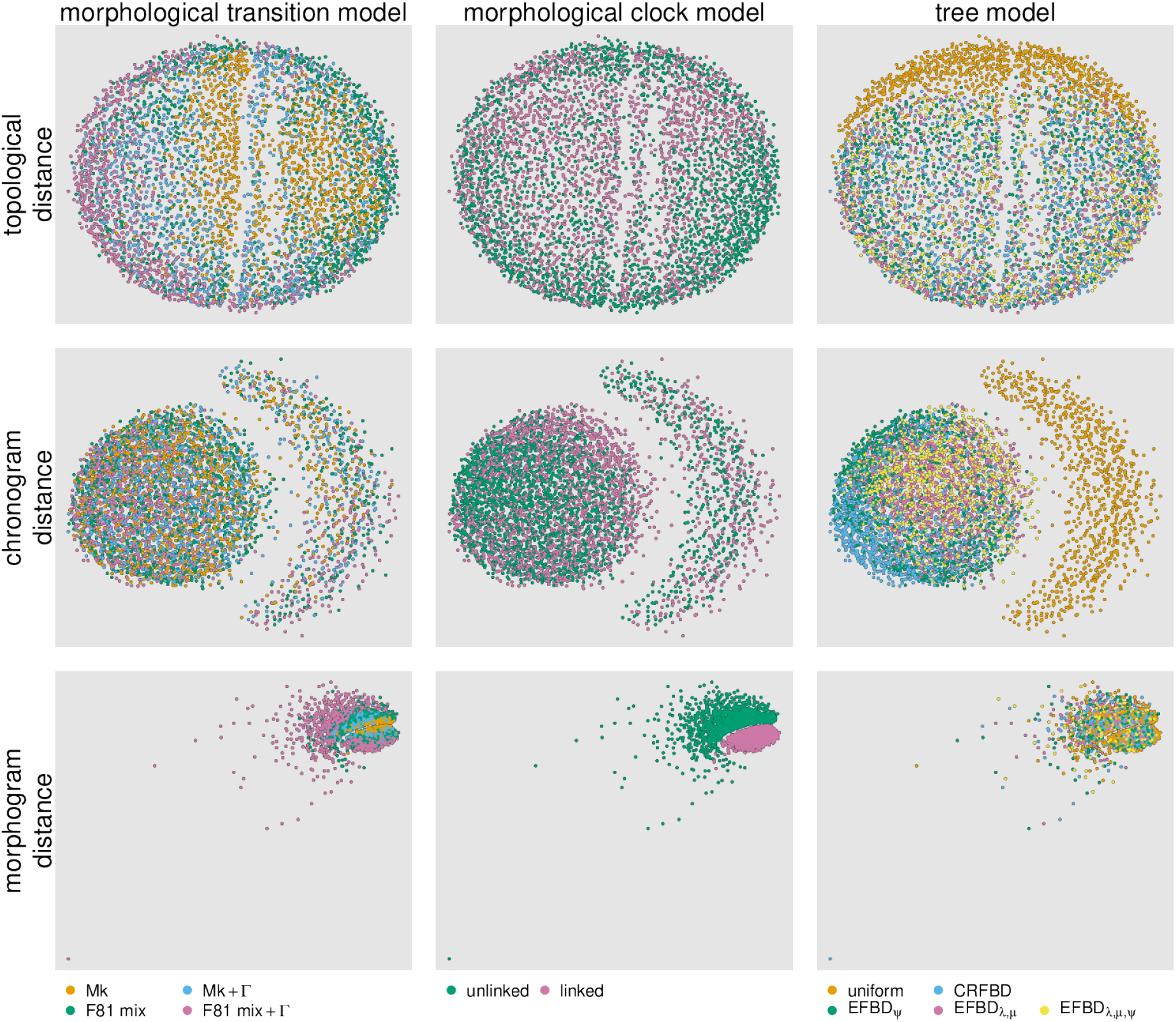
Comparing distributions of trees among model combinations. We compute the Robinson-Foulds distance (RF, a measure of topological distance, top row), the Kühner-Felsenstein distance (KF, a distance metric that incorporates both topology and branch lengths) between chronograms (middle row) and morphological phylograms (“morphograms”, bottom row). We then plot the (square-root transformed) distances in two-dimensional space using multi-dimensional scaling (MDS); each point represents the location of a given sampled tree in tree space according to the distance metric. We color points according to the morphological transition model (column 1), the morphological clock model (column 2), or the tree model (column 3). These results assume the ingroup sampling fraction for all of the fossilized birth-death models; see Figure S30 for the results with the overall sampling fraction.

### Chronogram distance

Similar to the pattern observed in topological distances, the uniform tree model has the most striking impact on Kü hner-Felsenstein distances among chronograms (Fig. 2, middle row). However, in contrast to topological distances, KF distances among chronograms are differentiated by the FBD models, with the EFBD_*λ,µ*_ and EFBD_*λ,µ,ψ*_ models sampling from a region of tree space (Fig. 2, middle right, central region) that is rarely visited by the other models. The morphological transition and clock models appear to have a mild effect on KF distances (Fig. 2, middle row, left and middle columns).

### Morphogram distance

In contrast to its weak effect on topological and chronogram distances, the morphological clock model has the strongest impact on KF distances among morphograms (Fig. 2, bottom middle). Within the clusters defined by the clock models, there is clear differentiation among transition models, with the Mk+Γ and F81 mixture models being intermediate between the Mk and F81 mixture+Γ models (Fig. 2, bottom left). The tree models have a mild affect on distances among chronograms (Fig. 2, bottom right).

### Lineages through time

Consistent with our MDS plots, the greatest differences among LTT curves are between model combinations with and without the uniform tree model; however, there are also consistent differences in LTT curves among the fossilized birth-death models (Fig. 3, right). In particular, the EFBD_*λ,µ*_ and EFBD_*λ,µ,ψ*_ models indicate a later origin for the Marattiales and a more rapid increase to peak diversity at the end of the Carboniferous, whereas the EFBD_*ψ*_ model predicts a more gradual accumulation of diversity. Overall, the EFBD_*λ,µ*_ and EFBD_*λ,µ,ψ*_ models estimate very similar LTT curves, and the influence of the tree model on LTT curves decreases toward the present.

**Figure 3:**
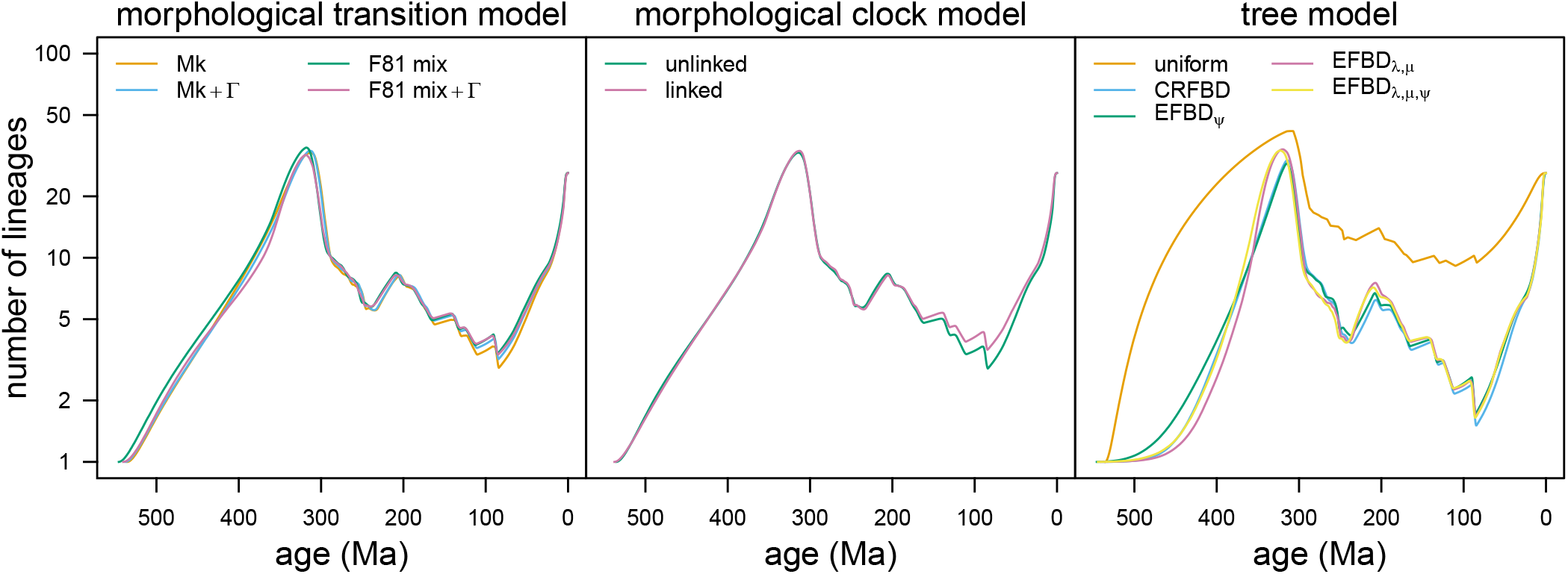
Comparing lineage-through-time curves among model combinations. For each focal model component (morphological transition model, and morphological clock model, and tree model, respectively), we compute the LTT curve averaged over the remaining model components, *i*.*e*., the average number of branches in the phylogeny at a given time, averaged over all model combinations that share the focal model component. We removed outgroup taxa to emphasize the influence of model specification on age estimates within our ingroup. Left) We compute the average LTT for each of the 40 models (from 2000 sampled trees for each model), then compute the mean of the resulting average LTT among models with the same morphological-transition model. Middle) As in left, but we compute the mean of the average LTT among models with the same morphological clock model. Right) As in left, but we compute the mean of the average LTT among models with the same tree model.

The morphological transition model has a mild but consistent impact on lineage-accumulation leading up to the Carboniferous (Fig. 3, left). Following the Carboniferous, the transition models generally result in very similar LTT curves, with the exception of the Mk model, which infers generally fewer lineages through the Cenozoic.

Whether rates of morphological evolution are linked or unlinked to rates of molecular evolution has almost no impact on LTT curves in the early history of the Marattiales. However, LTT curves for the clock models begin to diverge beginning about 175 Ma, after which the unlinked model infers younger clade ages (Fig. 3, middle).

### Absolute model fit

The morphological transition model had a strong impact on model adequacy (Fig. S23, colored boxplots). In particular, transition models without among-character rate variation did the worst according to the parsimony variance statistic, *V*; additionally, the F81 mixture+Γ model did the best at describing the amount of evolution, according to the total parsimony score statistic, *S*. We therefore identified the F81 mixture+Γ as the preferred morphological transition model.

Posterior-predictive distributions were largely insensitive to the morphological clock model (Fig. S23, shaded regions), a result that is in strong contrast to the influence of the clock model on the posterior distributions of morphograms (Fig. 2, bottom middle). The strong effect of the clock models on the morphograms indicates that the absence of a signal in the PPS is not related to nonidentifiability. Because this model component had a modest effect on divergence-time estimates, we preferred the simpler, linked morphological clock model for further analyses.

As predicted based on nonidentifiability, the tree model had essentially no impact on posterior-predictive distributions (Fig. S23, columns), consistent with its very limited impact on posterior distributions of morphograms (Fig. 2, bottom right). We therefore compared the relative fit of the tree models using Bayes factors.

We also assessed absolute model fit with either the extinct or extant taxa pruned from the tree and morphological data before computing the summary statistics. When extinct taxa were removed, the simulated datasets consistently overpredicted the parsimony score (Fig. S24); by contrast, when extant taxa were removed, simulated datasets had lower parsimony scores than the observed morphological dataset (Fig. S25). The variance in the parsimony score across characters, *V*, was relatively insensitive to the exclusion of either extant or extinct taxa.

### Relative model fit

Bayes factor comparison of the tree models were decisive (Table 1). The uniform tree model is by far the worst model: it is very strongly outperformed by the second worst model, the constant-rate FBD model (2 ln BF *>* 40). Interestingly, models that allow fossilization rates to vary among epochs do not improve model fit: the EFBD_*ψ*_ is disfavored compared to the constant-rate FBD, and the EFBD_*λ,µ,ψ*_ model is disfavored compared to the EFBD_*λ,µ*_ model. Overall, the EFBD_*λ,µ*_ model is the best-performing model, and is very strongly favored over the second-best model, EFBD_*λ,µ,ψ*_ (2 ln BF = 14.25; Table 1).

**Table 1:**
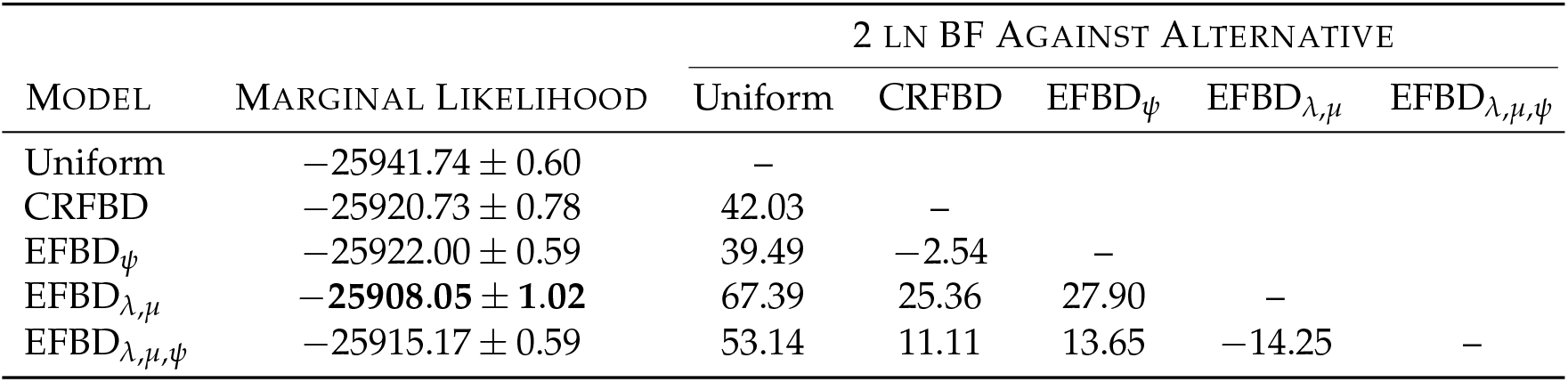
Comparing the fit of alternative tree models with Bayes factors. We report the average marginal likelihood *±* the standard deviation among four runs computed using the stepping-stone estimator, and the 2 ln Bayes factor between each pair of models. Marginal likelihoods computed using the path-sampling estimator were essentially identical.

| MODEL | MARGINAL LIKELIHOOD | 2 LN BF AGAINST ALTERNATIVE |  |  |  |  |
| --- | --- | --- | --- | --- | --- | --- |
|  |  | Uniform | CRFBD | EFBD<sub>ψ</sub> | EFBD<sub>$\lambda, \mu$</sub> | EFBD<sub>$\lambda, \mu, \psi$</sub> |
| Uniform | $-25941.74 \pm 0.60$ | – | | | | |
| CRFBD | $-25920.73 \pm 0.78$ | 42.03 | – | | | |
| EFBD<sub>ψ</sub> | $-25922.00 \pm 0.59$ | 39.49 | –2.54 | – | | |
| EFBD<sub>$\lambda, \mu$</sub> | **$-25908.05 \pm 1.02$** | 67.39 | 25.36 | 27.90 | – |  |
| EFBD<sub>$\lambda, \mu, \psi$</sub> | $-25915.17 \pm 0.59$ | 53.14 | 11.11 | 13.65 | –14.25 | – |

### Taxon sampling fractions

Results under fossilized birth-death models assuming the overall sampling fraction (*ρ* = 36 ÷ 12000; S§6.2) are qualitatively similar to those assuming the ingroup sampling fraction (*ρ* = 27 ÷ 111); in particular, clade-age estimates under the overall sampling fraction are modestly older than those under the ingroup sampling fraction (S§6.3).

### Taxon sample and rooting analyses

The broad topological patterns and timing of divergences are similar among these analyses (Figs. S§57, S58, S59). Nevertheless, there are some notable trends in the effect of the different approaches on inferred ages of specific clades, which we discuss in the Supplemental Material (S§5). Notably, the similarity between ages inferred from our primary analyses with the ingroup sampling fraction and those inferred under the ingroup-only analysis suggests that age estimates based on the ingroup sampling fraction are reliable. By contrast, those under the overall sampling fraction are probably overestimates (S 6.3). Because these analyses used different datasets, we can not correctly compare among them with posterior predictive simulation or Bayes factors. However, given that these analyses provided broadly consistent results, we ultimately base our discussion of marattialean phylogeny on the most inclusive dataset (*i*.*e*., the “ancient plants” analysis).

### Phylogenetic results

#### Topological results

Topologies varied among analyses, but all analyses except those under the uniform tree prior resolved a clade of extant Marattiaceae (exclusive of any fossil representatives) and a grade of fossil taxa diverging from the stem of this extant clade that includes a Psaroniaceae clade comprising many Carboniferous to Triassic (– Cretaceous) taxa (Figs. 4, S26, S43, S52, S55). In the MCC tree for our focal phylogeny (from the “ancient plants” analysis), *Scolecopteris* species do not fall within a single clade (Fig. 4). Instead, most *Scolecopteris* representatives, along with a few other traditional psaroniaceous fossil genera, fall in a “core” Psaroniaceae clade, while other *Scolecopteris* species along with *Grandeuryella renaulti* and *Floratheca apokalyptica* form a small grade of lineages at the base of the remaining Marattiales with which they share a reduced number of sporangia per synangium (Fig. S70) and spore ornamentation characteristics (Figs. S73, S74). The analyses with polarized-character rooting and the ingroup-only dataset produced similar results but with different arrangement of taxa in the grade (Figs. S51, S55).

**Figure 4:**
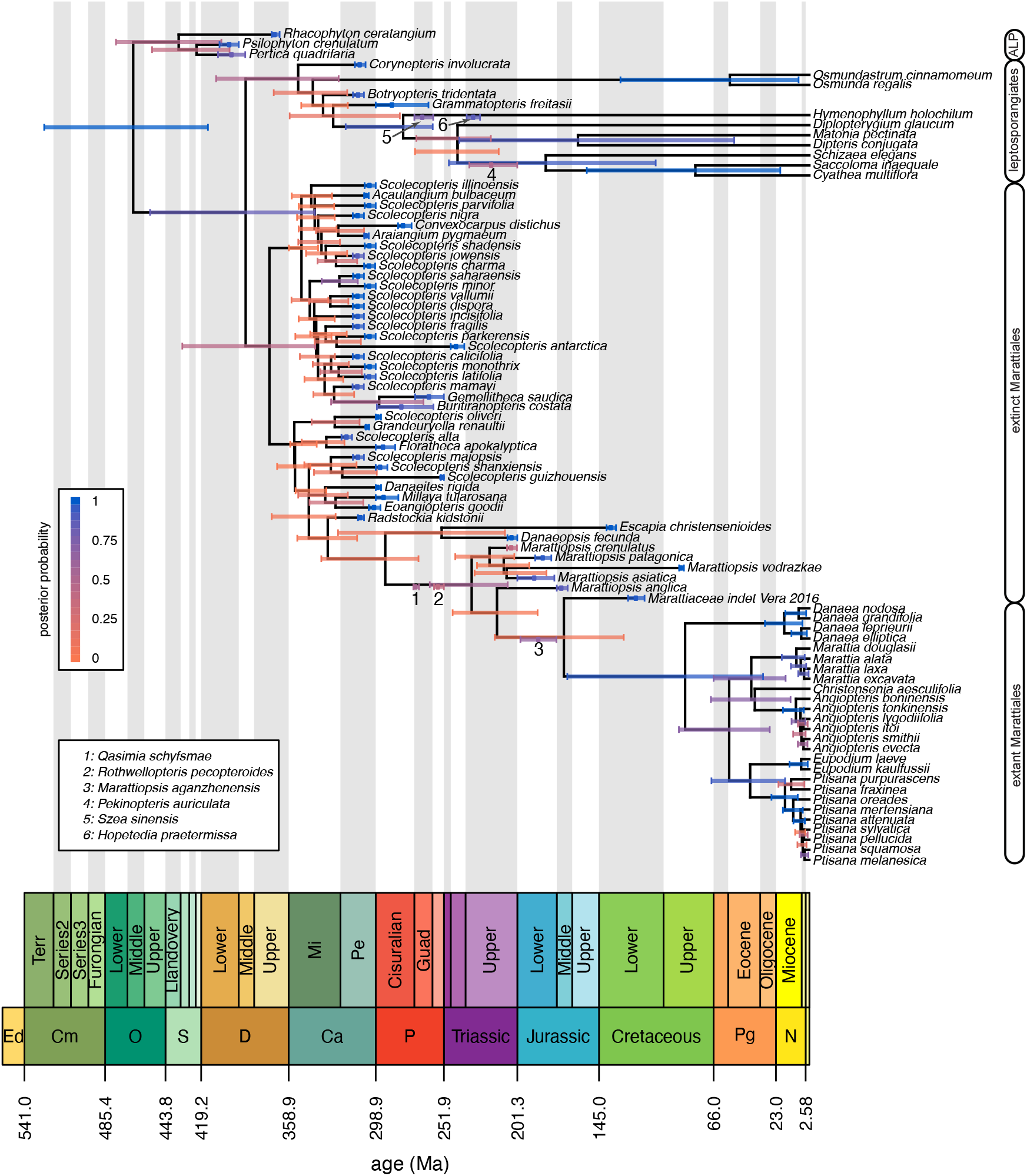
The maximum clade credibility tree under the preferred model with the “ancient plants” analysis. Bars correspond to the 95% credible interval of clade ages (for internal nodes) and tip ages (for fossils). Numbers and associated age bars along internal branches correspond to sampled ancestors (key bottom left). Bars are colored in proportion to the posterior probability of the clade for internal nodes, by the probability that the specimen is not a sampled ancestor for tips, and by the probability that the specimen is a sampled ancestor for sampled ancestors (legend, left). Major groups are labeled (right; ALP = ancient land plants). We plot time intervals according to the International Chronostratigraphic Chart (2020, updated from Cohen et al. 2013).

The “standard dataset” results weakly support an alternative hypothesis, in which there is an expanded Psaroniaceae clade that consists of all *Scolecopteris* species plus representatives of other genera, most of which have traditionally been considered to belong to Psaroniaceae (*e*.*g*., *Araiangium pygmaeum, Acaulangium bulbaceum, Grandeuryella renaulti, Convexocarpus distichus, Floratheca apokalyptica, Buritiranopteris costata*, and *Gemellitheca saudica*; Fig. S26). This topology does not appear to be supported by obvious character-state transitions. In each case, support for the topology is low, as it is in all fossil-rich regions of the tree (Figs. 4, S26, S51, S55). Support for the monophyly of the ingroup was greatest for the ancient and polarized analyses (*i*.*e*., the ones with additional information available to inform the root position; posterior probabilities 0.43 and 0.85, respectively, compared to 0.33 with the standard dataset; Figs. 4, S51). Conversely, the ingroup-only analysis was the most topologically distinct in this fossil-rich region of the tree (Figs. S57, S55), and in general has lower posterior support for divergences along the backbone of the tree.

The “early diverging” stem lineages related to *Marattiopsis* and extant Marattiaceae consistently include *Radstockia kidstonii, Qasimia shyfsmae*, and *Rothwellopteris pecopteroides*, as well as a clade comprising *Millaya tularosana, Eoangiopteris goodii*, and *Danaeites rigida* (Figs. 4, S26, S27, S51, S52, S54, S55, S56). *Escapia christensenioides* and *Danaeopsis fecunda* were highly unstable and inferred in various positions among these stem lineages (Figs. 4, S26, S55) or with *Scolecopteris* species (Fig. S51). Pruning these two taxa did not have an appreciable effect on clade support (not shown).

*Marattiopsis* fossils, from the Triassic through Jurassic, are morphologically very similar to extant marattialeans and, with “Marattiaceae indet. Vera (2016)”, form a grade of lineages most closely related to the extant taxa. *Qasimia shyfsmae* and/or *Rothwellopteris pecopteroides* are often inferred to be directly ancestral to the *Marattiopsis* + extant Marattiaceae clade. Notably, the phylogenetic separation of the extant Marattiaceae genera from the morphologically nearly indistinguishable *Marattiopsis* species (see Fig. S1) and the superficially similar *Daneopsis fecunda* only occurs when temporal data are incorporated: in a separate analysis under a non-dated tree model (*i*.*e*., without temporal data), the support for the monophyly for the extant clade dropped dramatically (to *<* 0.001 posterior probability, Fig. S76). Relationships within crown Marattiaceae are consistent among analyses (Figs. 4, S26, S43, S51, S55). *Danaea* is sister to the remaining genera and *Marattia* is sister to a weakly supported *Christensenia* + *Angiopteris* clade; they are in turn sister to *Eupodium* + *Ptisana*. Our leptosporangiate out-group taxa are themselves also monophyletic, although the position of fossils within the group varies among analyses (Figs. 4, S26, S43, S51), with the Permian *Szea sinensis* and in some cases the Triassic *Hopetedia praetermissa* inferred to be directly ancestral to extant taxa.

#### Divergence-time estimates

The overall temporal pattern we infer suggests that the Marattiales and leptosporangiate fern lineages diverged from each other in the Lower or Middle Devonian (posterior mean 414.84 Ma, 95% CI = [350.90 − 505.74]; Fig. S58, Table S.3).

Early-diverging stem lineages of Marattiales, mostly involving taxa traditionally assigned to Psaroniaceae, diverged through the earliest Permian. Fossil taxa attributed to *Marattiopsis* begin to diverge in the Triassic (Fig. 4, S51, S55), followed by a gap of ∼ 115 million years (Table S.5) before the inferred divergence of the crown Marattiaceae in the Upper Cretaceous, although there is considerable uncertainty around the age of this crown node (posterior mean 85.50 Ma, 95% CI = [37.44 − 173.74], Table S.4). While all extant genera are inferred to have diverged from each other by the Eocene, the extant species diverged from each other beginning in the Neogene (Figs. 4, S26, S51, S55).

## Discussion

Our results bear both on broad methodological issues, and on details of marattialean phylogeny and biology. We begin by discussing the consequences of model specification on total-evidence dating, as well as prospects for future model and method development. Next, we discuss the implications of our results in the context of existing work on the phylogeny of Marattiales. Finally, we reconcile our methodological results with prior knowledge about the fossil record and marattialean paleoecology to draw a synthetic inference about the evolutionary history of the group, and to demonstrate the harmony between our inferences—particularly under the fossilized birth-death model—and the fossil record.

### Model specification and total-evidence dating

The fundamental advance of total-evidence dating is that fossil specimens are included as extinct samples in the tree, and their topological and temporal relationships to extant lineages are inferred from the available data, rather than assumed *a priori* (or inferred in separate cladistic analyses) as in traditional node-based fossil calibration approaches. Estimating the phylogenetic position of fossils requires that we specify models that describe how morphological characters evolve, how rates of morphological evolution vary among lineages, and how lineages are distributed in time (Warnock and Wright 2020). As with any model-based method, the inferences we make will naturally be influenced by these modeling choices. However, this sensitivity is of special concern for divergence-time estimation analyses because some parameters in the model—namely, the node ages and clock rates—are nonidentifiable (Rannala 2002; Stadler and Yang 2013): the data cannot distinguish, for example, between a branch having a fast rate and a short duration, or a slow rate and a long duration. This pathology makes it difficult to use standard maximum-likelihood methods to estimate divergence times because the ML divergence-time estimates will not be unique (though see Sanderson 1997, 2002; Paradis 2013, for almost-Bayesian ML solutions to this problem). In a Bayesian framework the problem is less acute, but at the cost of posterior estimates of time and rate that are very prior sensitive, even with large datasets. Specifically, the clock and tree models are the priors on time and rate in the Bayesian total-evidence dating model, and it is therefore critical to understand their role in total-evidence dating analyses.

We discuss the influence of each model component— and the prospects for elaborating upon these models—in the following sections.

*The relative impact of model components: Which models matter?—*Among the three model components, the tree model had the greatest influence on divergence-time estimates (Fig. 2, row 2, column 3), consistent with the expected prior sensitivity of this portion of the model. In particular, chronograms inferred under the uniform model strongly differed from those under the other models, which we attribute, counter-intuitively, to the uniform prior being very informative (as we discuss in the next section). However, distributions of chronograms were markedly different even among the fossilized birth-death models (Fig. S17). Lineage-through-time curves indicate that the influence of these models was greatest in (but not restricted to) the older, fossil-rich parts of the tree (Fig. 3, right), presumably because there is less information available from the molecular data to constrain older branch lengths, particularly for extinct clades.

Consistent with previous simulation results (Klopfstein et al. 2019), the model of morphological evolution had a negligible impact on divergence-time estimates in our study (Fig. 2, row 2). This result held after pruning extant taxa (Fig. S16), suggesting that this phenomenon is not the result of the information in the molecular data overwhelming the information in the morphological data. However, the morphological transition model had an obvious impact on the posterior distribution of tree topologies (Fig. 2, row 1 column 1), especially for fossil taxa (Fig. S16). The discrepancy between the impact of the transition model on chronograms (negligible) and topology (considerable) may be due to the fact that our fossil dataset is dominated by a large number of closely related and similarly-aged taxa: there may be subtle (and weakly supported) differences about inferred relationships among these taxa that ultimately have little impact on clade ages. Similarly, the two morphological clock models we compared (rates linked to the molecular clock, or not) had a profound influence on inferred *morphograms* (Fig. 2, row 3 column 2) but only a minor impact on chronograms (Fig. 2, row 2, column 2). As expected, the linked model had the largest consequences on the inferred ages of young clades (Fig. 3, middle), where the influence of the molecular data should be strongest.

The contrasting influence of the tree model and morphological-clock model on old and young clades makes sense in light of what we know about divergence-time estimation under relaxed-clock models. Specifically, divergence-time estimates are a compromise between the plausibility of the node ages (from the perspective of the tree model) and the plausibility of the branch rates (from the perspective of the relaxed-clock model) that are needed to explain the effective branch lengths (the product of rate on time of each branch). Where there is a lot of information about effective branch lengths (*e*.*g*., in clades with extant species and therefore abundant molecular data), the tree model and clock model must come to a compromise about how to explain the effective branch lengths, and the constraints of the clock model may reduce differences among the tree models. Where there is less information about effective branch lengths (*e*.*g*., in fossil clades), then the tree model will be less constrained by the clock model and will have a larger role in determining the branch lengths; in the extreme case when there are no character data, the tree model will completely determine the length of branches subtending fossils (as in Heath et al. 2014). However, the relative impact of these models in our analyses could be a consequence of the topological and temporal distribution of our fossil accessions, and should not be taken as a general pattern. We emphasize that we did not compare different (prior) forms of clock models, as the only difference in the morphological clock models that we examined was whether the morphological rates were linked to the molecular ones or not. As molecular clock models are known to have a strong potential influence on divergence-time estimation—both from first principles (the nonidentifiability of rate and time) and from empirical results (*e*.*g*., Ronquist et al. 2012; Rothfels and Schuettpelz 2014; Crisp et al. 2014; Zhang et al. 2016)— further studies are needed to examine the impact of these models, and their interaction with the tree models.

### The uniform tree model

A noninformative prior is one that maximizes the ability of the data to express its preference for different parameter values, a property often attributed to “uniform” priors. However, the informativeness of a particular prior can depend on what aspect of the model is being conceived; for example, a uniform prior over tree topologies is not uniform over clades (Pickett and Randle 2005). Technically, a noninformative prior expresses the same amount of prior evidence regardless of how we parameterize the model (Jaynes 1968); this is rarely (if ever) the case with uniform priors (Zwickl and Holder 2004, and references therein).

The uniform tree model (*sensu* Ronquist et al. *2012) is likewise informative (i*.*e*., not noninformative) about node ages (Warnock and Wright 2020); however, our results suggest that the strength of the uniform tree prior on node-age estimates has been underappreciated. Conceptually, this prior distribution on node ages is constructed as follows: for each tip, draw a uniform random variable between the age of the tip and the origin time (stem age) of the tree, order the resulting uniform random variables, then use the smallest (*i*.*e*., youngest) uniform random variable as the first node age (looking backward in time), the second smallest as the second node age, and so forth. Perhaps counter-intuitively, ordering the uniform random variables results in a prior distribution on node ages that becomes increasingly informative as the number of tips increases. To develop an intuition for why this is the case, we can imagine the procedure for a set of *n* contemporaneous (extant) tips with stem age of 1 arbitrary time unit, so that each (unordered) node age is a uniform random variable between 0 and 1. As the number of random variables increases, the variance around any given (ordered) node age shrinks (Figs. S46, S47). Formally, the *i*th node age is the *i*th order statistic of a uniform distribution, which is a Beta random variable with *α* = *i* and *β* = *n* + 1 − *i*. The variance of this Beta random variable approaches zero as *n* → ∞, indicating that the prior distribution on node ages becomes increasingly narrow as the number of tips increases (Figs. S46, S47). The exact distribution is somewhat different with extinct tips because the uniform random variables are not identically distributed, but the same general logic applies.

The distribution of node ages in a birth-death tree (of extant taxa) can also be represented by order statistics (Yang and Rannala 1997) and one may worry that they could exhibit similar behavior. However, in the case of birth-death models, the distribution of order statistics depends on the hyperparameters of the model (speciation and extinction rates), which are typically (effectively always) estimated from the data. The birth-death (and the fossilized birth-death) prior should therefore be able to adjust the distribution of order statistics to suit the data, especially if rates are allowed to vary over time. Indeed, the fossilized birth-death model with fixed hyperparameters also appears to be very informative, and can mimic the uniform tree model when rates are arbitrarily low (Fig. S48).

Consistent with its high informativeness, the uniform tree model had a profound impact on divergence times in our analyses (Figs. 2 and 3), and resulted in systematically increased estimates of clade ages (to the point of absurdity, such as Cambrian age for the Marattiales stem and Silurian for the crown group). On a finer scale, when compared to the fossilized birth-death models, the uniform tree model tends to spread node ages out as evenly as possible, which resulted in older ages for large, closely related clades, such as *Scolecopteris* and the extant Marattiaceae. These results are consistent with previous work, which reported unexpectedly ancient divergences under the uniform tree prior (Slater 2013; Wood et al. 2013; Arcila et al. 2015). As a consequence of clades being very old, there was also an increased opportunity for fossils to be nested within clades that would otherwise be too young to contain them, resulting in large differences in the inferred topologies (Figs. 2 and S50).

### Model evaluation: Which model is best?

There is strong evidence that rates of morphological evolution vary among characters: transition models that exclude gamma-distributed rates fail to generate patterns of variation among characters that are similar to those in the observed morphological dataset (Fig. S23). There is also evidence that the process of evolution is not the same among characters and states, as the F81 mixture models generally simulate more realistic datasets than the Mk models. Additionally, there is some evidence that the process of morphological evolution may vary among branches of the tree: removing either extinct or extant lineages from the simulated and observed morphological datasets results in somewhat different patterns of model adequacy, particular among morphological-transition models (Figs. S24 and S25).

Posterior-predictive simulations were unable to detect differences among the morphological clock models (Fig. S23). This result is somewhat surprising because, in principle, different posterior distributions of morphograms (as observed under the morphological-clock models, Fig. 2) could affect model adequacy. The apparent insensitivity to the morphological-clock model persists even after removing extinct lineages from the morphological datasets (Fig. S24), indicating that this is not simply a consequence of the topological and temporal distribution of fossils in our tree (*i*.*e*., a large clade of ancient fossils with branch lengths that do not depend on any molecular data).

All of the tree models estimated very similar morphograms (Fig. 2), and consequently, the posterior-predictive distribution for these models were effectively the same. The failure of the PPS approach to differentiate among the tree models, despite the extreme effect of these models on the resulting inference (Figs. 2, 3) is a manifestation of the nonidentifiability of this model component: differences in time (node heights) can be compensated for by differences in rates, with no effect on the expected distribution of the data. However, Bayes factors (which are sensitive to differences in prior distributions) clearly differentiated among the tree models (Table 1). The low marginal likelihood under the uniform tree model presumably reflects the fact that the clock rates necessary to achieve a reasonable fit to the data given the highly distorted node ages are implausible from the perspective of the relaxed-clock model.

The other major, and surprising, result of our Bayes factor comparisons was the relatively poor performance of the FBD models that allow fossilization rates to vary. Considering that our fossils are overwhelmingly from the Pennsylvanian, and that this concentration appears to be due (at least in part) to the unique preservation potential of that epoch (via the formation of coal balls), we expected that fossilization-rate variation would be an important model component. However, the number of fossils in a given time interval is a function of both the fossilization rate and the number of lineages in that time interval. The EFBD_*ψ*_ model struggles to produce enough lineages in the Pennsylvanian to generate a sufficient number of fossils, even with an elevated fossilization rate (Fig. S29); presumably, increasing the diversification rate would increase the number of lineages available to fossilize, but would also predict many more extant taxa than we observe. By contrast, the EFBD_*λ,µ*_ model implies a spike of diversity in the Pennsylvanian (Fig. 5, top), and is therefore able to explain both the large number of fossils in that time period and the observed diversity at the present. Allowing fossilization rates to vary on top of diversification-rate variation does not appear to improve model fit, *i*.*e*., the epoch-specific diversification-rate variation appears to be sufficient to describe the observed patterns of fossil and extant diversity.

**Figure 5:**
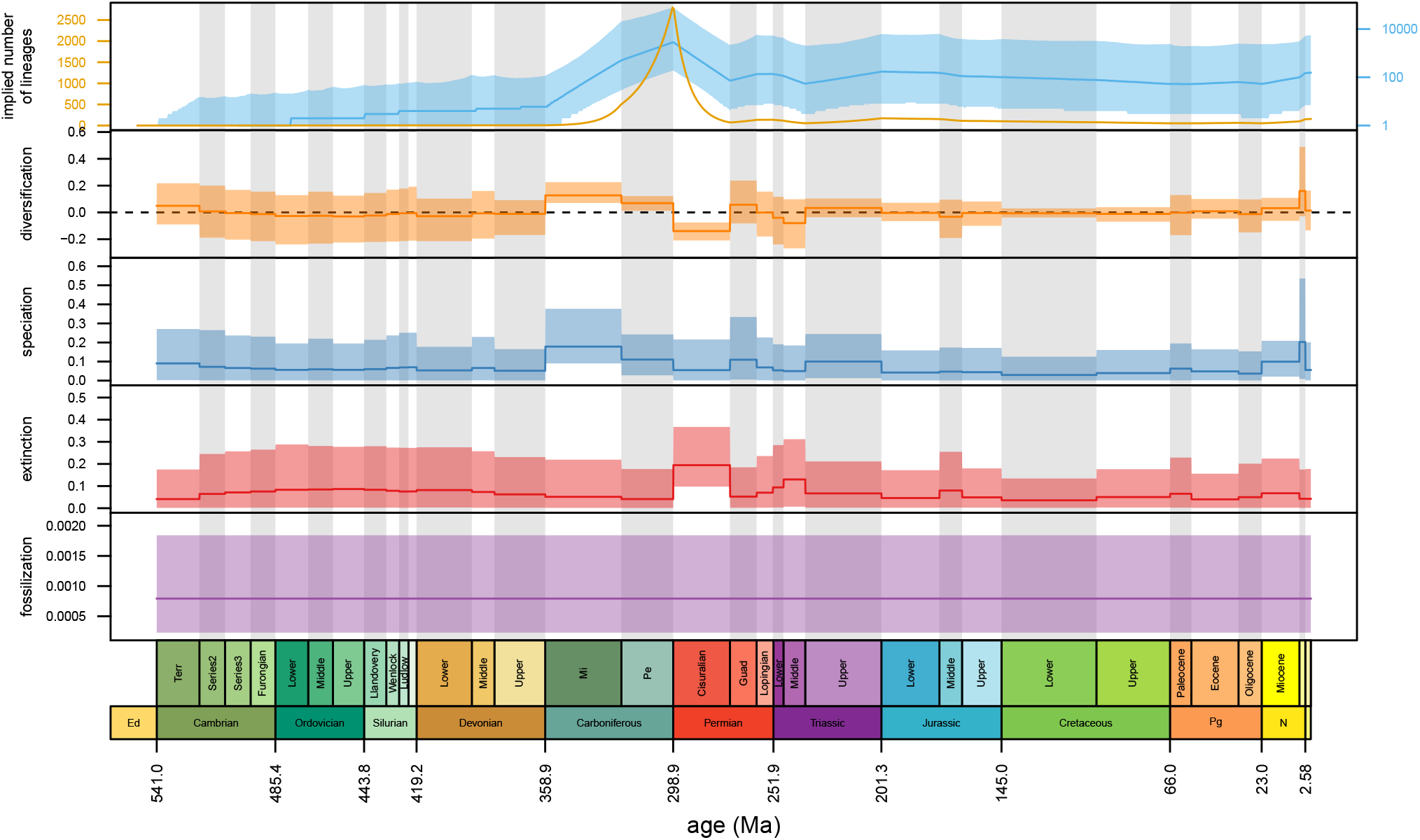
Diversification over time under the preferred model. The top panel shows the implied total number of lineages over time (before pruning unsampled lineages); the median value is shown on the linear scale in orange, and the median value and 95% credible interval are shown on a log scale, in blue. The next four panels correspond to the posterior distribution of epoch-specific net-diversification rates, speciation rates, extinction rates, and fossilization rates, respectively. In the top panel, the dark line represents the posterior median estimate, in the remaining panels it represents the posterior mean estimate. In all cases, and the shaded region corresponds to the 95% credible interval. Note that the preferred tree model, EFBD_*λ,µ*_, assumes that fossilization rates are constant (contrast against the model with fossilization-rate variation, Figure S28).

### Modeling prospects

While divergence-time estimates appear to be relatively insensitive to the morphological transition model—despite the fact that we investigated arguably the most biologically realistic morphological models applied in a TED analysis to date—these models still provide the opportunity to learn about the processes that govern morphological evolution. It is notable, for example, that the best overall morphological-transition model—the F81 mixture+Γ model—is also the most parameter-rich, and that even this model struggles to adequately model variation in the process of morphological evolution among branches of the tree (Figs. S24 and S25). These results may indicate that our morphological dataset could support even more complex models, for example, ones that allow the process of evolution to vary among branches (Beaulieu et al. 2013; Goloboff et al. 2019), or that accommodate correlated evolution among characters (Pagel and Meade 2006; Meyer et al. 2019) and other complex dependencies (Maddison 1993; Tarasov 2019). However, such models generally require increasing the state-space of the characters, which complicates the calculations used for correcting for variable-only characters, so more work is necessary before they should be used in total-evidence dating analyses. Our models also assume that morphological data subsets (defined by the number of states) evolve at different relative rates, and that rate variation within data subsets follows a gamma distribution. Models that accommodate variation in the rate and process of evolution using biologically meaningful data partitions (for example, partitioning between feeding and non-feeding characters, as in Wright et al. 2020, or between reproductive and vegetative characters, etc.) provide another opportunity to improve model realism.

Ultimately, our choice of candidate morphological clock models was guided by practical considerations, but, as with the morphological transition models, it is easy to imagine more biologically rich and meaningful models. Autocorrelated relaxed clocks, which assume that rates of evolution themselves evolve over the branches of the tree, are a biologically natural class of models with a long history in Bayesian divergence-time estimation (Heath and Moore 2014), and can accommodate gradual patterns of rate change (Thorne et al. 1998; Thorne and Kishino 2002), rare episodic changes (Huelsenbeck et al. 2000), clade-specific rates (Drummond and Suchard 2010), and complex mixtures of gradual and episodic change (Lartillot et al. 2016). We attempted to use autocorrelated Brownian motion and random-local-clock models (Thorne et al. 1998; Thorne and Kishino 2002; Drummond and Suchard 2010) in the early stages of our study, but were unable to make the MCMC analyses function adequately: as the number of branches in the tree grows, the dependency between the autocorrelated rates and the tree topology makes it increasingly difficult to efficiently sample over tree space. As total-evidence-dating approaches gain in popularity, it will be desirable to develop more efficient MCMC procedures for sampling tree and branch rates under auto-correlated models.

Consistent with previous studies (Zhang et al. 2016; Wright et al. 2020), our results suggest that the tree model can have a large—or, in the case of the uniform tree model, overwhelming—impact on total-evidence dating analyses. While the fossilized birth-death model improves our ability to jointly model extinct and extant diversity, and to have a biologically meaningful tree model, it is nonetheless limited in some important ways. For example, our results assume that the sampling fraction applies uniformly to all extant taxa. While the qualitative impact of each model component was consistent regardless of whether we assumed the total extant sampling fraction (*ρ* = 36 ÷ 12000) or the ingroup fraction (*ρ* = 27 ÷ 111; Supplemental Material S§6.2), exact age estimates are different between the two assumed sampling fractions, with the outgroup sampling fraction driving older age estimates (see Supplementary Material S§6.2). Unfortunately, neither of these sampling fractions is realistic, given that the sampling intensities for our in-group and outgroup are highly imbalanced.

While a model of “diversified” taxon sampling exists for the fossilized birth-death process (Zhang et al. 2016), this model assumes that all unsampled extant lineages attach to the tree more recently than the youngest internal node or fossil. This assumption is clearly inappropriate for the Marattiales, where most extant species arose relatively recently (Fig. 4). Given that empirical datasets frequently exhibit imbalanced taxon sampling schemes, and some degree of “diversified” or “deep-node” sampling (Matschiner 2019), an important avenue of development for the fossilized birth-death process will be to derive more flexible models of incomplete taxon sampling. Beyond the details of accommodating incomplete taxon sampling, the fossilized birth-death models we used in this study make many simplifying assumptions about the diversification and sampling process. For example, they assume that diversification and fossilization rates are the same among lineages and within each epoch, and do not depend on ecology, morphology, geography, etc. Tree models that allow for state-dependent and lineage-specific rates exist (in principle) for phylodynamic (epidemiological) models (Kü hnert et al. 2016), but while the theoretical machinery underlying the phylodynamic and fossilized birth-death models is nearly identical, to our knowledge these models have not been adapted for the fossilized birth-death model. They therefore represent a major untapped resource for future development and application.

Finally, our use of posterior-predictive simulation to assess the adequacy of morphological models assumes that the statistics we have chosen—the total parsimony score, *S*, and the variance in parsimony scores, *V*—are sensitive to realistic violations of our models. More work is needed to determine the power and utility of these model evaluation tools to assess different components of the total-evidence model.

### Marattiales phylogeny and divergence times

#### Extant relationships

Topologically, our results for extant relationships are generally consistent with other molecular studies of Marattiales phylogeny, including in supporting *Danaea* as sister to the rest of the extant marattialeans (Murdock 2008b; Lehtonen et al. 2020), and supporting the monophyly of each genus (Murdock 2008b; Rothwell et al. 2018b; Lehtonen et al. 2020). Among the extant taxa, the main point of uncertainty remains the position of *Christensenia*. Whereas Murdock (2008b) found *Christensenia* sister to the remainder of the *Christensenia* + *Marattia* s.str. + *Angiopteris* clade, in our results *Marattia* is in that position. However, support is relatively weak for these relationships in our results as well as in Murdock (2008b). Lehtonen et al. (2020), while likewise having only weak support, resolve the third possible relationship: *Christensenia* sister to *Marattia*, and that clade sister to *Angiopteris*. This phylogenetic uncertainty is mirrored in previous morphological cladistic studies (see Hill and Camus 1986), as well as the sampling-focused analyses of Rothwell et al. (2018b), which have supported each of these positions for *Christensenia*, in addition to placing it sister to *Danaea* or to all other extant genera. Morphologically, our topological resolution of *Christensenia* sister to *Angiopteris* is supported by spherical spores (Fig. S64), synangia oval in longitudinal section (Fig. S67), and a raised stomatal complex (Fig. S75). Regardless of its precise position, *Christensenia* is clearly nested within extant Marattiaceae and the characters that it shares with Psaroniaceae species (most notably, its radially symmetrical synangia; Fig. S1K) are independently derived, a result that contrasts strongly with pre-phylogenetic and early morphological cladistic hypotheses (*e*.*g*., Campbell 1911; Hill and Camus 1986).

The extant Marattiales are deeply isolated from their closest extant relatives, the leptosporangiate ferns, as has been repeatedly demonstrated (Pryer et al. 2001; Qiu et al. 2007; Murdock 2008b; Rai and Graham 2010; Lehtonen 2011; Rothfels et al. 2015b; Lehtonen et al. 2020). Given this situation—a very long stem branch connecting to a series of much shorter branches within the Marattiales crown group—one might expect uncertainty within the crown group relationships driven by uncertainty in the attachment position of the stem branch (Huelsenbeck et al. 2002; Rothfels et al. 2012), as in the maximum likelihood results of Murdock (2008b) and as has been seen in other similarly isolated groups (*e*.*g*., *Isoetes, Equisetum*, and *Cycas*; Des Marais et al. 2003; Schuettpelz and Hoot 2006; Nagalingum et al. 2011). With this concern in mind, we questioned whether *Danaea* was correctly inferred as sister to the remaining extant taxa. Our analyses allow for the long stem branch to be broken up by fossils, which may additionally provide important information about the polarity of morphological characters (*i*.*e*., which character states are ancestral for crown Marattiaceae), potentially allowing for more reliable inference of the “root” position for this crown clade (Doyle and Donoghue 1987; Gauthier et al. 1988; Donoghue et al. 1989; Huelsenbeck 1991; Smith 1998; Wills and Fortey 2000; Mongiardino Koch and Parry 2020). Nonetheless, our results further support the growing consensus (Pryer et al. 2001; Schuettpelz et al. 2006; Qiu et al. 2007; Murdock 2008b; Rai and Graham 2010; Lehtonen et al. 2020) that *Danaea* is sister to the remaining extant lineages; it shares synapomorphic states of the extant clade (*e*.*g*., the presence of stipules), and has apomorphic states (*e*.*g*., once-pinnate, dimorphic leaves), but few character states argue for it being nested among the other extant taxa; those that do appear to be homoplastic.

#### Overall relationships

Our results depart in significant ways from classic interpretations of Marattiales evolution (see Rothwell et al. 2018a), the topologies inferred by previous total-evidence analyses (Rothwell and Good 2000; Lehtonen et al. 2020), and the morphological cladistic analysis by Liu et al. (2000). Namely, in our MCC trees Psaroniaceae appears paraphyletic, comprising a grade of early-diverging lineages in Marattiales, a component of which eventually gave rise to the Marattiaceae (Figs. 4, S26). To a lesser extent, Lehtonen et al. (2020) found a similar pattern, with a polytomy comprised of *Araiangium, Danaeites rigida* + *Millaya tularosana*, and *Sydneia* + *Radstockia* sister to the remaining Psaroniaceae and Marattiaceae clades, and Liu et al. (2000) also recovered a non-monophyletic Psaroniaceae, even though they used a much different fossil sample.

In contrast to Rothwell et al. (2018b), we find little evidence for the alternative hypothesis that Psaroniaceae and Marattiaceae are reciprocally monophyletic clades that diverged from a common ancestor and thereafter had separate evolutionary histories (Figs. S54, S27). Nonetheless, we do resolve a large Psaroniaceae clade that includes most (Fig. 4) or all (Fig. S26) of the *Scolecopteris* species along with, at least, *Araiangium pygmaeum, Acaulangium bulbaceum, Convexocarpus distichus*, and *Buritiranopteris costata* + *Gemellitheca saudica*. Lehtonen et al. (2020) and Rothwell et al. (2018b) found several other genera to be included in the large Psaroniaceae clade—such as *Zhutheca, Taiyuanitheca, Acitheca, Pectinangium, Acrogenotheca, Symopteris*, and *Sydneia*—but we did not include these taxa in our analyses. Notably, *Radstockia* is inferred by both Rothwell and Good (2000) and Lehtonen et al. (2020) (the latter with *Sydneia*), to be sister to the rest of Marattiales (Psaroniceae + Marattiaceae), whereas our analyses place *Radstockia* well within the Marattiales, consistent with Hill and Camus (1986).

Overall, the grade of lineages recovered in our MCC trees (which, other than *Scolecopteris* species, was largely consistent among our different empirical consideration analyses) captures the morphological transitions accompanying the floristic turnover in the late Pennsylvanian and Permian that resulted in the modern Marattiaceae. These critical transitional forms, for example, *Eoangiopteris goodii, Millaya tularosana*, and *Radstockia kidstonii*, could be assignable to a broad interpretation of Marattiaceae, owing to their morphological departure from most Psaroniaceae, and specifically in their sessile bilateral synangia that contain numerous sporangia borne on flattened rather than downturned pinnules, and by having spores with an ornamented exine (Fig. S71). In other respects, however, these plants retain traits characteristic of Psaroniaceae, such as highly dissected leaves (Fig. S63). The picture that emerges is of a remnant of the formerly diverse Psaroniaceae that, through a series of morphological changes (some of which are captured in the fossil record) evolves into the ecologically and morphologically distinct modern Marattiaceae (Liu et al. 2000).

The position of *Escapia*—another potentially “transitional” taxon—is highly uncertain in our analyses, and its alternative phylogenetic resolutions have notable implications for the interpretation of the temporal history of the Marattiales, particularly the persistence of Psaroniaceae. *Escapia* is a fragmentary fossil (Rothwell et al. 2018a) that exhibits a unique suite of character states, including autapomorphies such as synangia served by transfusion tracheids (Fig. S68), and states that are otherwise found in the *Scolecopteris* clade, namely having both radial and bilateral synangia (Fig. S66), with ovate eusporangium cavities (Fig. S72), and extended sporangium tips (Fig. S69). In long-section, the synangia shape appear as two crescents (Fig. S67), a state that otherwise only occurs within our dataset in *Scolecopteris alta*. In consequence, in some of our analyses *Escapia* is resolved sister to *Scolecopteris alta* (Fig. S51), or nested among *Scolecopteris* species (Fig. S43), supporting the hypothesis that multiple lineages of Pennsylvanian Psaroniaceae persisted into the early Cretaceous and that psaroniaceous species may have extensively coexisted with members of the Marattiaceae (Rothwell et al. 2018a,b). However, in our favored model *Escapia* is resolved among stem groups more closely related to the extant clade (Fig. 4; see also Figs. S26, S55). In this case, *Escapia* would not be interpreted as the last vestige of the Psaroniaceae extending into the Cretaceous, but instead as another unusual transitional form, which converged upon synangial morphology similar to *Scolecopteris alta*.

The Permian *Qasimia schyfsmae* has been generally regarded as the oldest representative of the Marattiaceae (e.g., Hill et al. 1985; Hill and Camus 1986; Rothwell et al. 2018a) based on synangial characters and foliage similarities with *Marattiopis* species. Similar to other analyses (Rothwell and Good 2000; Lehtonen et al. 2020), we found *Qasimia schyfsmae* to be most closely related to the clade of *Marattiopsis* + extant Marattiaceae (Fig. 4), and in many cases we reconstruct it as a direct ancestor of the extant species (as was suggested by Hill and Camus 1986).

Our other major phylogenetic result, and one of our most surprising inferences, is that the extant marattialeans form a clade (with 0.96 posterior support) phylogenetically apart from any of the fossils (Fig. 4), despite the fact that a number of these fossils closely resemble extant taxa and have in the past even been considered congeneric with extant species (see discussions in Bomfleur et al. 2013; Escapa et al. 2015). This result only holds when temporal data are incorporated; in the non-dated analyses (Fig. S77) some of these fossils are resolved among the extant species, as they were in, *e*.*g*., Rothwell et al. (2018b) and Lehtonen et al. (2020).

The position of these fossil taxa outside the crown group is not completely unexpected: the extant genera mostly lack unique apomorphies and are instead defined by suites of states (see Murdock 2008a; Escapa et al. 2015). *Marattiopsis* species, in turn, are mostly based on fragmentary fossils, and generally exhibit combinations of states that do not match any of the extant genera. There is, therefore, enough homoplasy that the fossils could be placed in a number of positions among extant taxa or on the stem, as exemplified by their alternative placements in other analyses (Rothwell and Good 2000; Lehtonen et al. 2020; Liu et al. 2000).

#### Timescale of Marattiales evolution

Our mean Marattiales stem-age estimate (414 Ma, 95% CI = [351, 505]; Fig. 4, Table S.3) is older than, but broadly consistent with, earlier studies that have inferred ages for this node using node-dating approaches (*e*.*g*., ∼ 325 Ma (Rothfels et al. 2015b), ∼ 366 Ma (Testo and Sundue 2016), and ∼ 360 Ma (Qi et al. 2018); Shen et al. (2018) infer a much younger date, ∼ 250 Ma, but they also infer a different topology). Similarly, our inferred stem age for the Marattiaceae is consistent with other analyses, both when defined as the clade including extant taxa + *Marattiopsis* spp. + *Qasimia*, or more narrowly as the extant taxa + *Marattiopsis* spp. (Lehtonen et al. 2020; Rothwell et al. 2018b). Previous node-based estimates of the crown group age, however, have varied tremendously. On the younger end, Qi et al. (2018) estimated an age of ∼ 75 Ma, and Testo and Sundue (2016) place this node at ∼ 160 Ma. Even the 160 Ma estimate, however, is dramatically younger than the 185 *—* 224 Ma or ∼ 200 Ma inferences from Lehtonen et al. (2017) and Smith et al. (2010), respectively. The latter two inferences differed from the studies that inferred a younger crown age in assuming that a fossil marattialean was congeneric with an extant taxon, and thus they constrained the Marattiales crown node to be older than the fossil. Based on morphological similarities, this assumption is well-justified: parsimony analyses of morphology (*e*.*g*., Rothwell et al. 2018b; Lehtonen et al. 2020) consistently resolve fossils among the extant taxa, as do our non-clock trees (Fig. S77); the (effectively non-clock) parsimony-based dating analyses of Lehtonen et al. 2020 also infer a very old date, of ∼ 220 Ma).

#### The influence of temporal data on estimates of topology and divergence times

Discrepancies between the clock and non-clock analyses (see Fig. S77), and between node-based and TED analyses, provide a strong example of the potential for temporal data to alter our inferences of phylogenetic relationships (Drummond et al. 2006; Gavryushkina et al. 2017; Lee and Yates 2018; Wright et al. 2020). TED analyses, by co-estimating the position of the fossils and the divergence times of the tree, allow for both the morphological characteristics and the temporal data associated with the fossils (their ages) to influence their position in the phylogeny. Effectively, if the full model— incorporating morphological and temporal information from all the samples, as well the influences of the tree and clock priors—prefers a clade age that is younger, such that it overwhelms the specific morphological data that might resolve the fossil inside the clade, the model can place the fossil elsewhere. This outcome might be more likely if there is considerable homoplasy in the morphological data (as there is in extant Marattiales, and in fossil *Marattiopsis*; see Escapa et al. 2015), resulting in a high inferred rate of morphological evolution and a reasonable chance of repeated evolution of particular character states. We infer strong support for a monophyletic crown group comprising all extant taxa to the exclusion of any fossils (see Marattiales phylogeny and divergence times, above) and a relatively young age for the crown Marattiales (mean = 84 Ma, CI = [46, 169]; Fig. 4, Table S.4), in marked contrast to studies that assumed that fossils fell among the extant species. The Marattiales, then, provide a powerful illustration of dangers of the *a priori* assignment of fossils to nodes that is required for node-based divergence-time dating: phylogenetic positions of fossils that seem compelling based on morphology may not be supported in the context of the morphological and temporal data of the full sample (for a similar result, see Lee and Yates 2018).

### Reconciling model inferences with marattialean biology

#### Inferred diversification dynamics

While not the primary focus of our analyses, our application of the FBD model allowed us to infer the fine-scale (epoch-level) diversification dynamics of this clade, informed by the fossil record. Throughout its history, we estimate a diversification rate for the Marattiales near zero, with both speciation and extinction rates being low. There is some notable variation around this trend, however. Specifically, speciation rates increase in the Carboniferous (and particularly in the Mississippian), the middle Permian (Guadalupian), and to a lesser extent, the late Triassic, whereas extinction rates spike in the early Permian and lower and middle Triassic (the latter potentially reflecting the end-Permian mass extinction; Fig. 5). In combination, these rates result in a picture of Marattiales evolution where species-level diversity increases rapidly in the Pennsylvanian, culminating in a peak of ∼ 2800 species (median estimate) before crashing abruptly in the early Permian (Fig. 5). However, allowing fossilization rates to vary reduces the inferred peak diversity: the EFBD_*λ,µ,ψ*_ model, while disfavored over our optimal tree model by Bayes factors [Table 1], infers a peak of “only” ∼ 1200 species, which may be more biologically plausible. The other major pattern in our diversification inferences— the sudden spike in speciation rates in the Pliocene— coincides with the modern taxa and likely relates to a switch in species concepts rather than underlying biology.

Instead of reflecting the reality of the fossil record, the apparent lack of a signal of fossilization-rate variation may be due to a combination of confounded epoch-specific diversification and fossilization rates, taxonomic practice, our sampling regime, or simply a lack of power. Given that most preserved Pennsylvanian marattialeans were wetland species (DiMichele and Phillips 2002), the early Permian aridification of the tropics would have reduced both their fossilization potential and their diversity (Montañez et al. 2007); by modeling the latter, we may have been able to simultaneously account for the former. In addition, fossilization-rate variation is likely muted by the tendency for taxonomists to pay greater attention to taxa that are unexpected or otherwise interesting. In the case of the Marattiales, this bias results in small fragmentary Cretaceous fossils being described in great detail owing to their rarity (*i*.*e*., *Escapia*; Rothwell et al. 2018a) and therefore included in our dataset, while the great bulk of the fossils—fine anatomical preservations in Pennsylvanian coal balls—are relatively under-represented. Our sampling regime likely exacerbates this bias: of the many Carboniferous and Permian Marattiales fossils, we included relatively few (*e*.*g*., only 12 of the over 30 described genera; Lundgren et al. 2019; Rothwell et al. 2018a). A final potential explanation is that we simply do not have enough information to distinguish between models with and without fossilization-rate variation, and that with more fossil data we would have rejected the constant-fossilization-rate model. Regardless, our primary results are nearly the same when we allow fossilization rates to vary (Figs. 2, 3).

#### A paleoecological perspective

While the Devonian origin time inferred here for the Marattiales significantly predates any known fossils, we find that the major initial diversification of the Marattiales likely began in the Mississippian (359 − 323 Ma). Although there is not a lot of unequivocal evidence of Mississippian marattialeans in the macrofossil record, such a pattern is expected for the initial diversification of a group; the existence of only a few lineages, likely with small geographical ranges, would lead to exceedingly low probability of fossil recovery, which is compounded by the difficulty in recognizing, or preserving, the defining characteristics of a group early in its evolution (Marshall 2019).

The first potential macrofossil evidence of marattialeans, from the earliest stage of the Mississippian (Tournaisian, 359 − 347 Ma), is *Burnitheca pusilla*, an isolated permineralized synangium (Meyer-Berthaud and Galtier 1986). Compression fossils of trunks with distichous or helically arranged leaf scars have also been described from Mississippian localities (Crookall 1959); these fossils have a growth habit similar to early Marattiales taxa, but, like *Burnitheca pusilla*, lack the details needed to confidently assign them to the order. Specimens of the genus *Megaphyton*, stem compression and impressions from latest Mississippian-age sediments, are generally regarded as the oldest evidence for Marattiales (Pfefferkorn et al. 1976).

The relatively small size of the early specimens, and the absence of more delicate leaf fragments, indicate that they likely represent allochtonous plant material, meaning they were transported from their habitat before ending up in the depositional environment in which they were preserved (Greenwood 1991). This pattern implies that marattialeans initially grew in habitats with a low preservation potential, which could reconcile their scarcity in the fossil record with the high diversification rates we infer during the Mississippian (Fig. 5). During the Mississippian-to-Pennsylvanian transition, the climate in the Euramerican tropics during the glacial intervals changed from seasonally dry to tropical everwet (Calder and Gibling 1994); both the wetter climate and the associated expansion of peat swamp habitats substantially increased chances of structural preservation of the swamp vegetation (Cecil 2003; Gastaldo and Demko 2011). The earliest commonly accepted, unequivocal marattialeans are stems known from these conditions of high fossilization potential.

All finds suggest that the early Psaroniaceae were trees of relatively small stature, with monocyclic steles, distichous leaf arrangement, and a small root mantle (DiMichele and Phillips 1977; Millay 1997). It was not until the middle Pennsylvanian that Psaroniaceae species became important understory elements and common canopy trees (with thick root mantles and large decom-pound leaves) in lowland clastic swamp communities and more widely distributed in Euramerica (DiMichele and Phillips 2002; Millay 1979, 1997). After an initial decline during the late Middle Pennsylvanian lycopod collapse, Psaroniaceae further increased in size and ecological importance, became canopy dominants in the clastic swamps, and firmly established themselves in the dryer part of the peat swamps (Cleal 2015; DiMichele and Phillips 1996, 2002; Phillips et al. 1985). These, and the earlier taxa, had highly dissected leaves and synangia with small numbers of sporangia, traits that we reconstruct as persisting until the early Permian (Figs. S62, S65). The well-known members of the Psaroniaceae, including most in our analysis, are plants from these peat swamps, depositional environments where chances of preservation were high. Given that we have a good idea how plant communities changed over time in the wetter parts of the landscape, less information from the local drier habitats, and very little information on the evolution of plants and their communities in the extrabasinal environments (Looy et al. 2014), the drop in diversification in the early Permian (Cisuralian; Fig. 5) likely represents the extinction of the swamp taxa, associated with the aridification of the Euramerican tropics (Montañez et al. 2007).

Marattialeans produce massive amounts of spores and more than a dozen dispersed spore taxa have been found *in situ* in marattialean sporangia (for an overview see Balme 1995). The taxonomic resolution of these spores is quite variable; some are known from multiple distantly related taxa, while others have only been recorded from members of the Psaroniaceae or Marattiaceae (see *e*.*g*., Millay and Taylor 1984; Lesnikowska 1989; Lesnikowska and Willard 1997). The more taxonomically restricted taxa, including the genera *Fabasporites, Spinosporites, Thymospora, Torispora*, and smaller forms of *Cyclogranisporites* and *Laevigatosporites*, can be used as evidence for Marattiales in the absence of distinctive larger fossils (Lesnikowska 1989; Looy and Hotton 2014). Combined with the initial rarity of the marattialeans in the fossil record, the spore record corroborates our inferred diversification patterns (Fig. 5). Small amounts of minute *Cyclogranisporites* and *Punctatisporites* (Lesnikowska 1989) spores have been described from the Tournasian on, and are followed by a stepwise increase in species diversity and abundance when marattialeans become dominant elements of the swamp communities.

## Conclusion

While our focus was on the Marattiales, our results pertain to the general power and utility of the total-evidence dating framework, in conjunction with the fossilized birth-death class of tree models. The arguments in support of TED approaches and FBD models are not trivial: as our results show, different modeling choices can cause dramatic differences in the resulting phylogenetic inferences, owing in large part to the nonidentifiability inherent to these models. Fortunately, statistical models are accompanied by a robust toolkit that allows us to assess the influence of model and prior specification on parameter estimates, to compare the fit of competing models, to assess their ability to describe the true data-generating process, and to identify ways in which the models can be made more realistic. These features are most available in biologically interpretable models, like the FBD model, because the parameter estimates can be compared against empirical expectations. With the support of this toolkit, these modeling choices transform from an analytical nuisance into an opportunity to learn about the processes that produced our data, and subsequently to identify avenues for increasing the biological realism of our models.

By applying this toolkit to the Marattiales, we were able to infer a nuanced picture of the evolution and diversification of this clade over its ∼ 400-million-year history. This inference was possible despite the fact that much of the abundant Marattiales fossil record was left by lineages without extant descendants, the extant taxa are a young clade very distant from their closest living relatives, and the placement of fossils is compromised by their often fragmentary nature and morphological homoplasy among fossil and extant species. We infer that the Marattiales began to diversify in the Mississippian, prior to a well-established fossil record. Considerations of the ecology of potential early marattialeans suggest that this timescale may be reasonable: early marattialeans appear to be small, uncommon taxa, which occurred in habitats with low preservational potential. We also infer that the Marattiales experienced peak diversity at the end of the Pennsylvanian, before a sharp decline in the Permian to relatively stable levels of standing diversity that persisted throughout the Mesozoic to the present. Again, this pattern makes sense in light of the ecology of marattialeans and our understanding of paleoclimate: the relatively wet climate in Euramerica during the Pennsylvanian would have been ideal for the proliferation of wetland-adapted marattialeans, while subsequent aridification of that region during the early Permian would have driven high rates of extinction. The broad concordance of phylogenetic, ecological, and paleoclimatological evidence demonstrates the potential for total-evidence dating—particularly in conjunction with the fossilized-birth-death model—not only to harmonize “rocks and clocks”, but also to elucidate macroevolutionary processes.

## Supplementary Material

Supplementary scripts and data can be found in the Data Dryad repository DOI:X and the GitHub repository https://github.com/mikeryanmay/marattiales_supplemental/releases/tag/1.0.

## Funding

This research was supported by the National Science Foundation (NSF) grant DEB-1754705 to CJR and CVL, DEB-1754723 to NSN, and DEB-1754385 to MAS.

## Acknowledgements

We thank Andy Murdock, members of the Rothfels lab, Jack Tseng, Ixchel González Ramírez, Ziad Khouri, Xavier Meyer, and Brian Moore for their thoughtful discussion, Bill DiMichele, Scott Elrick, and Ignacio Escapa for images of fossil specimens, and associate editor Ryan Folk and three anonymous reviewers for providing suggestions that greatly improved the manuscript.

This research used the Savio computational cluster resource provided by the Berkeley Research Computing program at the University of California, Berkeley (supported by the UC Berkeley Chancellor, Vice Chancellor for Research, and Chief Information Officer).

## Notes

https://github.com/mikeryanmay/marattiales_supplemental/releases/tag/1.0

### Competing Interest Statement

The authors have declared no competing interest.

### Summary of Updates

Paper accepted in Systematic Biology.

