## supplemental material for "Inferring the Total-Evidence Timescale of Marattialean Fern Evolution in the Face of Model Sensitivity"

### Supplemental Material for: Inferring the Total-Evidence Timescale of Marattialean Fern Evolution in the Face of Extreme Model Sensitivity

#### Contents


**S§1 Marattiales Data** **S7**
S§1.1 Morphological Data . . . . . S8
S§1.2 Accessions . . . . . S11

**S§2 Graphical Models** **S17**
S§2.1 Substitution Model . . . . . S17
S§2.2 Molecular Clock Model . . . . . S18
S§2.3 Morphological Transition Models . . . . . S19
S§2.4 Morphological Clock Models . . . . . S22
S§2.5 Tree Models . . . . . S24

**S§3 MCMC Analyses** **S29**
S§3.1 MCMC Diagnosis . . . . . S29
S§3.2 Computational Details . . . . . S30

**S§4 Posterior-Predictive Simulation** **S31**

**S§5 Empirical Considerations: Taxon Sample and Rooting Strategies** **S32**
S§5.1 Ancient plants . . . . . S32
S§5.2 Polarized rooting . . . . . S32
S§5.3 Ingroup-only . . . . . S33
S§5.4 Results . . . . . S33

**S§6 Extended Results** **S35**
S§6.1 Ingroup Sampling Fraction . . . . . S35
S§6.2 Overall Sampling Fraction . . . . . S48
S§6.3 Ingroup vs. Overall Sampling Fraction . . . . . S61
S§6.4 Uniform Tree Model . . . . . S62
S§6.5 Polarized Analysis . . . . . S67
S§6.6 Ancient Plants Analysis . . . . . S69

S§6.7 Ingroup Analysis . . . . . S71
S§6.8 Comparing Empirical Considerations . . . . . S73
S§6.9 Extant Phylogenies . . . . . S77
S§6.10 Stochastic Maps . . . . . S79
S§6.11 Dated vs. Non-Dated Topologies . . . . . S94

#### List of Figures


S1 Sample of Marattiales fossil and extant diversity . . . S7
S2 The GTR+I+$\Gamma$ substitution model . . . S17
S3 The uncorrelated lognormal (UCLN) relaxed molecular clock model . . . S18
S4 The Mk morphological model . . . S19
S5 The Mk+$\Gamma$ morphological model . . . S19
S6 The F81 mixture morphological model . . . S20
S7 The F81 mixture +$\Gamma$ morphological model . . . S21
S8 The "linked" relaxed morphological model . . . S22
S9 The "unlinked" relaxed morphological model . . . S23
S10 The uniform tree model . . . S24
S11 The CRFBD tree model . . . S25
S12 The EFBD$_{\psi}$ tree model . . . S26
S13 The EFBD$_{\lambda,\mu}$ tree model . . . S27
S14 The EFBD$_{\lambda,\mu,\psi}$ tree model . . . S28
S15 Comparing distributions of molecular trees among model combinations . . . S35
S16 Comparing distributions of extinct trees among model combinations . . . S36
S17 Comparing distributions of trees among model combinations, excluding the uniform tree model . . . S37
S18 Comparing lineage-through-time curves among model combinations . . . S38
S19 Comparing lineage-through-time curves of extant taxa among model combinations . . . S38
S20 Comparing divergence-time estimates for each clade by morphological transition model S39
S21 Comparing divergence-time estimates for each clade by morphological clock model . . S40
S22 Comparing divergence-time estimates for each clade by tree model . . . S41
S23 Comparing model adequacy among model combinations . . . S42
S24 Comparing model adequacy for extant taxa among model combinations . . . S42
S25 Comparing model adequacy for extinct taxa among model combinations . . . S43
S26 The maximum clade credibility tree under the preferred model . . . S44
S27 The 20%-rule consensus tree under the preferred model . . . S45
S28 Diversification over time under the EFBD$_{\lambda,\mu,\psi}$ model . . . S46
S29 Diversity through time under each fossilized birth-death model . . . S47
S30 Comparing distributions of trees among model combinations using the overall sampling fraction . . . S48
S31 Comparing distributions of molecular trees among model combinations using the overall sampling fraction . . . S49
S32 Comparing distributions of extinct trees among model combinations using the overall sampling fraction . . . S50
S33 Comparing distributions of trees among model combinations using the overall sampling fraction, excluding the uniform tree model . . . S51
S34 Average lineage-through-time curves among model combinations using the overall sampling fraction . . . S52
S35 Comparing lineage-through-time curves among model combinations using the overall sampling fraction . . . S52
S36 Comparing lineage-through-time curves of extant taxa among model combinations using the overall sampling fraction . . . S53
S37 Comparing divergence-time estimates for each clade by morphological transition model using the overall sampling fraction . . . S54

S38 Comparing divergence-time estimates for each clade by morphological clock model using the overall sampling fraction . . . . S55
S39 Comparing divergence-time estimates for each clade by tree model using the overall sampling fraction . . . . S56
S40 Comparing model adequacy among model combinations using the overall sampling fraction . . . . S57
S41 Comparing model adequacy for extant taxa among model combinations using the overall sampling fraction . . . . S57
S42 Comparing model adequacy for extinct taxa among model combinations using the overall sampling fraction . . . . S58
S43 The maximum clade credibility tree under the preferred model with overall sampling fraction . . . . S59
S44 The 20%-rule consensus tree under the preferred model with overall sampling fraction . S60
S45 Comparing divergence-time estimates for each clade by assumed sampling fraction . . S61
S46 The uniform tree model is an informative prior on node ages . . . . S62
S47 The uniform tree model is an informative prior on lineage-through-time curves . . . . . S63
S48 The fossilized birth-death model is an informative prior on node ages when hyperparameters are fixed . . . . S64
S49 The maximum clade credibility tree under the uniform tree model . . . . S65
S50 The 20%-rule consensus tree under the uniform tree model . . . . S66
S51 The maximum clade credibility tree under the preferred model with polarized root . . . S67
S52 The 20%-rule consensus tree under the preferred model with polarized root . . . . . . . S68
S53 The maximum clade credibility tree under the preferred model with ancient plant fossils S69
S54 The 20%-rule consensus tree under the preferred model with ancient plant fossils . . . . S70
S55 The maximum clade credibility tree under the preferred model and just ingroup taxa . . S71
S56 The 20%-rule consensus tree under the preferred model and just ingroup taxa . . . . . . S72
S57 The impact of empirical considerations on phylogenetic estimates . . . . S73
S58 The impact of modeling and empirical considerations on divergence-time estimates of major clades . . . . S73
S59 Comparing divergence-time estimates for each clade with different empirical considerations . . . . S75
S60 The maximum clade credibility trees for extant taxa among tree models . . . . S77
S61 The maximum clade credibility trees for extant taxa among empirical datasets . . . . . . S78
S62 Stochastic map for character 23: degree of pinnation . . . . S80
S63 Stochastic map for character 28: foliar abaxial idioblasts . . . . S81
S64 Stochastic map for character 52: synangial suture between valves . . . . S82
S65 Stochastic map for character 54: number of sporangia per synangium . . . . S83
S66 Stochastic map for character 55: synangium symmetry in x.s. . . . . S84
S67 Stochastic map for character 57: synangium shape in long section pre-dehiscence . . . . S85
S68 Stochastic map for character 58: sporangium or synangium pedicel or receptacle histology S86
S69 Stochastic map for character 59: sporangium tip extension . . . . S87
S70 Stochastic map for character 62: bilateral synangium dehiscence . . . . S88
S71 Stochastic map for character 70: spore ornamentation location . . . . S89
S72 Stochastic map for character 75: eusporangium cavity shape . . . . S90
S73 Stochastic map for character 77: synangium sporangia spread out from center on dehiscence . . . . S91
S74 Stochastic map for character 78: synangium location on pinnule . . . . S92
S75 Stochastic map for character 87: annulus of thick-walled cells . . . . S93

S76 The maximum clade credibility tree under the non-dated (non-clock) model . . . . . . . S95
S77 Topological conflict beween dated and non-dated MCC trees . . . . . . . . . . . . . . . . . S96
S78 Topological conflict beween dated and non-dated MRC trees . . . . . . . . . . . . . . . . . S97

### S§1 Marattiales Data

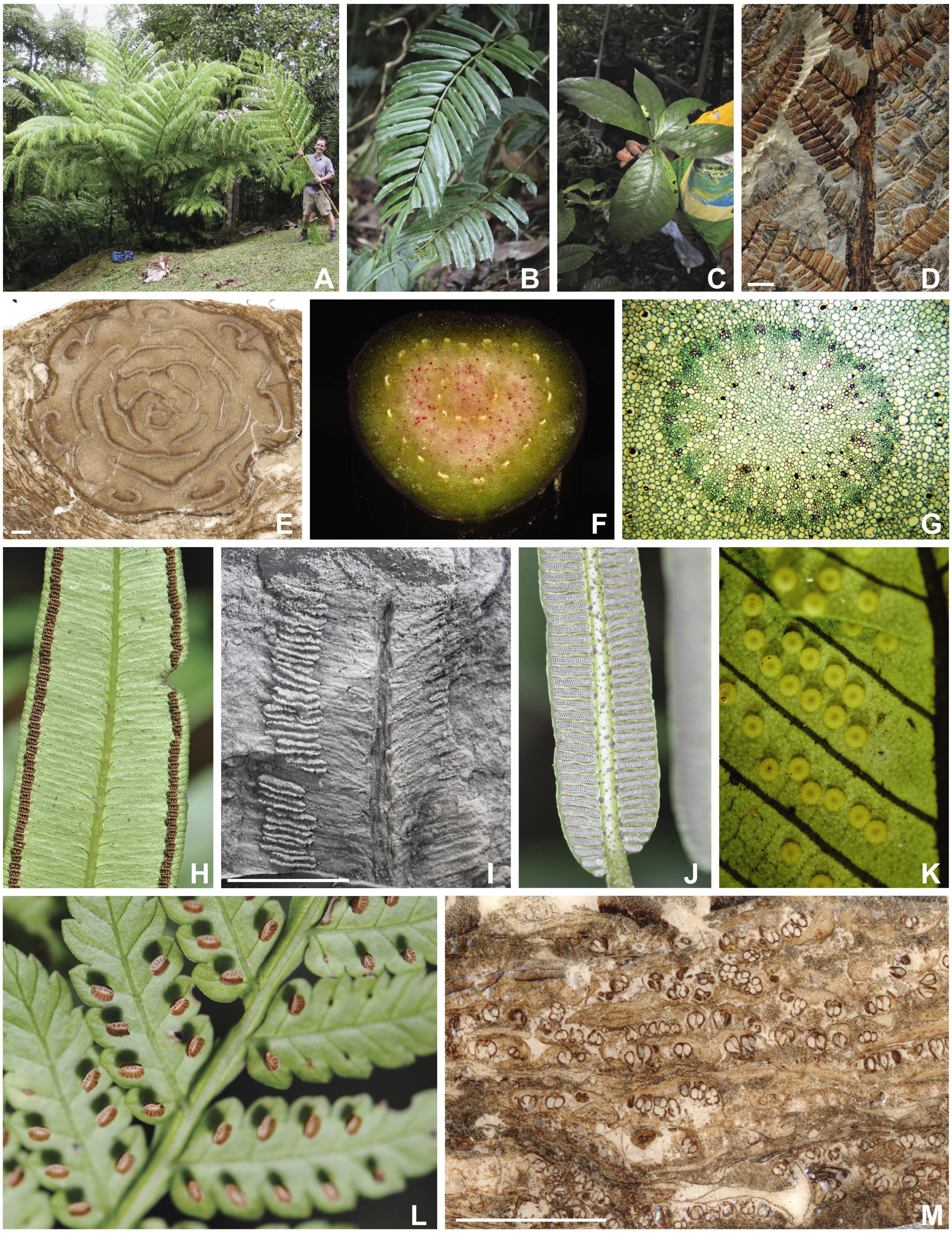


**Figure S1: Sample of Marattiales fossil and extant diversity.** A–D: Growth habit and frond morphology. A: *Angiopteris evecta*, Costa Rica, image credit R.C. Moran. B: *Danaea cuspidata*, Costa Rica, image credit W.L. Testo. C: *Christensenia aesculifolia*, Papua New Guinea, image credit M. Sundue. D: *Pecopteris*, West Fork Trinity, Texas, Late Pennsylvanian, USNM number 546865, image credit W.A. DiMichele. E–G: Stem, petiole, and root anatomy in cross section. E: Cross section of *Psaronius melandrus* higher up in the trunk, and an oblique section of inner and outer root mantle. University of Illinois Coal Ball Teaching Collection 10F, Middle Pennsylvanian, Herrin Coal Member, Sahara Coal Company Mine, Illinois, image credit S.D. Elrick. F: Petiole cross section, *Eupodium kalfusii*, Brazil, image credit F.B. Matos. G: Root cross section of *Angiopteris evecta* showing polyarchy, image credit R.C. Moran. H–M: Examples of synangia morphology. H: *Angiopteris evecta*, Papua New Guinea, image credit M. Sundue. I: Silicone cast of *Marattiopsis patagonica*, Early Jurassic of Patagonia, Argentina, MPEF-Pb 5295, image credit I.H. Escapa. J: *Danaea cuspidata*, Costa Rica, image credit W.L. Testo. K: *Christensenia aesculifolia*, Papua New Guinea, image credit M. Sundue. L: *Eupodium laeve*, Puerto Rico, image credit M. Sundue. M: *Scolecopteris* pinnules, University of Illinois Coal Ball 30932E. Middle Pennsylvanian, Calhoun Coal Member, Bonpas Creek, North Caroline, image credit S.D. Elrick. (D, E, I, M: scale bar = 5 mm.)

#### S§1.1 Morphological Data

Our morphological dataset was largely derived from Rothwell et al. (2018b), which itself relied heavily on Hill and Camus (1986) and Murdock (2008), and amended as necessary. We based character and character-state circumscriptions, as well as the character coding itself, on specimens examined at VT and on additional literature (Holttum 1978; Rolleri 1993; Rolleri et al. 2003; Christenhusz 2010a,b; He et al. 2013; Senterre et al. 2014). In a departure from Rothwell et al. (2018b)—which simplified stomata and pulvinus characters, and altered coding to avoid nested states—we instead used contingent character coding (Forey and Kitching 2000; Brazeau 2011). Our coding for the outgroup taxa was based on literature and our examination of specimens. Our new characters were primarily focused on the leptosporangiate ferns (our outgroup), *e.g.*, the presence of linear aerophores (character 27) and morphology and variation within leptosporangia (characters 93 − 98). Our final morphological matrix comprised 98 discrete characters describing anatomy and gross morphology; in total, there were 79 binary characters, 10 three-state characters, four four-state characters, three five-state characters, one six-state character, and one seven-state character. Twenty-two percent of the matrix was unscored due to contingent character coding (*i.e.*, inapplicable characters for a specific taxon), and an additional eight percent of the data were missing because they are unknown.

Here we provide our final list of characters and character states. Except where noted, characters and states are from Rothwell et al. (2018b), which incorporated characters and states from Hill and Camus (1986) and Murdock (2008). Character coding primarily follows Rothwell et al. (2018b) with some corrections.

1. Leaf-bearing stems : erect (0); creeping (1).
2. Erect adult stem : squat (0); elongate (1).
3. Stipule (aphlebia) pairs at leaf base : absent (0); present (1).
4. Root mantle around stem : absent (0); present (1).
5. Sclerified pith in root stele : absent (0); present (1).
6. Root cortex with continuous sclerenchyma band : absent (0); present (1).
7. Polyarch root stele : absent (0); present (1).
8. Multicellular root hairs : absent (0); present (1).
9. Mucilage canals in root : absent (0); present (1).
10. Stem symmetry (external morphology) 2 : radial (0); dorsiventral (1).
11. Basal pinnae more elaborated : absent (0); present (1).
12. Stem stele : protostele (0); solenostele (1); dictyostele (2); equisetostele (3).
13. Protoxylem development : exarch (0); mesarch (1); endarch (2).
14. Stem hypodermis of sclerenchyma : absent (0); present (1).
15. Stem/pinna internal sclerenchyma strand : absent (0); present (1).
16. Lysigenous lacunae in stem/pinnae : absent (0); present (1).
17. Mucilage canals in axial components of frond : absent (0); present (1).
18. Tannin cells in stem or pinnae cortex : absent (0); present (1).
19. Gum sacs in stems or petioles : absent or sparse (0); present (1).
20. Petiole or pinna vascular strand : continuous (0); discontinuous (1).
21. Phloem maturation : exarch (0); endarch (1).
22. Circinnate vernation : absent (0); present (1). New character
23. Number of fronds (megaphylls) : multiple fronds produced (0); one frond produced at a time (1).
24. Petiole trace : single undifferentiated trace (0); C-shaped (1); Omega-shaped (2); dissected Omega (3); multiple concentric rings of dissected traces (4); 3-shaped (5); elongate (6). New character.
25. Pulvinuloids at base of segments : absent (0); present (1). New character. A simplification of

Rothwell et al. (2018b) characters 109–113.

26. Polycylic dictyostele : absent (0); present (1). New character. Coding contingent on dictyostele being present (character 12, state 2).
27. Aerating tissue upon petiole : lateral linear aerophores (0); scattered and lenticel-like (1). New character based upon **???**.
28. Degree of pinnation : once (0); twice (1); three or more (2).
29. Ultimate segments articulated with sutures : absent (0); present (1).
30. Pinnule pinna base : margins parallel (0); narrowly confluent/cordate (1).
31. Pinnule pinna tips : acute (0); obtuse (1).
32. Epidermal cell walls thickened : absent (0); present (1).
33. Epidermal cell wall thickening : even (0); uneven (1).
34. Foliar abaxial idioblasts : absent (0); present (1).
35. Laminar idioblast groupings : solitary or small groupings (0); dense areas of idioblasts (1).
36. Idioblast chains on lateral veins : absent (0); present (1).
37. Idioblasts on synangium walls : absent (0); present (occasional) (1).
38. Scale cell shape : isodiametric (0); elongate (1).
39. Peltate scales : absent (0); present (1).
40. Peltate laminar scale morphology : with centrally attached stalks (0); with strongly asymmetrical stalk attachment (1).
41. Pinna/pinnule epidermal trichomes : absent (0); present (1).
42. Glandular trichomes : absent (0); present (1).
43. Scale margins : entire (0); lobed or fimbriate (1).
44. Sporangia individually vascularized : absent (0); present (1).
45. Sporangium/synangium base trichomes : absent (0); present (1).
46. Pinnule venation : simple (0); open dichotomous (1); reticulate (2).
47. Awns on veins : absent (0); present (1).
48. Fertile veins : all potentially fertile (0); preference towards distal vein branches (1).
49. Frond dimorphism : absent or slight (0); strongly dimorphic (1).
50. Synangia (sori) separated by surface foliar partitions : absent (0); present (1).
51. Lamina thickness : chartaceous to coriaceous (0); membranaceous (1). New character
52. Leaf cuticle (adaxial) : thin (0); thick (1).
53. Mesophyll cells with microscopic projections on exterior walls : absent (0); present (1).
54. Fertile pinnule margin morphology : flat (0); downturned (1); thin inrolled (2); thick inrolled (3).
55. Sporangia enclosed by pinnule margin : absent (0); present (1).
56. Functional sporangium annulus : absent (0); present (1).
57. Sporangia developmentally attached (synangium) : absent (0); present (1).
58. Extent of lateral sporangium fusion before dehiscence : none (0); partial (1); complete-almost complete (2).
59. Synangial suture between valves : deeply cut without central tissue pad (0); shallowly cut with central tissue pad (1).
60. Sporangia of sorus around central cellular area at base : present (0); absent (1).
61. Number of sporangia per synangium : 2–5 (0); 6–11 (1); 12–22 (2); more than 22 (3).
62. Synangium symmetry in x.s. : radial (0); radial and bilateral (1); bilateral (2).
63. Synangium receptacle morphology : flat or raised mound (0); radial stalk or pedicel (1).
64. Synangium shape in longitudinal section pre-dehiscence : cordate (0); fusiform (1); oval (2); two crescents (3).
65. Sporangium/synangium pedicel/receptacle histology : vascular (0); vascular + parenchyma (1); + (2); fiber + parenchyma (3); fiber (4); transfusion tissue (5).

66. Sporangium tip extension : present (0); absent (1).
67. Sporangium tip extension length : long, obvious (0); short, inconspicuous (1).
68. Initial synangium dehiscence : sporangia separate distally (0); sporangia do not separate distally (1); two rows of fused sporangia (valves) separate (2).
69. Bilateral synangium dehiscence : sporangial rows fused but not opening as two valves (0); sporangial rows opening as two valves (1).
70. Eusporangium aperture shape : pore (0); slit (1).
71. Eusporangium dehiscence position : inner/side facing wall (0); terminal (1).
72. Eusporangium aperture labiate : simple (0); labiate (1).
73. Eusporangium dehiscence area wall uniseriate : absent (0); present (1).
74. Spore suture : trilete (0); monolete (1); alete (2).
75. Spore morphology : spherical (0); ovoid (1); reniform (2); tetrahedral (3). Altered from Rothwell et al. (2018b), 2018 character 93 to include state 3.
76. Perine (perispore) : present (0); absent (1).
77. Spore ornamentation location : perine (perispore) (0); exine (1); perine and exine (2).
78. Perine (perispore) ornamentation units : smooth (0); spines (1); papillae (2); warts (3); rods (4). Altered from Rothwell et al. (2018b) character 96 to include state 4.
79. Exine ornamentation units : smooth (0); spines (1); papillae (2); warts (3); ridges (4).
80. Spore ornamentation units joined : absent (0); present (1).
81. Spore lasurae raised/thickened : absent (0); present (1).
82. Eusporangium cavity shape : cylinder tapering near tip (0); ovate wider in basal half (1); obovate wider in distal half (2). should the first one be "ovate"?
83. Eusporangium cavity length (mm) : greater than 1.0 (0); less than 1.0 (1).
84. Synangium sporangia spread out from center on dehiscence : absent (0); present (1).
85. Synangium location on pinnule : marginal (0); between margin and midvein (1).
86. Stipe pulvini : absent (0); present (at least in juvenile) (1).
87. Stomata of lamina cell arrangement : anomocytic to weakly cyclocytic (0); mostly cyclocytic (1); paracytic (2).
88. Stomata of lamina function : guard cells functional (0); guard cells permanently open (1).
89. Stomatal shape : elliptical (0); circular (1).
90. Stomatal complex : surficial or slightly sunken (0); raised (1).
91. Stomatal density : low (0); high (1).
92. Sporangium cavity base immersed in parenchyma : absent (0); present (1).
93. Number of sporangial initials : multiple (0); one (1). New character.
94. Annulus of thick-walled cells : absent (0); present (1). New character.
95. Position of annulus : lateral (0); oblique (1); transverse (2); apical (3); vertical (4). New character.
96. Sporangium wall thickness : thick walled (0); thin walled (1). New character.
97. Number of spores per sporangium : hundreds (0); 64 (1). New character.
98. Sorus protected by funnelform involucre : absent (0); present (1). New character.

#### S§1.2 Accessions

**Table S.1: Molecular Accessions**

| Taxon | atpB | rbcL | rps4-trnS | trnS-trnG+trnG |
|---|---|---|---|---|
| *Angiopteris caudata* | — | — | EU439172.1 | — |
| *Angiopteris evecta* | EU439071.1 | EU439092.1 | EU439139.1 | EU439228.1 |
| *Angiopteris henryi* | — | DQ838062.1 | — | — |
| *Angiopteris itoi* | EU439073.1 | EU439094.1 | EU439170.1 | EU439247.1 |
| *Angiopteris lygodiifolia* | X58429.1 | X58429.1 | EU439163.1 | EU439241.1 |
| *Angiopteris smithii* | EU439072.1 | EU439093.1 | EU439169.1 | EU439246.1 |
| *Angiopteris tonkinensis* | — | DQ838058.1 | — | — |
| *Christensenia aesculifolia* | EU439057.1 | EU439079.1 | EU439102.1 | EU439184.1 |
| *Cyathea multiflora* | EF463365.1 | AM410197.1 | FN667568.1 | — |
| *Danaea elliptica* | EU439054.1 | AF313578.1 | EU439096.1 | EU439178.1 |
| *Danaea grandifolia* | EU221704.1 | EU221766.1 | — | — |
| *Danaea leprieurii* | — | EU439077.1 | EU439097.1 | EU439179.1 |
| *Danaea nodosa* | — | EU439078.1 | EU439098.1 | EU439180.1 |
| *Diplopterygium glaucum* | KU877749.1 | KU936583.1 | KU936648.1 | — |
| *Dipteris conjugata* | AY612696.1 | EF588692.1 | AY612658.1 | — |
| *Eupodium kaulfussii* | — | — | EU439105.1 | EU439187.1 |
| *Eupodium laeve* | — | EU439081.1 | EU439104.1 | EU439186.1 |
| *Hymenophyllum holochilum* | NC_039753.1 | NC_039753.1 | NC_039753.1 | — |
| *Marattia alata* | — | EU439082.1 | EU439108.1 | EU439190.1 |
| *Marattia douglasii* | EU439061.1 | EU439083.1 | EU439109.1 | EU439191.1 |
| *Marattia laxa* | EU439062.1 | EU439084.1 | EU439111.1 | EU439193.1 |
| *Matonia pectinata* | EU352280.1 | EU352307.1 | AY612666.1 | — |
| *Osmunda regalis* | — | EF588706.1 | EF588771.1 | — |
| *Osmundastrum cinnamomeum* | AF313539.1 | EF588711.1 | — | — |
| *Ptisana attenuata* | — | AF313581.1 | EU439125.1 | EU439206.1 |
| *Ptisana fraxinea* | EU439067.1 | EU439088.1 | EU439131.1 | EU439212.1 |
| *Ptisana melanesica* | — | EU439090.1 | EU439134.1 | EU439214.1 |
| *Ptisana mertensiana* | — | — | EU439120.1 | EU439201.1 |
| *Ptisana oreades* | — | EU439087.1 | EU439130.1 | EU439211.1 |
| *Ptisana pellucida* | — | — | EU439121.1 | EU439202.1 |
| *Ptisana purpurascens* | — | EU439089.1 | EU439132.1 | EU439213.1 |
| *Ptisana salicifolia* | — | — | EU439133.1 | — |
| *Ptisana squamosa* | — | — | EU439119.1 | EU439200.1 |
| *Ptisana sylvatica* | — | — | EU439117.1 | EU439198.1 |
| *Saccoloma inaequale* | MK705756.1 | MK705756.1 | MK705756.1 | — |
| *Schizaea elegans* | NC_035807.1 | NC_035807.1 | NC_035807.1 | — |

**Table S.2: Fossil Accesions**

| Taxon | Age Range | Geologic and Stratigraphic Information | Basis for Numerical Age Determition | References |
|---|---|---|---|---|
| *Acaulangium bulbaceum* | [303.7 – 307] | Upper Pennsylvanian Callhoun Coal; Missourian series of the Matoon Formation, McLeansboro Group | The Missourian North American stage corresponds to the Kasimovian stage of the ICC. | Millay (1977); Jacobson (2006) |
| *Angiopteris lygodiifolia* | [0 – 0] | Living | — | — |
| *Angiopteris evecta* | [0 – 0] | Living | — | — |
| *Angiopteris itoi* | [0 – 0] | Living | — | — |
| *Angiopteris smithii* | [0 – 0] | Living | — | — |
| *Angiopteris boninensis* | [0 – 0] | Living | — | — |
| *Angiopteris tonkinensis* | [0 – 0] | Living | — | — |
| *Araiangium pygmaeum* | [303.7 – 307] | Upper Pennsylvanian Callhoun Coal; Missourian series of the Matoon Formation, McLeansboro Group | The Missourian North American stage corresponds to the Kasimovian stage of the ICC. | Millay (1982); Jacobson (2006) |
| *Botryopteris tridentata* | [307 – 315.2] | Occurs in coal balls from Upper Carboniferous coal swamp deposits from the Westphalian of Germany and North America. The whole-plant description used here comes from coal balls from the middle Pennsylvanian of southeastern Kansas. | Middle Pennsylvanian of the ICC. | Rothwell and Good (2000) |
| *Buritiranopteris costata* | [259.1 – 298.9] | The reconstructed fossils come from the middle part of the Balsas Group of the Paríba Basin, in the Araguaí-Filadélfia region of Brasil. Either in the Motuca Formation (which Tavares et al. (2014) suggested to be early Permian in age) or the Pedro de Fogo Formation (see Arauja et al. 2016). Occurrences in Germany and France are restricted to the Asselian-Sakmarian (early Permian). Based on these sources we use a range from the Early to Middle Permian (Asselian to end of the Capitanian). | Asselian to end of the Capitanian of the ICC. | Tavares et al. (2014); Araújo et al. (2016) |
| *Christensenia aesculifolia* | [0 – 0] | Living | — | — |
| *Convexocarpus distichus* | [272.95 – 283.5] | Lower Permian, Kungurian Stage of the middle Fore Urals | Kungurian Stage of the ICC. | Naugolnykh (2013) |
| *Corynepteris involucrata* | [305.9 – 313.8] | Desmoinesian Stage of the Cabaniss Formation, Cherokee Group | Desmoinesian Stage is 305.9 – 313.8 per FossilWorks, approximately Middle Pennsylvanian | Baxter and Baxendale (1976) |
| *Cyathea multiiflora* | [0 – 0] | Living | — | — |
| *Daea elliptica* | [0 – 0] | Living | — | — |
| *Daea grandifolia* | [0 – 0] | Living | — | — |
| *Daea leprieurii* | [0 – 0] | Living | — | — |
| *Daea nodosa* | [0 – 0] | Living | — | — |
| *Daeites rigida* | [295 – 298.9] | Taiyuan Formation; Northwest Chi Floral Province; Longshou Mt., Xizang Autonomous Region, Chi Asselian, earliest Permian | Asselian stage of the ICC. | Shen (1995); Shen (1995) |
| *Daeopsis fecunda* | [201.3 – 208.5] | Kustatscher et al. (2012) indicate that this species only occurs in the Rhaetian (late Triassic), but is distibuted in Europe, Asia, and possibly Argentina. | Rhaetian stage of the ICC. | Halle (1921); see also Kustatscher et al. (2012) |

| | | | | |
|---|---|---|---|---|
| *Diplopterygium glaucum* | [0 – 0] | Living | — | — |
| *Dipteris conjugata* | [0 – 0] | Living | — | — |
| *Eoangiopteris goodii* | [295 – 303.7] | Late Pennsylvanian; From the lower portion of the Monongahela Series in Ohio, from a coal seam equivalent to either the Pittsburgh (No. 8) or Redstone (No. 8A) Coals. | The lower part of the Monongahela group corresponds to the Virgilian stage (295 – 303.7 per FossilWorks) (see Nadon et al. 1998) | Nadon et al. (1998); Millay (1978) |
| *Escapia christensenioides* | [132.9 – 139.8] | Longarm Formation equivalent, Valanginian, Early Cretaceous, from Apple Bay (Vancouver Island) | Valanginian of the ICC. | Rothwell et al. (2018a) |
| *Eupodium kaulfussii* | [0 – 0] | Living | — | — |
| *Eupodium laeve* | [0 – 0] | Living | — | — |
| *Gemellitheca saudica* | [251.9 – 272.95] | Description based on material from two localities: Gomaniimbrik Formation (Capitanian to Changhsingian stages; Stolle 2007; Baud et al. 2016) of the Hazro inlier in southeastern Turkey and the Uyzah plant bed of central Saudi Arabia (Kazanian, early Late Permian; Senalp and Al-Duaiji 2001). | Kazanian corresponds to the Roadian stage of the ICC (FossilWorks), total range is Roadian to Changhsingian stages of the ICC. | Baud et al. (2016); Senalp and Al-Duaiji (2001); Stolle (2007); Wagner et al. (1985) |
| *Grammatopteris freitasii* | [259.1 – 298.9] | From the Pedro de Fogo Formation in the Maranhao Basin, NE Brazil. Considered to be the upper part of the Pedra de Fogo Formation by Araújo et al. (2016), from the exposures of the Araguaí-Filadélfia region of the Middle Permian (Rodian through Capitanian). These deposits have been ascibed to the Early Permian by other authors, thus a more conservative estimate ranges Early to Middle Permian (259.1 – 298.9). (see also *Buritiranopteris costata*) | Asselian to end of the Capitanian stages of the ICC. | Rößler and Galtier (2002); Rössler and Noll (2002); Araújo et al. (2016) |
| *Grandeuryella reultii* | [303.4 – 305.9] | From Poudingue Mosaique at Grand Croix, in the St. Etienne Basin of central France. Specimens come from a conglomerate that is at the base of the Stephanian B, but is reworked and contains material that is from Stephanian A. | Stephanian A is equivalent to the Khamovnichean, which is part of the Kasimovian (FossilWorks age 304.8 – 305.9), Stephanian B upper boundary given by the Dorogovilovian, which is still part of the Kasimovian (FossilWorks). | Weiss (1885); Lesnikowska and Galtier (1992) |
| *Hopetedia praetermissa* | [227 – 237] | Late Triassic Pekin Formation of North Carolina; from the middle portion of the formation in the Boren Clay Company pit near Gulf; said to be late Carnian | Carnian stage of the ICC. | Axsmith et al. (2001) |
| *Hymenophyllum holochilum* | [0 – 0] | Living | — | — |
| *Marattiopsis aganzhenensis* | [174.1 – 201.3] | Lower Jurassic Daxigou Formation of Lanzhou, Gansu, Chi. The formation consists of three members, the plants are collected from the upper one, but there is no dating to constrain the members further within the Lower (”Early”) Jurassic. | Lower Jurassic of the ICC. | Yang et al. (2008) |
| *Marattia alata* | [0 – 0] | Living | — | — |
| *Marattiopsis anglica* | [166.1 – 174.1] | Yorkshire Jurassic; the formations Saltwick to Scalby span part of the Aalenian to the end of the Bathonian (Livera and Leeder 1981) | Aalenian to Bathonian stages of the ICC. | Livera and Leeder (1981); Van Konijnenburg-Van Cittert (1975) |
| *Marattiopsis asiatica* | [174.1 – 201.3] | Described from the Lower Jurassic Hsiangchi Formation in Zigui, Hubei Province, Chi. | Lower Jurassic of the ICC. | Wang (1999) |

| | | | | |
|---|---|---|---|---|
| *Marattia douglasii* | [0 – 0] | Living | — | — |
| *Marattia excavata* | [0 – 0] | Living | — | — |
| *Marattia laxa* | [0 – 0] | Living | — | — |
| *Marattiaceae indet Vera 2016* | [113 – 125] | Specimens are synangia from the Lower Cretaceous (Aptian) of Antarctica, Cerro Negro Formation, Byers Group. | Aptian stage of the ICC. | Vera and Césari (2016) |
| *Marattiopsis crenulatus* | [201.3 – 208.5] | Triassic (Rhaetian) of Sweden | Rhaetian stage of the ICC. | Lundblad (1950) |
| *Marattiopsis patagonica* | [177.27 – 189.036] | Cerro Bayo Locality, Lonco Trapial Formation; Early Jurassic age (most likely Pliensbachian). | Ages constrained from U-Pb dating of ash layers above and below the Lonco Trapial formation, from Cuneo et al 2013. | Escapa et al. (2015); Cúneo et al. (2013) |
| *Marattiopsis vodrazkae* | [86.3 – 89.8] | Coniacian of the Hidden Lake Formation, James Ross Island, Antarctica | Coniacian of the ICC. | Kvaček (2014) |
| *Matonia pectita* | [0 – 0] | Living | — | — |
| *Millaya tularosa* | [280 – 298.9] | Lower Permian (Wolfcampian) Bursum Formation; clay shale pits in Tularosa, New Mexico. | North American Wolfcampian is 298.9 – 280 per GeoWhen. | Mapes and Schabilion (1979) |
| *Osmunda regalis* | [0 – 0] | Living | — | — |
| *Osmundastrum cinmomeum* | [0 – 0] | Living | — | — |
| *Pekinopteris auriculata* | [201.3 – 237] | Pekin Formation of the Newark Group, central North Carolina, Upper Triassic. | Late Triassic of the ICC. | Hope and Patterson III (1970); Delevoryas and Hope (1978) |
| *Pertica quadrifaria* | [387.7 – 407.6] | Described as being from the Trout Valley Formation (Emsian?) and revised as a member of the Tomhegan formation (Osberg et al. 1985, Maine Geological Survey). Gastaldo (2016) regards the Trout Valley Formation as being between the late Emsian (Lower Devonian) to early Eifelian (Middle Devonian) | Range provided is the Emsian to Eifelian stages of the ICC. | Kasper and Andrews (1972); Gastaldo (2016); Osberg et al. 1985 |
| *Psilophyton crenulatum* | [393.3 – 407.6] | Early Devonian (early to middle Esmian), possibly the Dalhousie Group | Emsian stage of the ICC. | Doran (1980) |
| *Ptisa attenuata* | [0 – 0] | Living | — | — |
| *Ptisa fraxinea* | [0 – 0] | Living | — | — |
| *Ptisa melanesica* | [0 – 0] | Living | — | — |
| *Ptisa mertensia* | [0 – 0] | Living | — | — |
| *Ptisa oreades* | [0 – 0] | Living | — | — |
| *Ptisa pellucida* | [0 – 0] | Living | — | — |
| *Ptisa purpurascens* | [0 – 0] | Living | — | — |
| *Ptisa squamosa* | [0 – 0] | Living | — | — |
| *Ptisa sylvatica* | [0 – 0] | Living | — | — |
| *Qasimia schyfsmae* | [268.8 – 272.95] | Late Permian plant bed at Uyzah in central Saudi Arabia; Kazanian, early Late Permian (Senalp and Al-Duaiji 2001). | Kazanian corresponds to the Roadian stage of the ICC (FossilWorks). | Hill et al. (1985) |
| *Radstockia kidstonii* | [306.95 – 311.45] | Middle Pennsylvanian, from the Mazon Creek locality,Francis Creek Shale, Carbondale Formation of the Kewanee Group. | The Westaphalian D is 306.95 – 311.45 per FossilWorks. | Clements et al. (2019); Taylor (1967) |
| *Rhacophyton ceratangium* | [364.7 – 370.6] | Upper Devonian Hampshire Formation, West Virginia, which is Cassadagan to earliest Bradfordlan, equivalent to the upper middle Famennian (Fa2c). | Middle Famennian is part of the Famennian stage of the ICC, age is 364.7 – 370.6 per FossilWorks. | Dittrich et al. (1983); Cornet et al. (1976) |
| *Rothwellopteris pecopteroides* | [251.9 – 259.1] | From the Xuanwei Formation in the Panxian mining district of western Guizhou Province, southwest Chi; Lopingian epoch, late Permian. | Lopingian epoch of the ICC. | He et al. (2019) |

| | | | | |
|---|---|---|---|---|
| *Saccoloma iequale* | [0 – 0] | Living | — | — |
| *Schizaea elegans* | [0 – 0] | Living | — | — |
| *Scolecopteris alta A* | [315.2 – 323.2] | Larkian Series; Shore, Lancashire, England, UK (Westaphalian A; Watson 1906) and Breathitt Formation, "Lewis Creek," Leslie County, KY, USA (lower Middle Pennsylvanian; Millay 1982) | Wesphalian A substage corresponds to the Bashkirian stage of the ICC. | Watson (1906); Millay (1982) |
| *Scolecopteris antarctica L* | [237 – 247.2] | Fremouw Peak locality in the central Transantarctic Mountains; fossils are from the base of the upper part of the Fremouw Fm., which is middle Triassic based on vertebrate fossils and palynology (see also Collinson et al. 2006). | Middle Triassic of the ICC. | Collinson et al. (2006); Delevoryas et al. (1992) |
| *Scolecopteris calicifolia L* | [307 – 315.2] | Middle Pennsylvanian Cherokee Group, Des Moines series; West Mineral, Kansas. | Middle Pennsylvanian of the ICC. | Millay (1979) |
| *Scolecopteris charma O* | [298.9 – 307] | Late Pennsylvanian Conemaugh Group, Duquesne Coal; Steubenville, Ohio. | Upper Pennsylvanian of the ICC. | Lesnikowska and Millay (1985) |
| *Scolecopteris dispora* | [307 – 315.2] | Early or Late Bolsovian, Pennsylvanian; Clifland Coal Member; Schuler Mine, India. | Bolsovian corresponds to being within the Moscovian stage of the ICC. | Lesnikowska and Willard (1997) |
| *Scolecopteris fragilis L* | [307 – 315.2] | Middle Pennsylvanian Cherokee Group, Des Moines Series; "What Cheer," Keokuk County, India | Middle Pennsylvanian of the ICC. | Millay (1979) |
| *Scolecopteris guizhouensis* | [251.9 – 254.14] | Uppermost Permian Wangjiazhai Formation in Guizhou Province, south-western Chi, corresponding to the Changhsingian stage of the Permian. | Changhsingian stage of the ICC. | He et al. (2006) |
| *Scolecopteris illinoensis* | [298.9 – 307] | Late Pennsylvanian coal balls of the Mattoon Formation, McLeansboro Group; "Berryville," Sumner, Illinois | Upper Pennsylvanian of the ICC. | Millay (1979) |
| *Scolecopteris incisifolia L* | [307 – 315.2] | Middle Pennsylvanian lower to middle Cherokee Group, Des Moines Series, Urbandale Mine, India | Middle Pennsylvanian of the ICC. | Mamay (1950) |
| *Scolecopteris iowensis O* | [307 – 315.2] | Late Pennsylvanian coal balls of theMattoon Formation, McLeansboro Group; "Berryville," Sumner+D76:D77, Illinois | Middle Pennsylvanian of the ICC. | Mamay (1950) |
| *Scolecopteris latifolia L* | [298.9 – 307] | Late Pennsylvanian coal balls of the Mattoon Formation, McLeansboro Group, Calhoun Coal; Calhoun coal mine, Illinois | Upper Pennsylvanian of the ICC. | Millay (1979) |
| *Scolecopteris majopsis O* | [307 – 315.2] | Middle Pennsylvanian Carbondale Formation of the Kewanee Group, Sahara Mine, Carrier Mills, Illinois | Middle Pennsylvanian of the ICC. | Millay (1979) |
| *Scolecopteris mamayi L* | [307 – 315.2] | Middle Pennsylvanian Lisman Formation, Allegheny Series; Providence, Kentucky | Middle Pennsylvanian of the ICC. | Millay (1979) |
| *Scolecopteris minor M* | [298.9 – 307] | Late Pennsylvanian McLeansboro Group, from the Danville #7 coal, Hegler mine, Danville, Illinois | Upper Pennsylvanian of the ICC. | Hoskins (1926) |
| *Scolecopteris monothrix L* | [298.9 – 307] | Late Pennsylvanian coal balls of the Mattoon Formation, McLeansboro Group; "Berryville," Sumner, Illinois | Upper Pennsylvanian of the ICC. | Ewart (1961) |

| | | | | |
|---|---|---|---|---|
| *Scolecopteris nigra A* | [307 – 315.2] | Middle Pennsylvanian Carbondale Formation of the Kewanee Group, Sahara Mine, Carrier Mills, Illinois | Middle Pennsylvanian of the ICC. | Millay (1979) |
| *Scolecopteris oliveri O* | [295 – 298.9] | From the Permian (Upper Autunian) Assise de Millery Formation, Autun Basin, Autun, Central France | The Autunian correlates with Permian Asselian stage of the ICC per FossilWorks. | Scott (1932) |
| *Scolecopteris parkerensis L* | [298.9 – 307] | Late Pennsyvlanian Patoka Formation, Missourian Provincial Series; "St. Wendel," Posey County, IN | Upper Pennsylvanian of the ICC. | Lesnikowska and Willard (1997) |
| *Scolecopteris parvifolia* | [298.9 – 307] | Late Pennsylvanian coal balls of the Mattoon Formation, McLeansboro Group; "Berryville," Sumner, Illinois | Upper Pennsylvanian of the ICC. | Millay (1979) |
| *Scolecopteris saharaensis M* | [307 – 315.2] | Middle Pennsylvanian Carbondale Formation of the Kewanee Group, Sahara Mine, Carrier Mills, Illinois | Middle Pennsylvanian of the ICC. | Millay (1979) |
| *Scolecopteris shadensis* | [298.9 – 307] | Late Pennsylvanian, from above the Connellsville Sandstone, Shade, Ohio | Upper Pennsylvanian of the ICC. | Stubblefield (1984) |
| *Scolecopteris shanxiensis* | [290.1 – 298.9] | Early Permian Taiyuan Formation, Taiyuan City, Shanxi Province, Chi. Taiyuan Formation in Huebei province is said to be equivalent to the Asselian-early Sakmarian stages of the Early Permian. | Asselian to Sakmarian stages of the ICC. | Wang (1999) |
| *Scolecopteris vallumii L* | [307 – 315.2] | Middle Pennsylvanian Carbondale Formation of the Kewanee Group, Sahara Mine, Carrier Mills, Illinois | Middle Pennsylvanian of the ICC. | Millay (1979) |
| *Szea sinensis* | [259.1 – 272.95] | From the late Early Permian near njing Chi, said to be approximately equivalent to sediments of Gaudalupian age in North America based on marine inverts. | Guadalupian stage of the ICC. | Zhaoqi and Taylor (1988) |

### S§2 Graphical Models

The Bayesian total-evidence model includes five separate components: 1) the substitution model; 2) the molecular clock model; 3) the morphological transition model; 4) the morphological clock model, and; 5) the tree model. For each of the first two components, we use only one model. For the remaining components, we perform analyses under several different possible models. We provide graphical model representations for each component in the following sections.

We represent our models as directed factor graphs using the `TikZ` library `BayesNet` (Dietz and Luttinen 2012). In this representation, free parameters (stochastic nodes) are represented as unfilled circles, data ("clamped" stochastic nodes) are represented as filled circles, and deterministic parameters (deterministic nodes) are represented as dashed circles. Edges represent dependencies between nodes. Black squares are "factors" that express either the probability distribution from which the descending random variable is drawn, or the functional relationship between the input nodes and the descending deterministic node.

In each of the following figures, we focus on the parameters that are specific to a particular model component. When a parameter depends on a value from another model component, we include a placeholder node from that model component that is meant to represent the entire module. These placeholder nodes are indicated with labels connected by dashed lines.

#### S§2.1 Substitution Model

##### S§2.1.1 GTR + I + Γ

We assume that each molecular data subset evolves under a GTR+I+Γ model, and that rates vary among the $k$ molecular data partitions according to the vector of relative-rate multipliers, $s$.

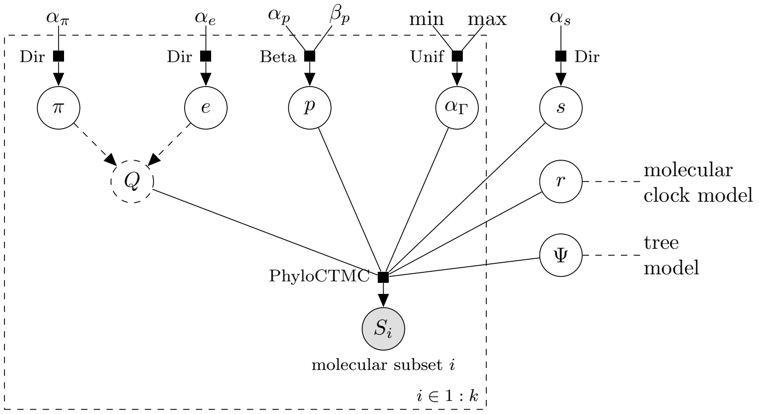


**Figure S2: The GTR+I+Γ substitution model.** The parameters are: 1) the stationary frequency, $\pi$; 2) the relative exchangeability rates, $e$; 3) the proportion of invariable sites, $p$; 4) the degree of among-site rate variation, $\alpha_\Gamma$, and; 5) the subset-specific rate multipliers, $s$.

#### S§2.2 Molecular Clock Model

##### S§2.2.1 Uncorrelated Lognormal Relaxed Clock

We assume the $(2n-2)$ lineage-specific rates of molecular evolution are drawn from a lognormal prior. We estimate the hyperparameters of the lognormal distribution from the data.

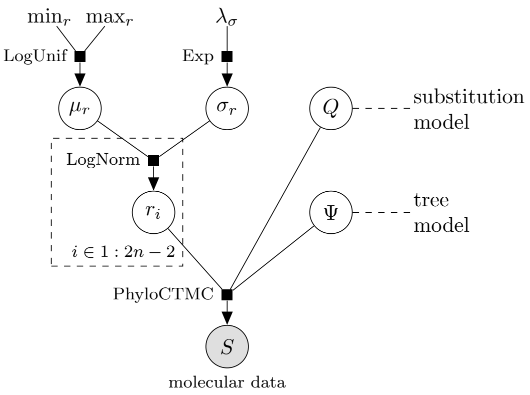


**Figure S3: The uncorrelated lognormal (UCLN) relaxed molecular clock model.** The parameters are: 1) the average rate of molecular evolution, $\mu_r$; 2) the degree of variation in rates of molecular evolution, $\sigma_r$, and; 3) the lineage-specific rates of evolution, $r_i$.

#### S§2.3 Morphological Transition Models

### S§2.3.1 Mk

The Mk model assumes that transitions among states occur at equal rates for the $l$ morphological partitions. We assume that rates vary among morphological data partitions according to the vector of relative-rate multipliers, $d$. Additionally, we assume that invariant characters are excluded from the dataset (not represented in the graphical model; Lewis 2001).

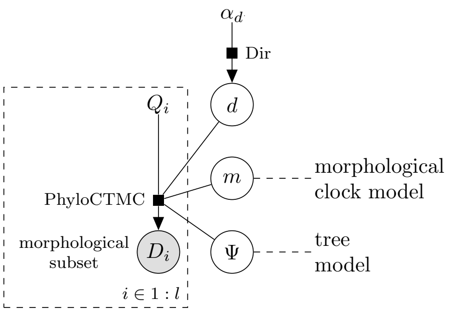


**Figure S4: The Mk morphological model.** The parameters are: 1) the partition-specific rate of evolution, $d$.

##### S§2.3.2 Mk $+\Gamma$

The Mk+Γ model assumes that transitions among states occur at equal rates for the $l$ morphological partitions. Additionally, we allow relative rates to vary among characters within a given data partition according to a mean-one Gamma distribution. We assume that rates vary among morphological data partitions according to the vector of relative-rate multipliers, $d$. Additionally, we assume that invariant characters are excluded from the dataset (not represented in the graphical model; Lewis 2001).

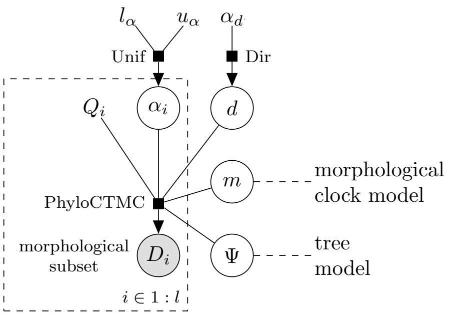


**Figure S5: The Mk+Γ morphological model.** The parameters are: 1) the partition-specific rate of evolution, $d$, and; 2) the degree of rate variation among characters, $\alpha_i$.

##### S§2.3.3 F81 Mixture

The F81 mixture model assumes that the morphological characters evolve toward a non-uniform stationary distribution, $\pi$. We allow the stationary distribution to vary among characters according to a discrete mixture model with $k$ categories. For binary characters, the $k$ categories are defined by a discrete Beta distribution with $k$ bins of equal probability mass. For multistate characters, the $k$ categories are drawn from a shared Dirichlet prior distribution and mixture weights, $\omega$. We assume that rates vary among morphological data partitions according to the vector of relative-rate multipliers, $d$. Additionally, we assume that invariant characters are excluded from the dataset (not represented in the graphical model; Lewis 2001).

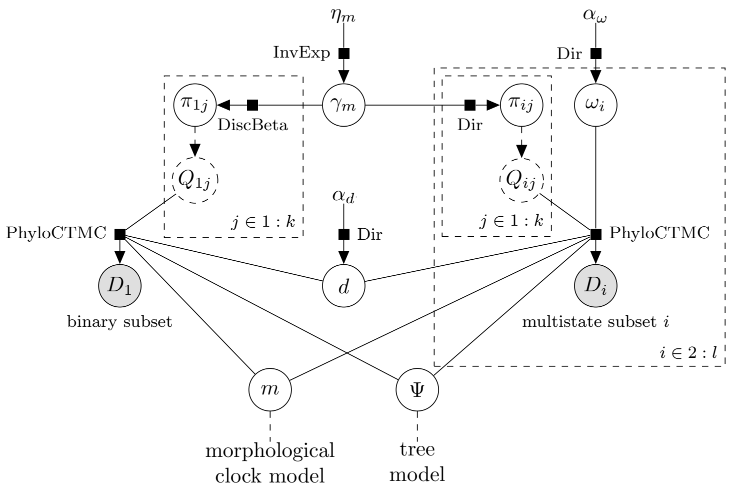


**Figure S6: The F81 mixture morphological model.** The parameters are: 1) the partition-specific rate of evolution, $d$; 2) the degree of variation among stationary frequencies, $\gamma_m$; 3) the partition-specific stationary frequencies, $\pi_{ij}$, and; 4) the mixture weight of multistate stationary frequencies, $\omega_i$.

##### S§2.3.4 F81 Mixture $+\Gamma$

The F81 mixture model assumes that the morphological characters evolve toward a non-uniform stationary distribution, $\pi$. We allow the stationary distribution to vary among characters according to a discrete mixture model with $k$ categories. For binary characters, the $k$ categories are defined by a discrete Beta distribution with $k$ bins of equal probability mass. For multistate characters, the $k$ categories are drawn from a shared Dirichlet prior distribution and mixture weights, $\omega$. Additionally, we allow relative rates to vary among characters within a given data partition according to a mean-one Gamma distribution. We assume that rates vary among morphological data partitions according to the vector of relative-rate multipliers, $d$. Additionally, we assume that invariant characters are excluded from the dataset (not represented in the graphical model; Lewis 2001).

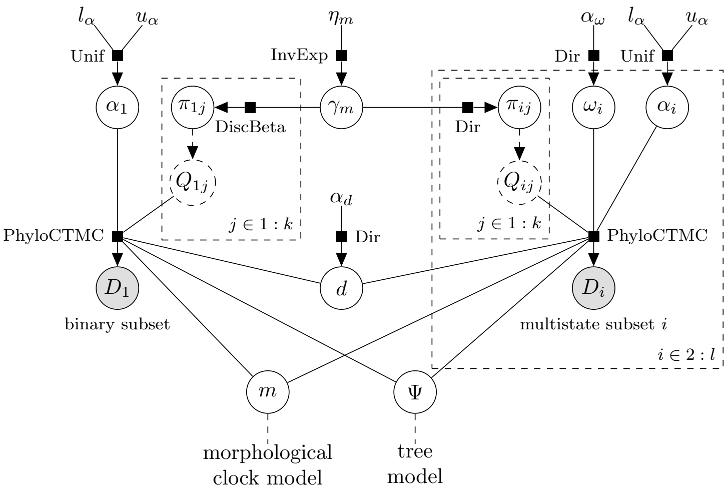


**Figure S7: The F81 mixture+$\Gamma$ morphological model.** The parameters are: 1) the partition-specific rate of evolution, $d$; 2) the degree of variation among stationary frequencies, $\gamma_m$; 3) the partition-specific stationary frequencies, $\pi_{ij}$; 4) the mixture weight of multistate stationary frequencies, $\omega_i$, and; 5) the degree of rate variation among characters, $\alpha_i$.

#### S§2.4 Morphological Clock Models

##### S§2.4.1 Linked

The linked morphological clock model assumes that lineage-specific rates of morphological evolution, $r_{m,i}$, are proportional to lineage-specific rates of molecular evolution. The parameter $\beta_m$ describes the relative rate of morphological to molecular evolution.

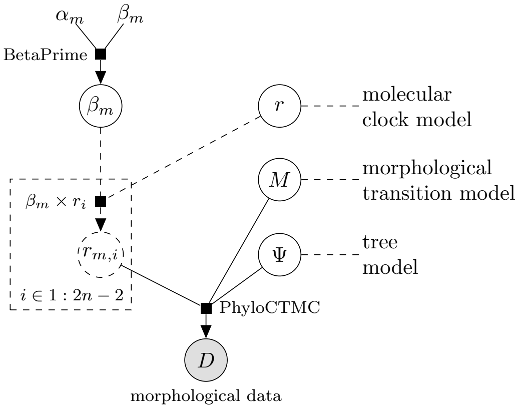


**Figure S8: The "linked" relaxed morphological model.** The parameters are: 1) the relative rate of morphological to molecular evolution, $\beta_m$, and; 2) the lineage-specific rate of morphological evolution, $m_i$.

##### S§2.4.2 Unlinked

The unlinked morphological clock model assumes that lineage-specific rates of morphological evolution are drawn from a lognormal prior distribution. The parameter $\beta_m$ describes the relative mean rate of morphological evolution to the mean rate of molecular evolution; the parameter $\sigma_m$ describes how much rates of morphological evolution vary among branches.

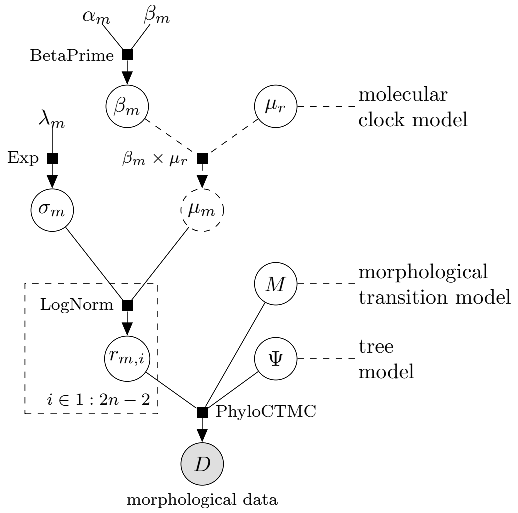


**Figure S9: The "unlinked" relaxed morphological model.** The parameters are: 1) the relative mean rate of morphological to molecular evolution, $\beta_m$; 2) the degree of variation in lineage-specific rates of morphological evolution, $\sigma_m$, and; 3) the lineage-specific rate of morphological evolution, $m_i$.

#### S§2.5 Tree Models

In all of the following tree models, we assume the process begins with the stem lineage with age $o$. Additionally, we accommodate uncertainty in the ages of fossil tips by requiring that the age of fossil tip $i$ in the tree $\Psi$ falls within the interval of uncertainty associated with the fossil. This is represented as an indicator function, $\mathbb{1}$, which has value 1 if the fossil tip is within the boundaries and 0 otherwise.

##### S§2.5.1 Uniform

The uniform tree prior model assumes that the ages of nodes are uniformly distributed *sensu* Ronquist et al. (2012).

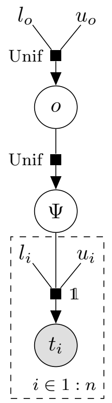


**Figure S10: The uniform tree model.** The parameters are: 1) the origin time (stem age) of the tree, $o$, 2) the phylogeny, $\Psi$, and; 3) the ages of the tips, $t_i$.

##### S§2.5.2 Constant-rate fossilized birth-death

The constant-rate fossilized birth-death model assumes that rates of fossilization, speciation, and extinction are constant through time and among lineages, and that extant species are sampled with probability $\rho$.

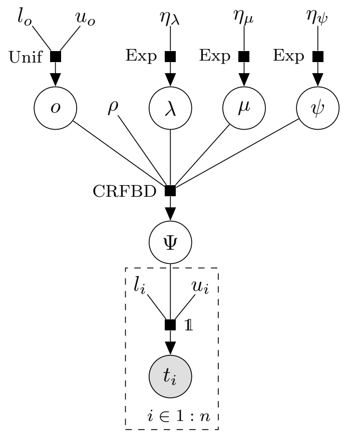


**Figure S11: The CRFBD tree model.** The parameters are: 1) the origin time (stem age) of the tree, $o$, 2) the fraction on extant species included in the dataset, $\rho$; 3) the rate of speciation, $\lambda$; 4) the rate of extinction, $\mu$; 5) the rate of fossilization, $\psi$; 6) the phylogeny, $\Psi$, and; 7) the ages of the tips, $t_i$.

##### S§2.5.3 Episodic fossilized birth-death with variable fossilization rates

This episodic fossilized birth-death model assumes that rates of speciation, and extinction are constant through time and among lineages, and that extant species are sampled with probability $\rho$. However, the fossilization rate is drawn from a mixture distribution with $m = 3$ categories. The rates and mixture weights of each category are free parameters.

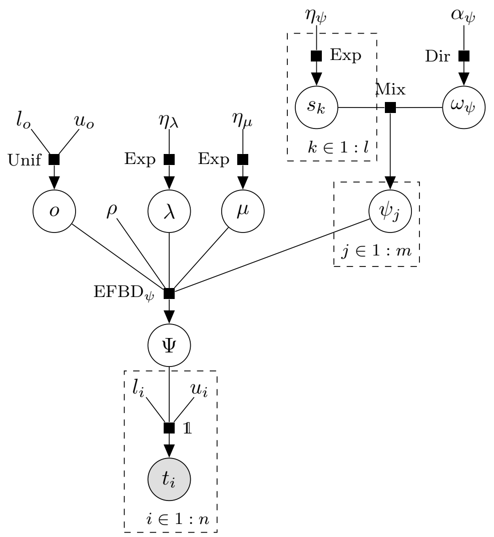


**Figure S12: The EFBD$_\psi$ tree model.** The parameters are: 1) the origin time (stem age) of the tree, $o$, 2) the fraction on extant species included in the dataset, $\rho$; 3) the rate of speciation, $\lambda$; 4) the rate of extinction, $\mu$; 5) the rates of the fossilization-rate mixture model, $s_k$; 6) the weights of the fossilization-rate mixture model, $\omega_\psi$; 7) the epoch-specific rates of fossilization, $\psi_i$; 8) the phylogeny, $\Psi$, and; 9) the ages of the tips, $t_i$.

##### S§2.5.4 Episodic fossilized birth-death with variable diversification rates

This episodic fossilized birth-death model assumes that rates of fossilization are constant through time and among lineages, and that extant species are sampled with probability $\rho$. However, the speciation and extinction rates are drawn from separate mixture distributions with $m = 3$ categories. The rates and mixture weights of each category are free parameters.

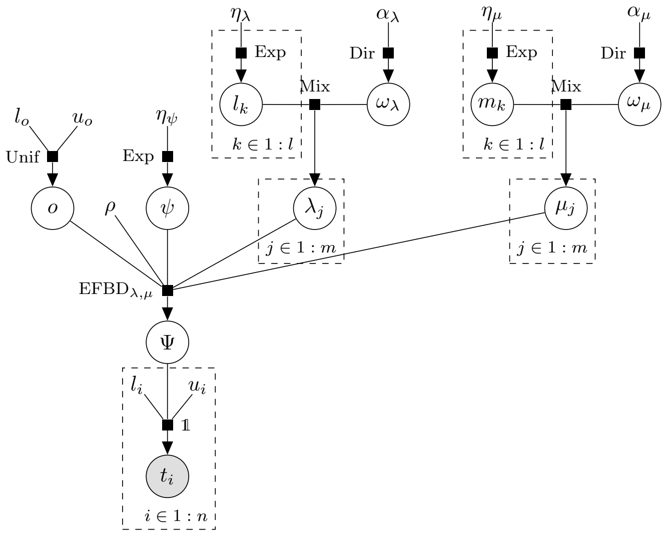


**Figure S13: The EFBD$_{\lambda,\mu}$ tree model.** The parameters are: 1) the origin time (stem age) of the tree, $o$, 2) the fraction on extant species included in the dataset, $\rho$; 3) the rates of the speciation-rate mixture model, $l_k$; 4) the weights of the speciation-rate mixture model, $\omega_\lambda$; 5) the epoch-specific rates of speciation, $\lambda_i$; 6) the rates of the extinction-rate mixture model, $m_k$; 7) the weights of the extinction-rate mixture model, $\omega_\mu$; 8) the epoch-specific rates of extinction, $\mu_i$; 9) the rate of fossilization, $\psi$; 10) the phylogeny, $\Psi$, and; 11) the ages of the tips, $t_i$.

##### S§2.5.5 Episodic fossilized birth-death with variable diversification and fossilization rates

This episodic fossilized birth-death model assumes that extant species are sampled with probability $\rho$. The fossilization, speciation and extinction rates are drawn from separate mixture distributions with $m = 3$ categories. The rates and mixture weights of each category are free parameters.

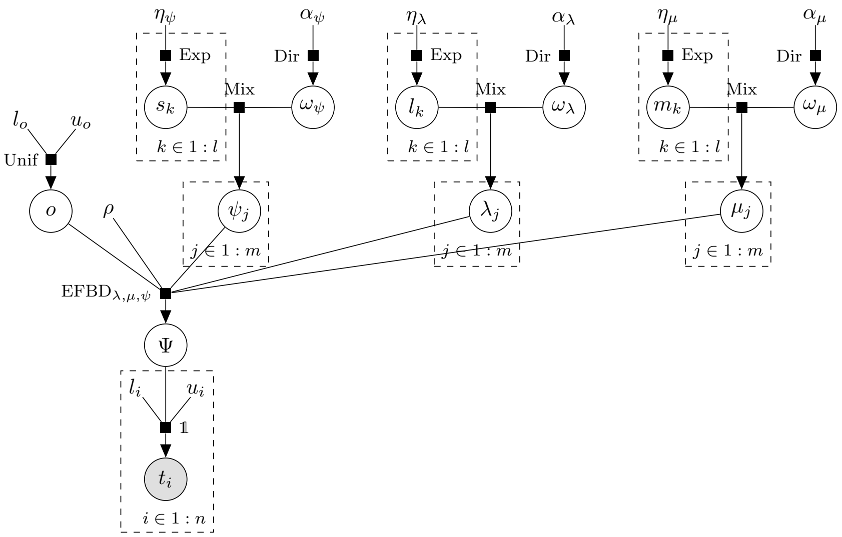


**Figure S14: The $\mathrm{EFBD}_{\lambda,\mu,\psi}$ tree model.** The parameters are: 1) the origin time (stem age) of the tree, $o$, 2) the fraction on extant species included in the dataset, $\rho$; 3) the rates of the speciation-rate mixture model, $l_k$; 4) the weights of the speciation-rate mixture model, $\omega_\lambda$; 5) the epoch-specific rates of speciation, $\lambda_i$; 6) the rates of the extinction-rate mixture model, $m_k$; 7) the weights of the extinction-rate mixture model, $\omega_\mu$; 8) the epoch-specific rates of extinction, $\mu_i$; 9) the rates of the fossilization-rate mixture model, $s_k$; 10) the weights of the fossilization-rate mixture model, $\omega_\psi$; 11) the epoch-specific rates of fossilization, $\psi_i$; 12) the phylogeny, $\Psi$, and; 13) the ages of the tips, $t_i$.

### S§3 MCMC Analyses

We performed all analyses in RevBayes, compiled from branch ssbdp_fix (commit ff39b2). In particular, we modified the episodic fossilized birth-death model (alias dnSSBDP) to use a user-defined starting tree, which significantly improved the initialization and convergence of the chain. However, the starting tree may also bias chains toward particular parts of the posterior distribution of trees, which could be an issue if the posterior distribution is rugged and mixing among trees is poor. To overcome this issue, we generated a marginal posterior distribution of trees under the uniform tree model (which does not require a user-defined starting tree), then initialized each independent chain under the FBD with a starting tree drawn randomly from the marginal posterior under the uniform model.

We refer readers to our Supplemental Archive at https://github.com/mikeryanmay/marattiales_supplemental/releases/tag/1.0 for specific details about chain lengths, proposal schemes, and prior specification.

#### S§3.1 MCMC Diagnosis

We diagnosed the convergence and mixing of each MCMC analysis, and ensured that each replicate analysis converged to the same joint posterior distribution. Owing to the large number of analyses, and the complexity of our models, we developed a semi-automated pipeline using custom R scripts (R Core Team 2019). This pipeline is available in our Supplementary Archive at https://github.com/mikeryanmay/marattiales_supplemental/releases/tag/1.0.

First, we determined the optimal burnin for each chain according to an ESS criterion. We computed the ESS for each continuous parameter using the R package coda (Plummer et al. 2006). To compute the ESS of the sampled phylogeny, we first computed the Robinson-Foulds and Kühner-Felsenstein distance from each sampled tree to the maximum clade-credibility (MCC) and maximum *a posterior* (MAP) tree for the analysis using the R package phangorn (Schliep et al. 2017). We then computed the ESS of these distance scores, as suggested by Warren et al. (2017); these ESS values reflect how well the chain mixes over tree topologies (the RF distance), and the tree topologies and branch lengths (the KF distance). To determine the optimal burnin fraction, we computed the ESS as described above at each burnin fraction from 1% to 95% in increments of 0.1%. For each increment, we computed the harmonic mean ESS among continuous parameters, and chose the burnin that yielded the highest harmonic mean ESS. (We used the harmonic mean as it is more sensitive to low values than the arithimetic mean, and therefore is a better reflection of the worst-behaving parameters in the chain.) Likewise, we found the burnin fraction that yields the highest ESS for the tree distance metrics by computing their harmonic mean. We then used the larger of these two burnin fractions to discard pre-burnin samples (for both the continuous-parameter and tree samples). Note: Unsurprisingly, in all cases the burnin fraction for the tree samples was much larger than for the continuous parameters. We therefore focused our assessment of multichain convergence on the tree samples.

Next, we assessed whether replicate chains converged to the same posterior distribution of trees. We used multidimensional scaling (MDS) to create plots of sampled tree space among the post-burnin samples from replicate chains (Hillis et al. 2005; Warren et al. 2017). We computed MDS plots of pairwise Robinson-Foulds and Kühner-Felsenstein distances within and among chains using the R package smacof (de Leeuw and Mair 2009). We visually assessed whether each chain sampled from the same part of tree space for each metric.

If a chain failed to achieve a harmonic-mean average ESS of at least 200, or if MDS plots indicated that some or all of the runs failed to converge, we deemed the run(s) failed and re-ran the analysis (or analyses). We repeated this procedure until we had at least four replicate chains that had sufficient

ESS and that demonstrated multichain convergence. We then combined the samples from the four replicate chains for all downstream analyses.

#### S§3.2 Computational Details

We performed all of our analyses on the University of California, Berkeley HPC cluster, `savio`, with compute nodes consisting of 20 Intel Xeon E5-2670 v2 processors and 64 GB of RAM. We used `openmpi` and `rb-mpi` to parallelize the individual chains of each MCMCMC analysis; nevertheless, each analysis requires approximately 70 hours (depending on the specific model) to converge and mix adequately. For our main analyses, a naive estimate of the total compute time would be 72 (the number of model combinations) $\times 5$ (the number of processors per analysis) $\times 4$ (the number of replicates per analysis) $\times 70$ (the number of hours per analysis) $\approx 100,000$ total compute hours. However, including failed MCMCs and secondary analyses, our study consumed $\approx 300,000$ hours of compute time: $\approx 200,000$ hours for the MCMCMC analyses and $\approx 100,000$ hours for the power-posterior analyses.

### S§4 Posterior-Predictive Simulation

We used posterior-predictive simulation to assess the adequacy of different models. Owing to the complexity of the morphological mixture model, patterns of missing morphological data, and acquisition bias (only sampling variable characters), we could not use existing simulators to perform these analyses. We therefore developed our own simulation protocol, which we provide in our Supplementary Archive. These scripts depend heavily on the `R` packages `ape` and `phytools` (Paradis and Schliep 2018; Revell 2012) for computing conditional likelihoods and simulating characters.

A given MCMC sample, $i$, includes a phylogeny $\Psi_i$, a vector of morphological branch rates, $m_i$, and a vector of morphological model parameters, $\theta_i$. To simulate a new dataset for sample $i$, we simulate each of the $c$ morphological characters. To simulate character $j$, we first determine the character-specific rate multiplier (if there is among-character rate variation) and character-specific rate matrix (if there is among-character matrix variation). We denote the conditional probability of character $D_j$, given that it has character-specific rate $r_k$ and character-specific matrix $Q_l$, as $P(D_j \mid r_k, Q_l, \Psi_i, \theta_i)$. We sample a character-specific rate and character-specific matrix from their joint posterior distribution:

$$P(r_k, Q_l \mid \Psi, \theta_i, D_j) \propto P(D_j \mid r_k, Q_l, \Psi_i, \theta_i) P(r_k) P(Q_l),$$

where $P(r_k)$ and $P(Q_l)$ are the prior probabilities of the character-specific rates and the character-specific matrices. In the among-character rate-variation models that we use (*i.e.*, discretized Gamma models), $P(r_k)$ is uniform among rate categories, as each category has equal cumulative probability under the Gamma distribution. The prior probability of the character-specific matrices are determined by the appropriate mixture-weight parameter, $\omega$ (as depicted in S6). After sampling the character-specific rate and matrix, we simulate a new character on the tree $\Psi_i$ with branch-specific rate multipliers $m_i$. We then remove simulated data according to the pattern of missing data for character $D_j$. If the remaining characters are invariant, we simulate a new character until the character is not invariant.

We repeat the above procedure for each character in the morphological matrix. We then compute the parsimony score for each simulated character, $p_{i,j}^{\text{sim}}$, and for each observed character, $p_{i,j}^{\text{obs}}$, given the tree $\Psi_i$. We compute the discrepency statistics $S_i$ and $V_i$ as:

$$S_i = \sum_j p_{i,j}^{\text{sim}} - p_{i,j}^{\text{obs}},$$

and

$$V_i = \left[ \frac{1}{j} \sum_j \left( p_{i,j}^{\text{sim}} \right)^2 - \left( \frac{1}{j} \sum_j p_{i,j}^{\text{sim}} \right)^2 \right] - \left[ \frac{1}{j} \sum_j \left( p_{i,j}^{\text{obs}} \right)^2 - \left( \frac{1}{j} \sum_j p_{i,j}^{\text{obs}} \right)^2 \right]$$

The statistic $S_i$, the sum of the difference in parsimony score among characters, is intended to capture the ability of the model to describe overall rates of evolution. The statistic $V_i$, the variance in the difference in parsimony scores among characters, is intended to capture whether the model adequately describes how rates vary among characters.

We repeat this procedure for each of $r$ samples from the posterior distribution. For a given posterior distribution, this generates a posterior-predictive distribution of $S$ and $V$. If these posterior-predictive distributions do not include 0 in their 95% predictive interval, we consider the model to the inadequate (it is unable to describe overall rates of evolution or how rates vary among characters, or both).

### S§5 Empirical Considerations: Taxon Sample and Rooting Strategies

The goal of our study is to estimate the divergence times within our ingroup (the extant and extinct Marattiales). Presumably, this inference depends critically on our ability to identify the position and age of fossil lineages, and especially the position and age of the root (*i.e.*, the node separating the Marattiales and leptosporangiate ferns, assuming that they are inferred to be reciprocally monophyletic). However, our fossil dataset is dominated by a cluster of late Carboniferous taxa whose relationships to each other and to surviving lineages are apt to be highly uncertain, which may limit our ability to infer ancient divergences. Additionally, the inclusion of a sparsely sampled outgroup makes it difficult to specify an appropriate taxon-sampling fraction for the fossilized birth-death models.

A specific set of analyses, described here, were designed to understand the robustness of divergence-time estimates within the Marattiales to these empirical considerations. In all of the analyses that follow, we assume the best-performing morphological transition model, morphological clock model, and tree model, as determined in the main set of analyses.

#### S§5.1 Ancient plants

Our standard dataset includes only our focal taxon (the Marattiales) and its sister group (the leptosporangiate ferns). Consequently, there may only be minimal information available about the age of the root, and whether the ingroup and outgroup are reciprocally monophyletic. In most applications of phylogenetic inference, the monophyly of the ingroup is enforced by the rooting procedure, which is not the case here; instead, inference of the root position comes from the clock and tree models, and small differences in root position can result in fossils switching from one side of the tree to the other. Therefore, including older fossils that are outside of the Marattiales + leptosporangiate clade might have a strong effect on divergence-time estimates, both by imposing stronger constraints on the maximum age, and by helping to inform the position of the first split within the Marattiales + leptosporangiates clade (the root in the standard analysis). To investigate this possibility, we conducted analyses with three additional ancient land-plant fossils: 1) *Psilophyton crenulatum* from the early Devonian (Doran 1980); 2) *Pertica quadrifaria* from the early Devonian (Kasper and Andrews 1972), and; 3) *Rhacophyton ceratangium* from the late Devonian (Cornet et al. 1976; Dittrich et al. 1983.

#### S§5.2 Polarized rooting

A common alternative to using a more inclusive taxon set to identify the root is to specify hard topological constraints: effectively, placing a prior probability of 0 on trees where the ingroup is not monophyletic. However, given potential uncertainty about the topological positions of our fossils, hard topological constraints could be overly informative.

We explored a novel strategy to place the root, without adding taxa outside of the Marattiales + leptosporangiates clade, by imposing *a priori* assumptions about the states of some morphological characters at the root of the tree (effectively "polarizing" the state of the character). For characters that evolve relatively slowly, imposing an assumption on the root state effectively constrains the branches on which the root may be placed. However, this constraint is weaker than a hard topological constraint, as the probabilities of alternative rootings depend on the implied amount of morphological evolution, and no root position is ruled out *a priori*.

In this experiment, we polarized four binary characters based on external information about the state at the root. Specifically, we polarized three binary characters for which there is good reason to believe the ingroup has a derived state (char. 7: polyarch root stele, absent at the root; char. 21:

phloem maturation, exarch at the root, and; char. 50: sporangia developmentally attached forming synangium, absent at the root), and one binary character that is derived in the outgroup (char. 89: sporangium wall thickness, thick at the root; Rothwell et al. 2018a). We achieved polarization by setting the prior probability of the appropriate state at the root to 1 and the remaining states to 0 (as opposed to using the stationary frequency as the prior probability).

#### S§5.3 Ingroup-only

The fossilized birth-death models require specifying a single taxon-sampling fraction ($\rho$) that applies across the full tree. However, our standard dataset includes a well-sampled ingroup (the Marattiales) and a very sparsely sampled outgroup (the leptosporangiate ferns); the assumption that an extant sample from each lineage has an equal chance of being included in the dataset will be incorrect whether we assume the ingroup sampling fraction ($\rho = 27 \div 111$) or the overall sampling fraction ($\rho = 36 \div 12000$). To understand the impact of the assumed taxon-sampling fraction on estimates of divergence times, we conducted analyses with only ingroup taxa (extant and extinct Marattiaceae). In this case, the ingroup sampling fraction is a realistic representation of the actual extant-taxon sampling scheme, and may provide more reliable estimates of divergence times.

#### S§5.4 Results

The broad pattern and timing of divergences are similar among these experimental analyses. Nevertheless, there are some notable trends in the effect of the different approaches on inferred ages of specific clades. The divergence estimates for older splits are consistently similar in the ancient plants analysis (*i.e.*, adding both character and age data) and the standard analysis (Figs. S57, S58). Conversely, polarizing characters at the root (*i.e.*, adding character data but not age data directly) resulted in older divergence estimates than all other analyses, for both the deeper divergences and for the crown group (Figs. S57, S58), although the effect was increasingly pronounced for older clades (Fig. S59). The ingroup-only analysis resulted in similar divergence-time estimates to the standard taxon set and to the ancient plants analysis (Fig. S59), even though the ingroup-only analyses were topologically the most distinct (Fig. S57), and support values for the deeper nodes were particularly low.

Including an extra set of ancient plant fossils or rooting the tree by polarizing a few characters each resulted in increased support for the ingroup-outgroup split (0.43 and 0.85 posterior probability, respectively, compared to 0.33 with the standard dataset). In other words, as expected, these approaches improved our ability to estimate the location of the root. The polarized analysis also resulted in markedly improved MCMC mixing, but, surprisingly, inferred noticeably older ages for the crown group (Figs. S57, S59), possibly because polarized characters require fewer state changes near the root, leading to lower rates of evolution and therefore older ages near the present.

By contrast, the ancient plants dataset resulted in ages that were very consistent with the standard dataset (Fig. S59), suggesting that, in our case at least, the addition of both rooting and age information (via the inclusion of an additional outgroup "layer") was more effective than adding rooting information alone (via polarizing some characters).

The similarity between the age inferences from our ingroup-only analyses and those under the standard dataset (with the ingroup sampling fraction) was encouraging, and suggests that ages estimated from the full dataset under the ingroup sampling fraction are reliable, whereas those under the overall sampling fraction are probably overestimates. However, using only ingroup taxa also lead to increased topological uncertainty near the root of the tree (Fig. S55, S56). More generally, using alternative sampling fractions and taxon sets proved to be an effective means of validating divergence-time estimates, and may be useful for rooting other TED analyses and for evaluating the impact of violations to the assumptions of existing incomplete-sampling models.

### S§6 Extended Results

#### S§6.1 Ingroup Sampling Fraction

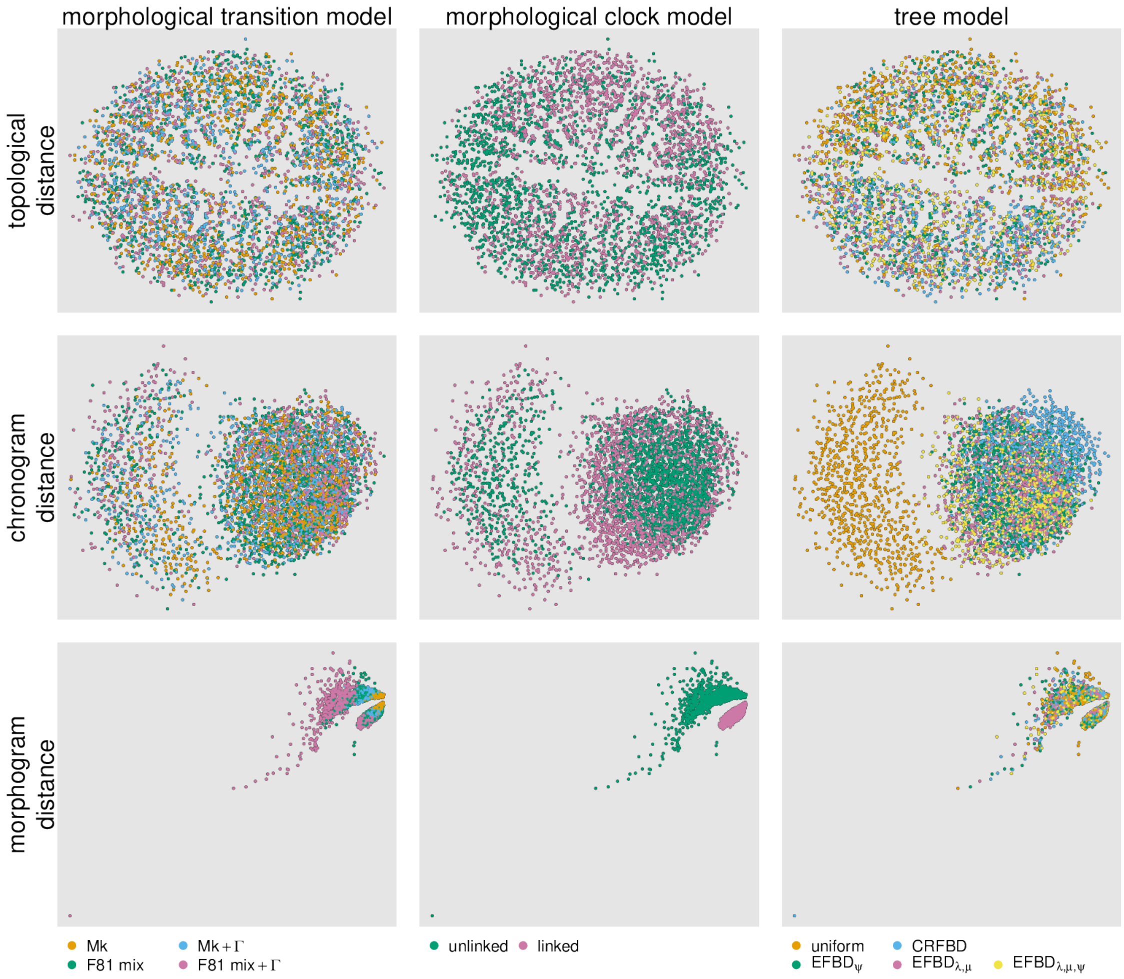


**Figure S15: Comparing distributions of molecular trees among model combinations.** We compute the Robinson-Foulds distance (RF, a measure of topological distance, top row), the Kühner-Felsenstein distance (KF, a distance metric that incorporates both topology and branch lengths) between morphological phylograms ("morphograms", middle row) and chronograms (bottom row). We removed taxa without molecular data before computing distances. We then plot the (square-root transformed) distances in two-dimensional space using multi-dimensional scaling (MDS); each point represents the location of a given sampled tree in tree space according to the distance metric. We color points according to the morpological transition model (column 1), the morphological clock model (column 2), or the tree model (column 3).

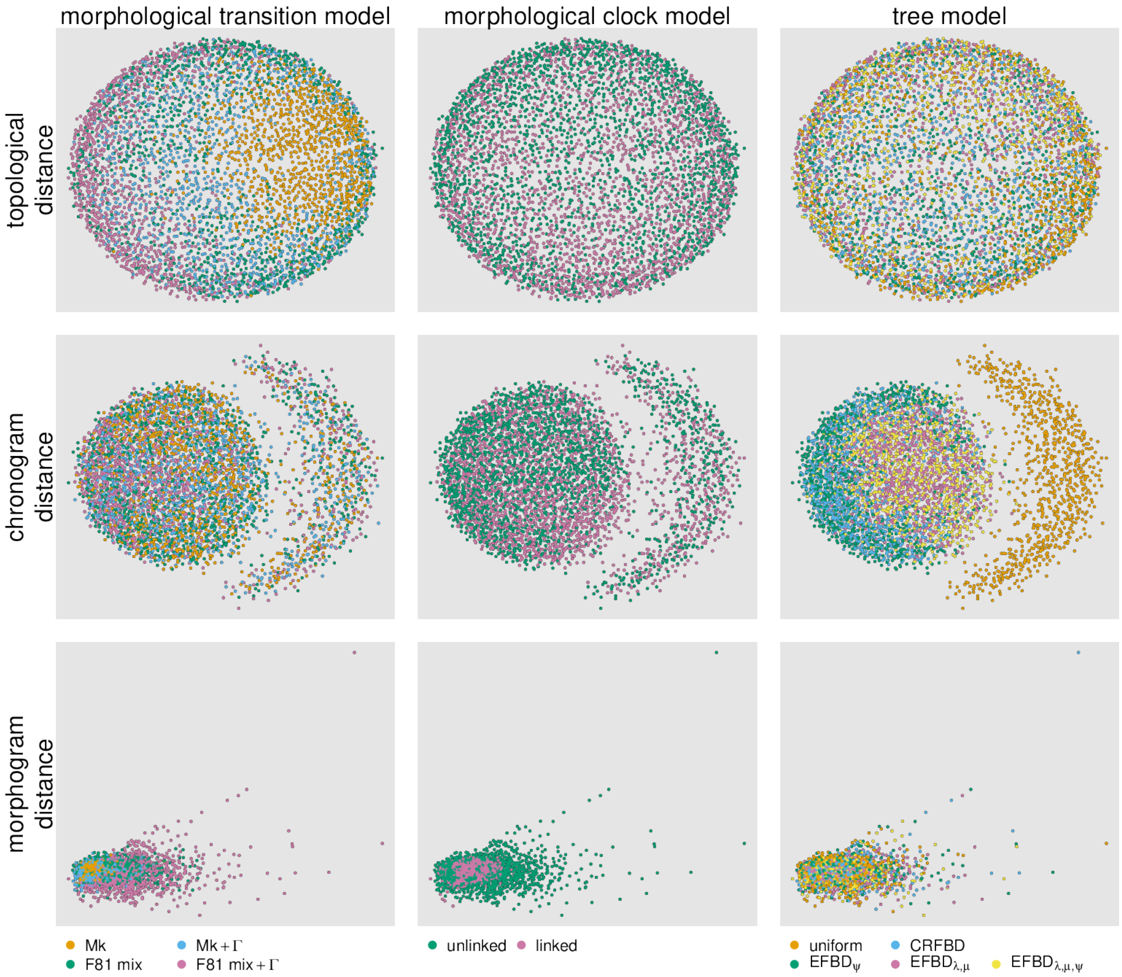


**Figure S16: Comparing distributions of extinct trees among model combinations.** We compute the Robinson-Foulds distance (RF, a measure of topological distance, top row), the Kühner-Felsenstein distance (KF, a distance metric that incorporates both topology and branch lengths) between morphological phylograms ("morphograms", middle row) and chronograms (bottom row). We removed extant taxa before computing distances. We then plot the (square-root transformed) distances in two-dimensional space using multi-dimensional scaling (MDS); each point represents the location of a given sampled tree in tree space according to the distance metric. We color points according to the morpological transition model (column 1), the morphological clock model (column 2), or the tree model (column 3).

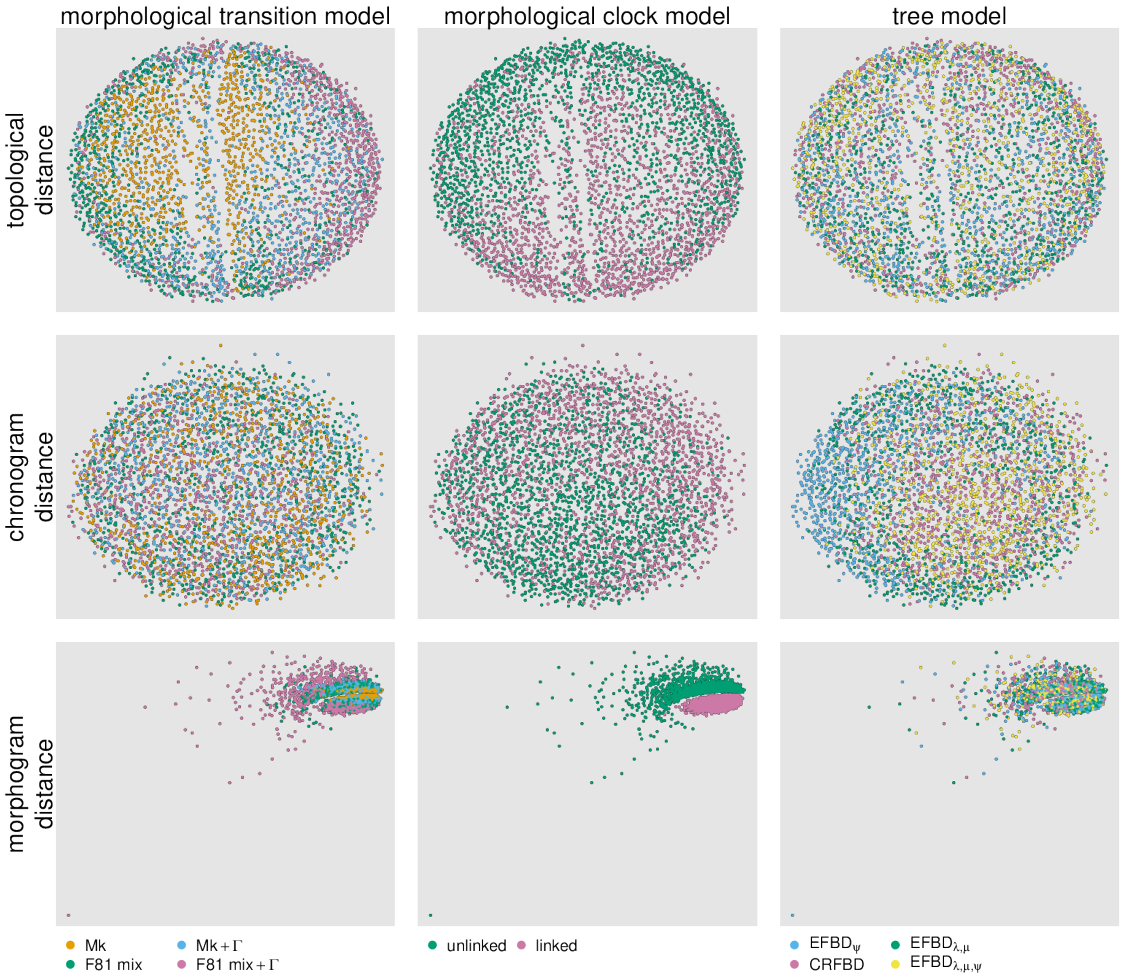


**Figure S17: Comparing distributions of trees among model combinations, excluding the uniform tree model.** We compute the Robinson-Foulds distance (RF, a measure of topological distance, top row), the Kühner-Felsenstein distance (KF, a distance metric that incorporates both topology and branch lengths) between morphological phylograms ("morphograms", middle row) and chronograms (bottom row). We removed extant taxa before computing distances. We then plot the (square-root transformed) distances in two-dimensional space using multi-dimensional scaling (MDS); each point represents the location of a given sampled tree in tree space according to the distance metric. We color points according to the morpological transition model (column 1), the morphological clock model (column 2), or the tree model (column 3).

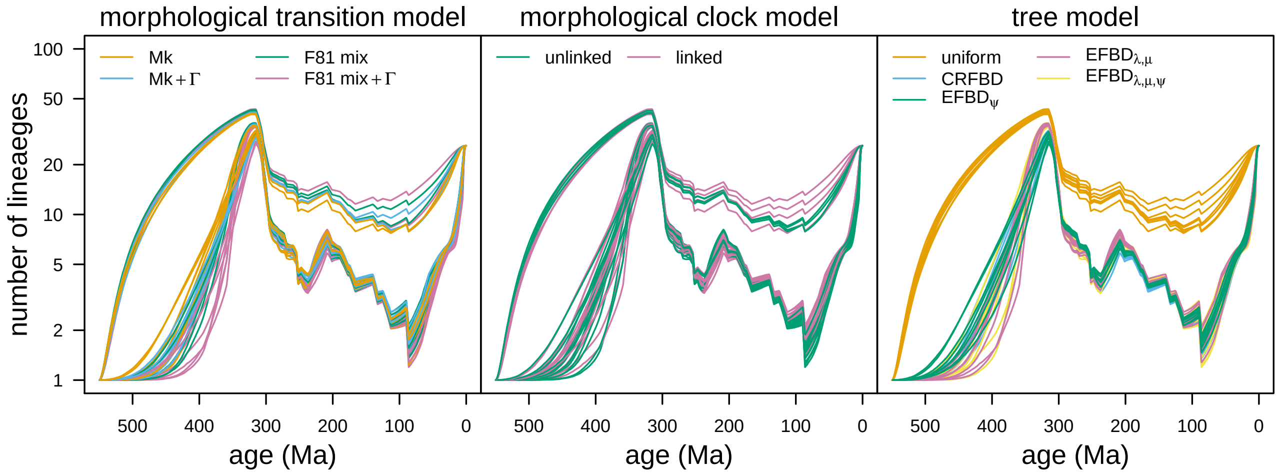


**Figure S18: Comparing lineage-through-time curves among model combinations.** Each curve corresponds to the lineage-through-time (LTT) curve for a given model, averaged over the posterior distribution of trees from that model. The curves in each panel are color coded by morphological-transition model (left), morphological-clock model (middle), and tree model (right), respectively. We removed outgroup taxa to emphasize the influence of model specification on age estimates within our ingroup.

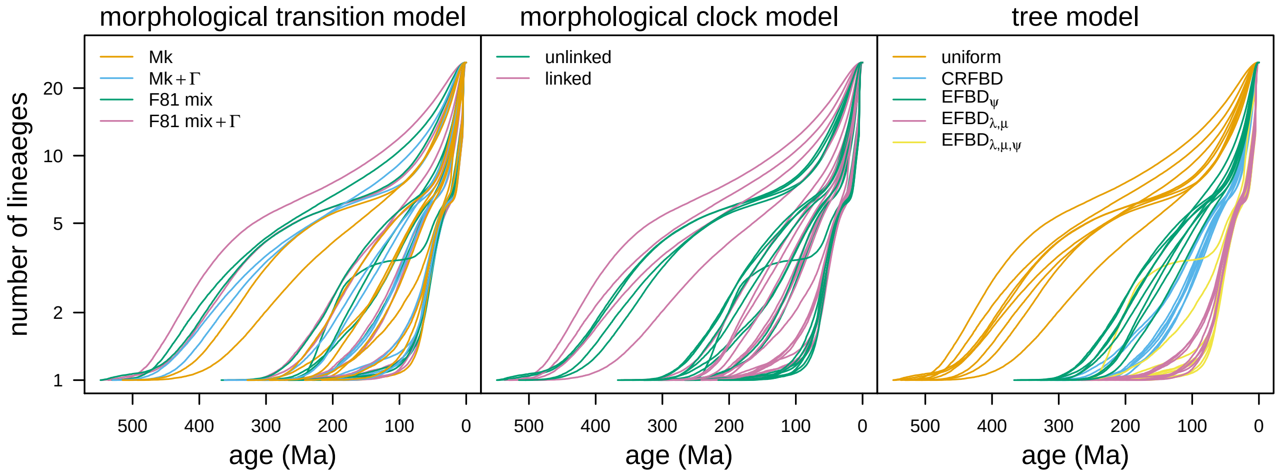


**Figure S19: Comparing lineage-through-time curves of extant taxa among model combinations.** Each curve corresponds to the lineage-through-time (LTT) curve of the extant Marattiaceae for a given model, averaged over the posterior distribution of trees from that model. The curves in each panel are color coded by morphological-transition model (left), morphological-clock model (middle), and tree model (right), respectively. We removed outgroup taxa to emphasize the influence of model specification on age estimates within our ingroup.

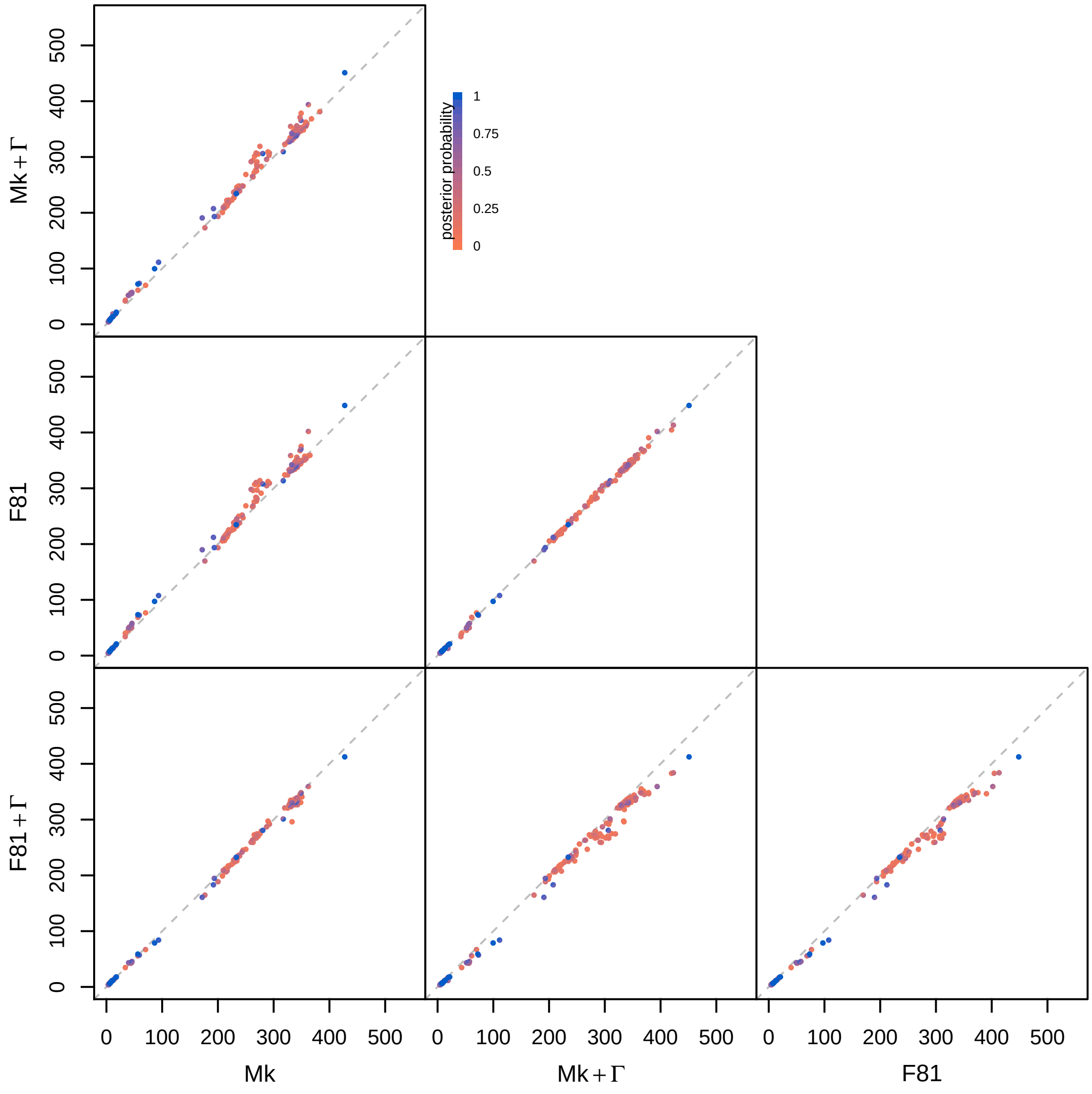


**Figure S20: Comparing divergence-time estimates for each clade by morphological transition model.** For each pair of models, we compare the posterior-mean age of each clade with posterior probability > 0.05 under the transition model on the x-axis against the transition model on the y-axis. Each point is divided in half, with the top-left semicircle colored by the posterior probability under the model on the y-axis and the bottom-right semicircle colored by the posterior probability under the model on the x-axis. In these comparisons, we assume the preferred morphological clock and tree models (linked and $\text{EFBD}_{\lambda,\mu}$, respectively).

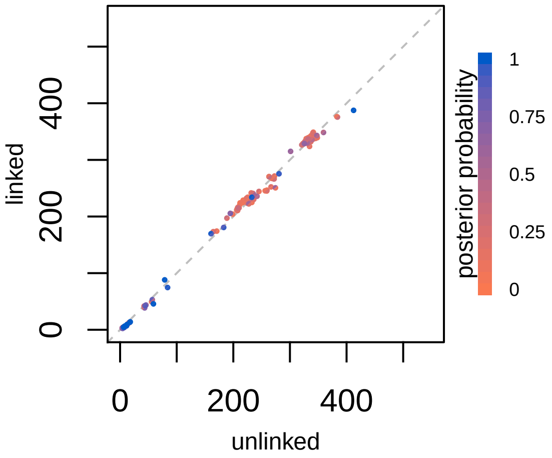


**Figure S21: Comparing divergence-time estimates for each clade by morphological clock model.** For each pair of models, we compare the posterior-mean age of each clade with posterior probability $> 0.05$ under the clock model on the x-axis against the clock model on the y-axis. Each point is divided in half, with the top-left semicircle colored by the posterior probability under the model on the y-axis and the bottom-right semicircle colored by the posterior probability under the model on the x-axis. In these comparisons, we assume the preferred morphological transition and tree models (F81 mixture $+\Gamma$ and $\text{EFBD}_{\lambda,\mu}$, respectively).

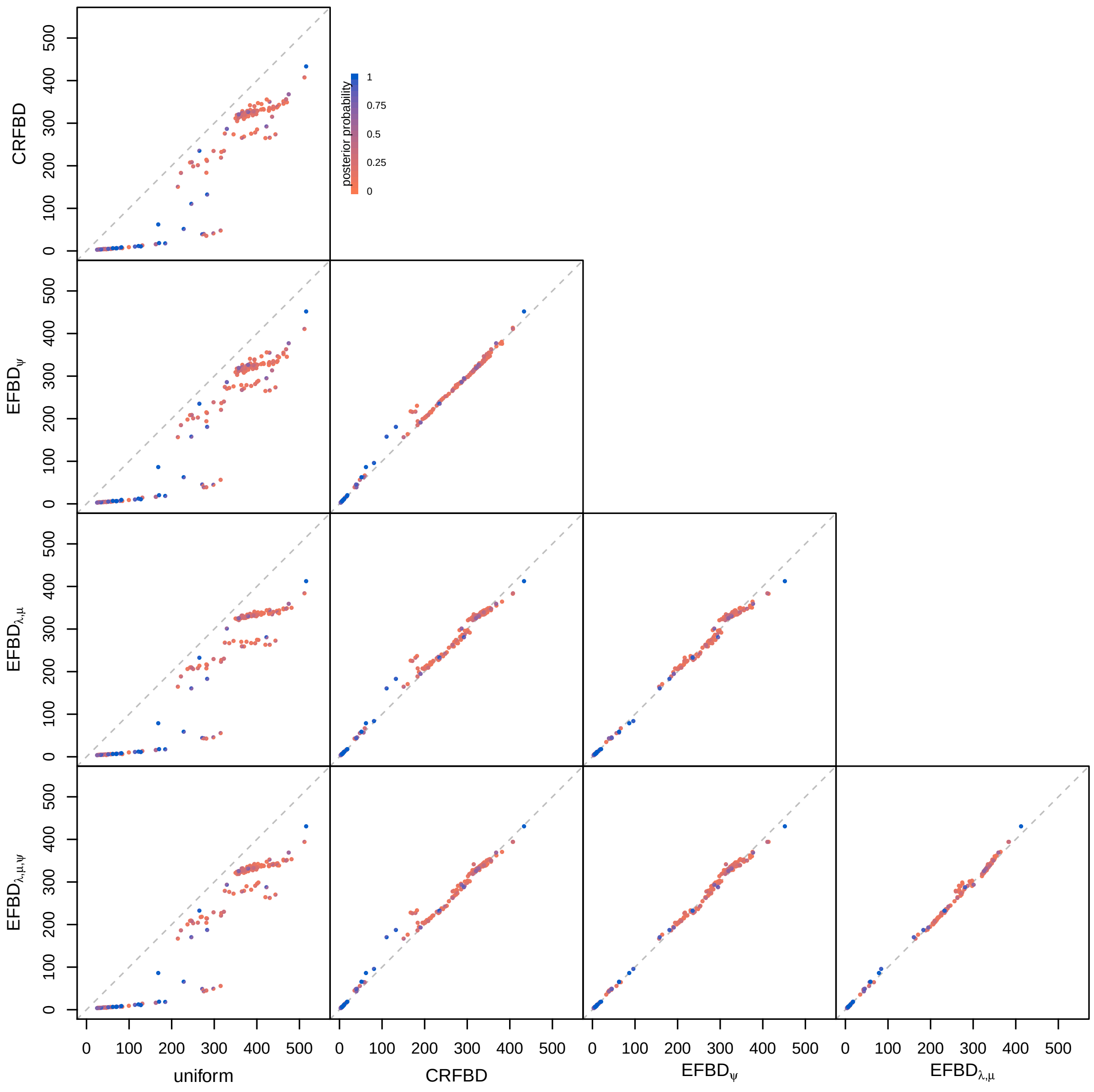


**Figure S22: Comparing divergence-time estimates for each clade by tree model.** For each pair of models, we compare the posterior-mean age of each clade with posterior probability $> 0.05$ under the tree model on the x-axis against the tree model on the y-axis. Each point is divided in half, with the top-left semicircle colored by the posterior probability under the model on the y-axis and the bottom-right semicircle colored by the posterior probability under the model on the x-axis. In these comparisons, we assume the preferred morphological transition and clock models (F81 mixture+Γ and linked, respectively).

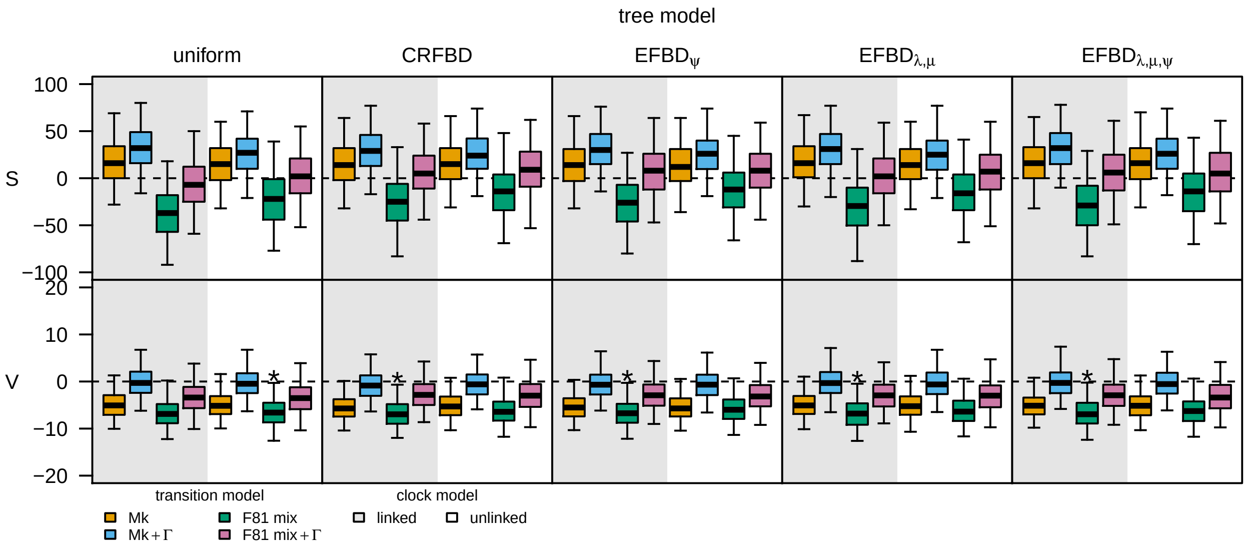


**Figure S23: Comparing model adequacy among model combinations.** Each boxplot represents the posterior-predictive distribution for a given model for a given statistic. The top row of panels correspond to the posterior-predictive distributions of the total parsimony score statistic, $S$; the bottom row of panels are for the variance in parsimony score statistic, $V$. Each column of panels corresponds to analyses under a given tree model. Within each panel, boxplots are colored by the morphological transition model, and regions of the panel are shaded according to the morphological clock model. Models that are inadequate at the $\alpha = 0.05$ level are indicated with an asterisk. These results assume the ingroup sampling fraction for all of the fossilized birth-death models; see Figure S40 for the results with the overall sampling fraction.

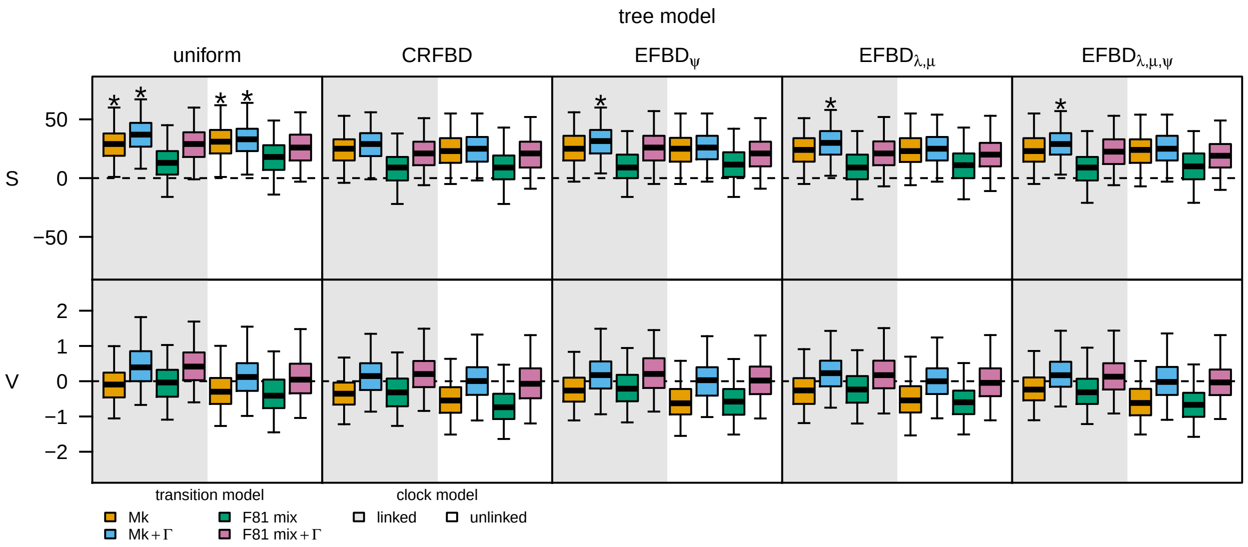


**Figure S24: Comparing model adequacy for extant taxa among model combinations.** Each boxplot represents the posterior-predictive distribution for a given model for a given statistic, computed after removing extinct taxa from the simulated and observed datasets. The top row of panels correspond to the posterior-predictive distributions of the total parsimony score statistic, $S$; the bottom row of panels are for the variance in parsimony score statistic, $V$. Each column of panels corresponds to analyses under a given tree model. Within each panel, boxplots are colored by the morphological transition model, and regions of the panel are shaded according to the morphological clock model. Models that are inadequate at the $\alpha = 0.05$ level are indicated with an asterisk.

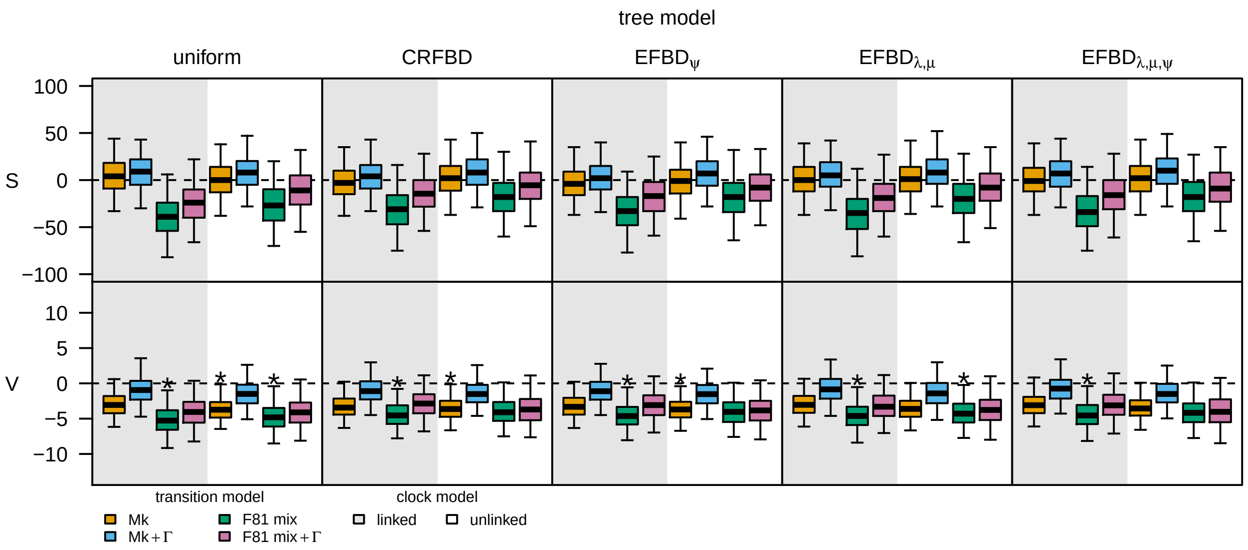


**Figure S25: Comparing model adequacy for extinct taxa among model combinations.** Each boxplot represents the posterior-predictive distribution for a given model for a given statistic, computed after removing extant taxa from the simulated and observed datasets. The top row of panels correspond to the posterior-predictive distributions of the total parsimony score statistic, $S$; the bottom row of panels are for the variance in parsimony score statistic, $V$. Each column of panels corresponds to analyses under a given tree model. Within each panel, boxplots are colored by the morphological transition model, and regions of the panel are shaded according to the morphological clock model. Models that are inadequate at the $\alpha = 0.05$ level are indicated with an asterisk.

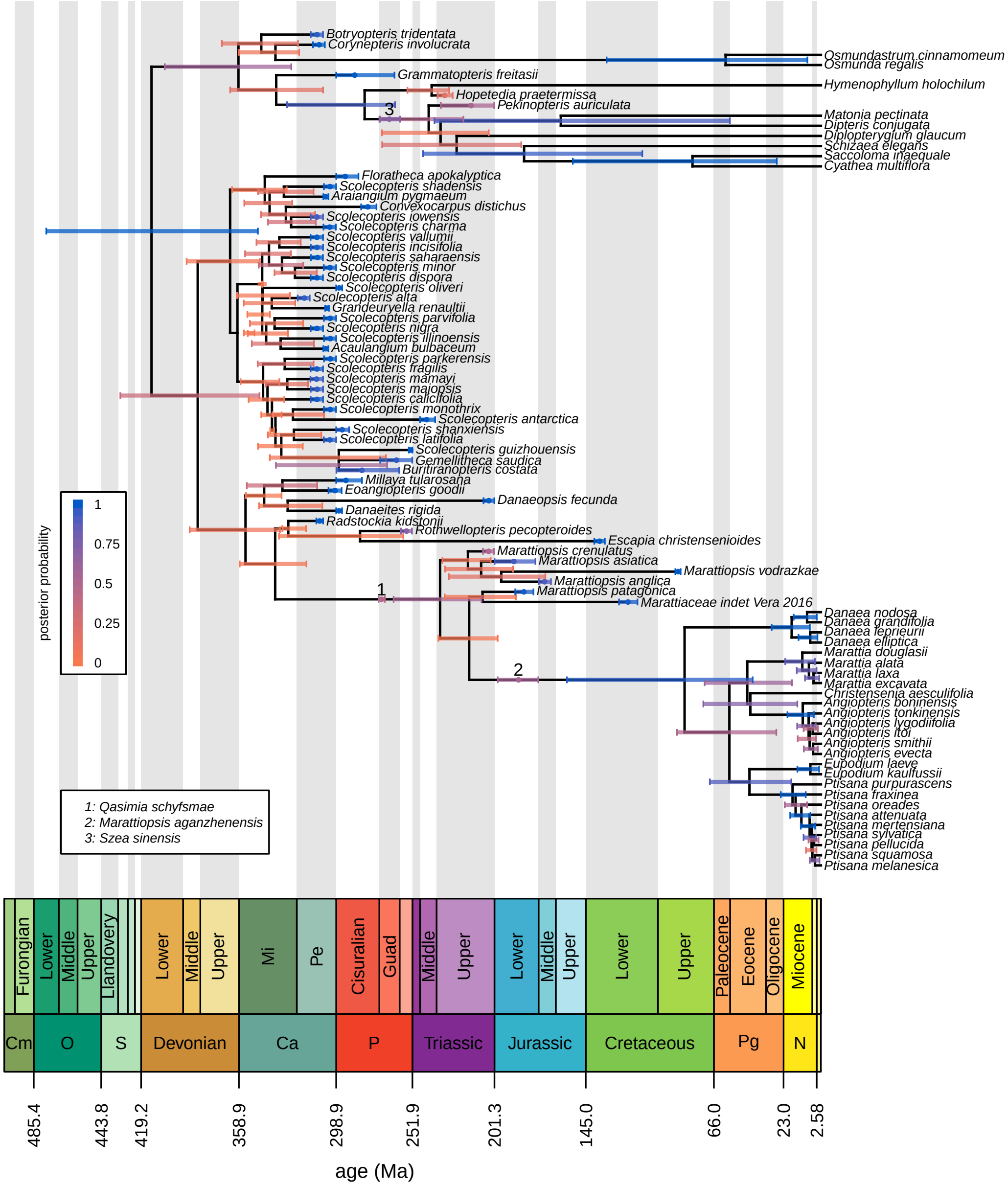


**Figure S26: The maximum clade credibility tree under the preferred model.** Bars correspond to the 95% credible interval of clade ages (for internal nodes) and tip ages (for fossils). Numbers on branches and associated age bars correspond to sampled ancestors (key bottom left). Bars are colored in proportion to the posterior probability of the clade for internal nodes, by the probability that the specimen is not a sampled ancestor for tips, and by the probability that the specimen is a sampled ancestor for sampled ancestors (legend, left).

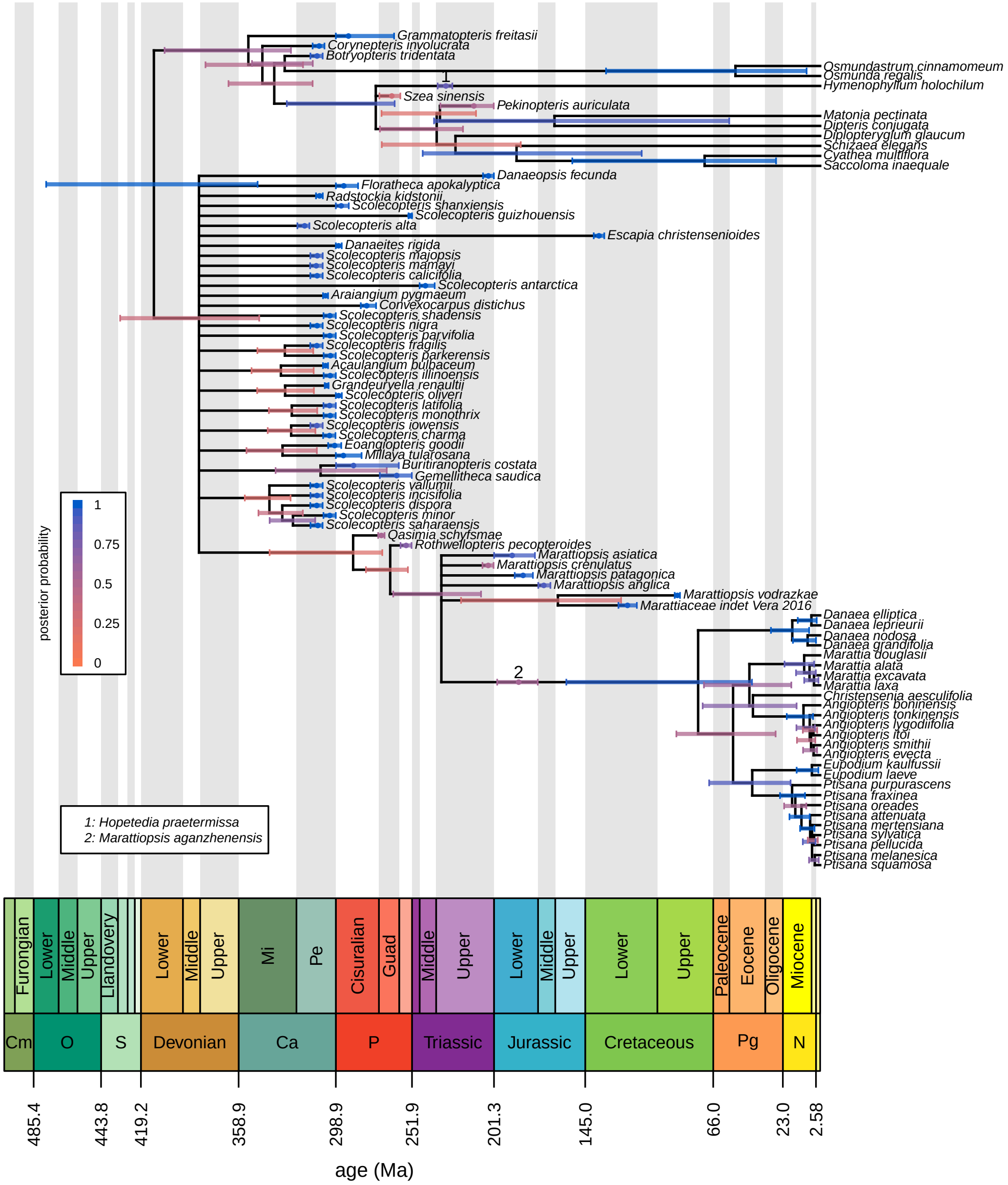


**Figure S27: The 20%-rule consensus tree under the preferred model.** Bars correspond to the 95% credible interval of clade ages (for internal nodes) and tip ages (for fossils). Numbers on braches and associated age bars correspond to sampled ancestors (key bottom left). Bars are colored in proportion to the posterior probability of the clade for internal nodes, by the probability that the specimen is not a sampled ancestor for tips, and by the probability that the specimen is a sampled ancestor for sampled ancestors (legend, left).

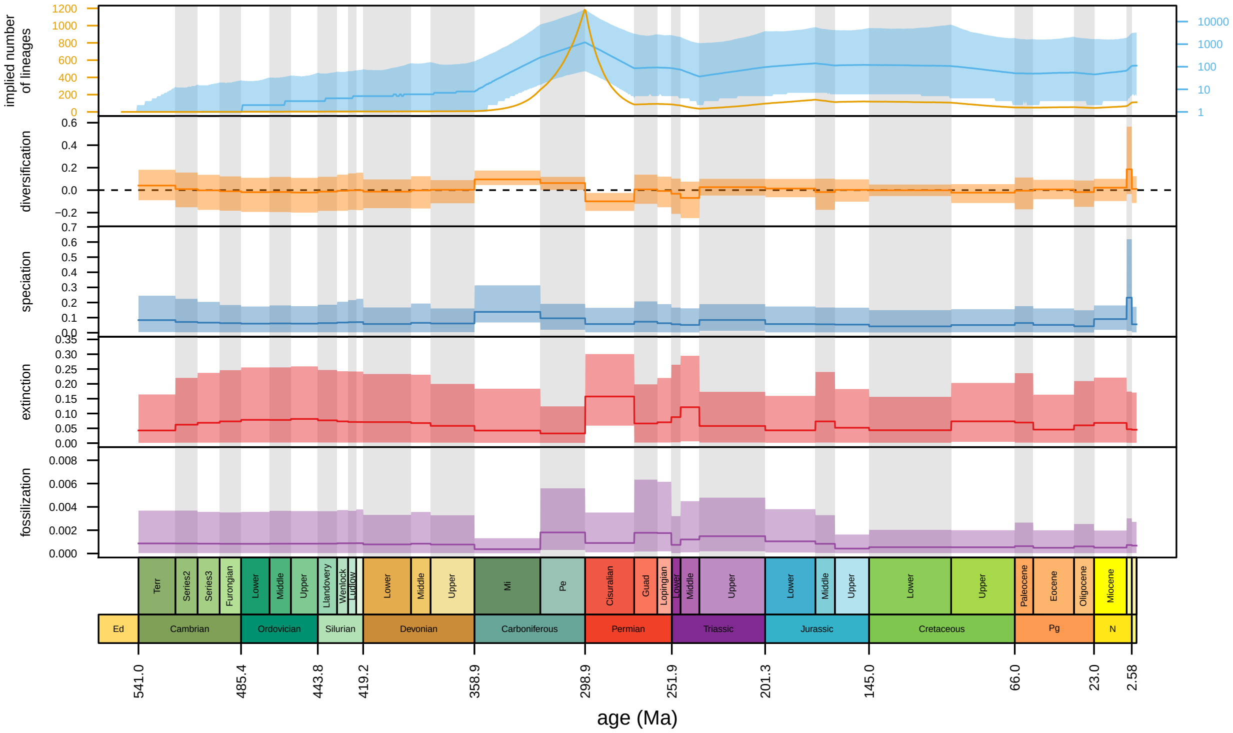


**Figure S28: Diversification through time under the EFBD$_{\lambda,\mu,\psi}$ model.** The top panel shows the implied total number of lineages over time (before pruning unsampled lineages). The next four panels correspond to the posterior distribution of epoch-specific net-diversification rates, speciation rates, extinction rates, and fossilization rates, respectively. In the top panel, the dark line represents the posterior median estimate, in the remaining panels it represents the posterior mean estimate. In all cases, and the shaded region corresponds to the 95% credible interval.

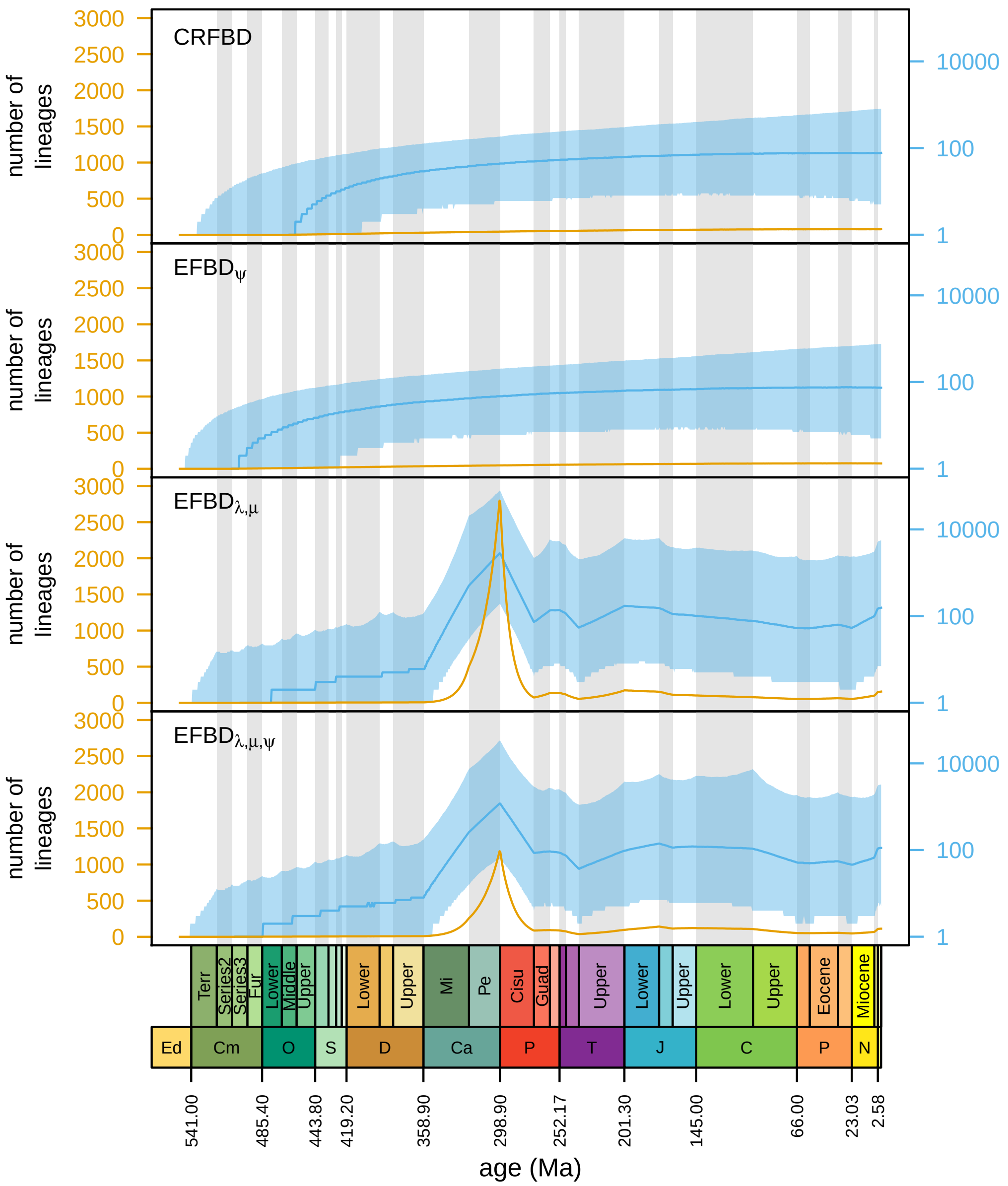


**Figure S29: Diversity-over-time under each fossilized birth-death model.** For each tree model, we draw samples from the posterior distribution and simulate lineages given those parameter values. We plot the median number of lineages at each time point on the absolute scale (orange, left axis) and the $\log_{10}$ scale (orange, right axis). Here, we assume the best morphological transition and clock models (F81 mixture+$\Gamma$ and linked, respectively).

#### S§6.2 Overall Sampling Fraction

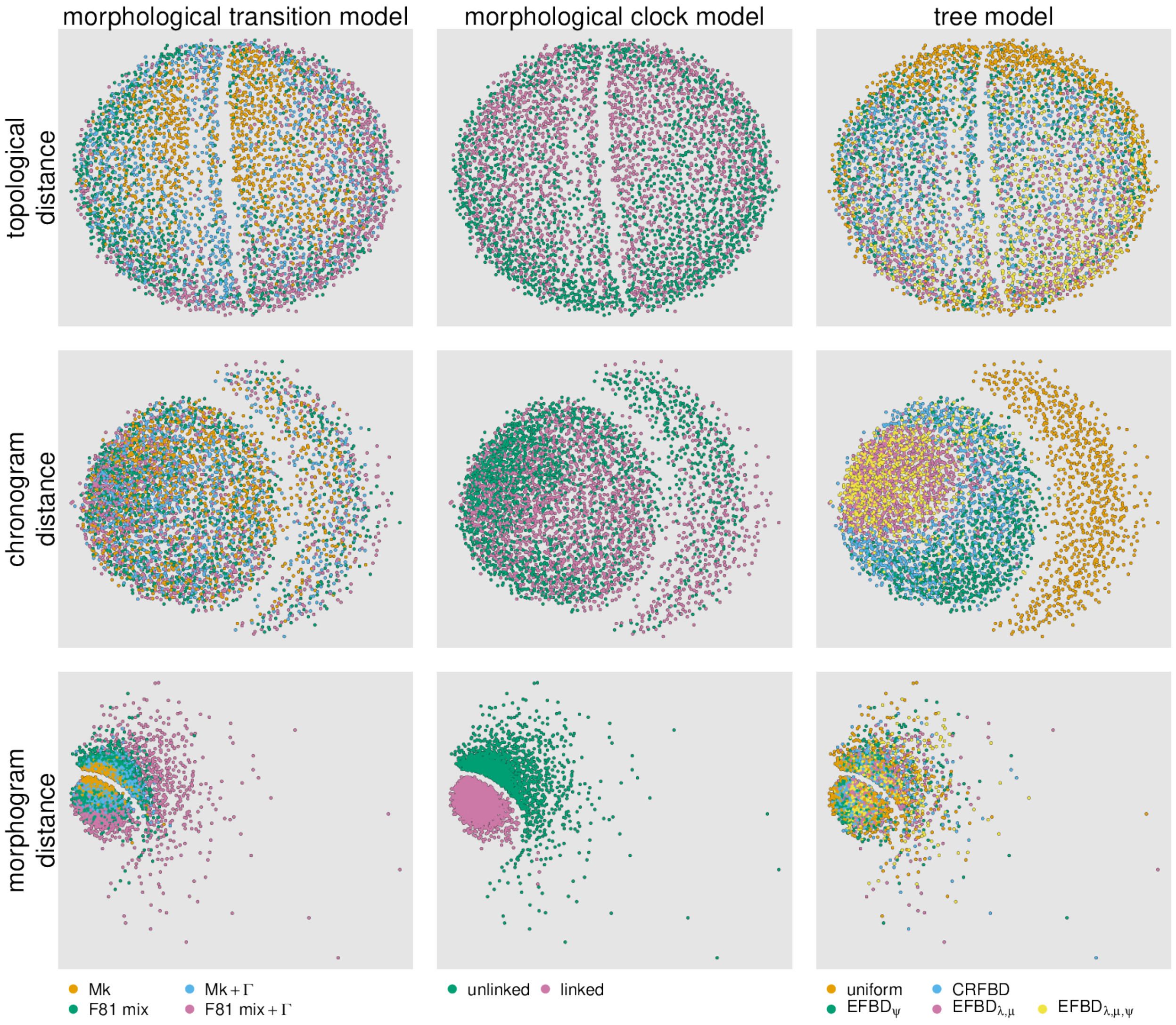


**Figure S30: Comparing distributions of trees among model combinations using the overall sampling fraction.** We compute the Robison-Foulds distance (RF, a measure of topological distance, top row), the Kühner-Felsenstein distance (KF, a distance metric that incorporates both topology and branch lengths between morphological phylograms ("morphograms", middle row) and chromograms (bottom row). We then plot the (square-root transformed) distances in two-dimensional space using multi-dimensional scaling (MDS); each point represents the location of a given sampled tree in tree space according to the distance metric. We color points according to the tree model (column 1), the morphological matrix model (column 2), or the morphological clock model (column 3). These results assume the ingroup sampling fraction for all of the fossilized birth-death models.

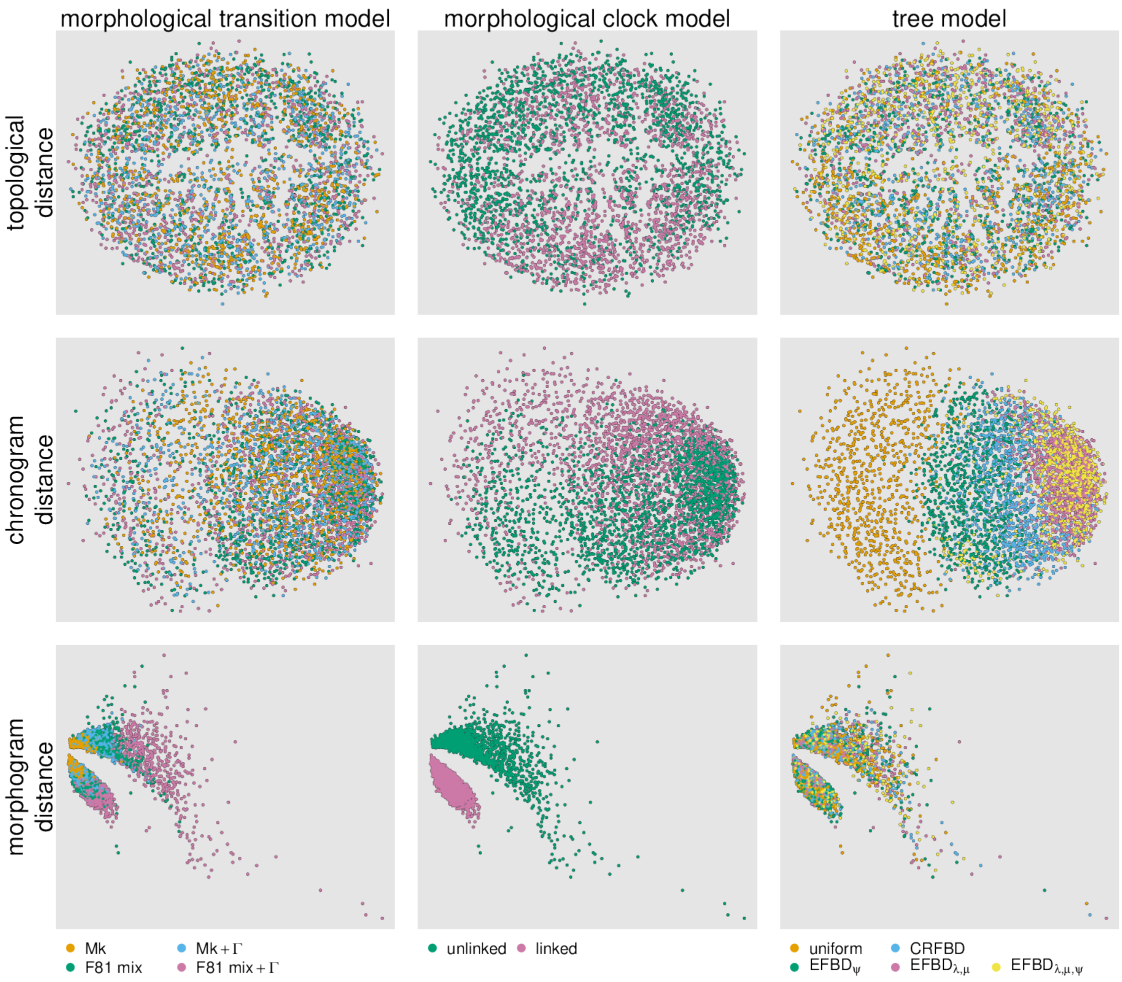


**Figure S31: Comparing distributions of molecular trees among model combinations using the overall sampling fraction.** We compute the Robinson-Foulds distance (RF, a measure of topological distance, top row), the Kühner-Felsenstein distance (KF, a distance metric that incorporates both topology and branch lengths) between morphological phylograms ("morphograms", middle row) and chronograms (bottom row). We removed taxa without molecular data before computing distances. We then plot the (square-root transformed) distances in two-dimensional space using multi-dimensional scaling (MDS); each point represents the location of a given sampled tree in tree space according to the distance metric. We color points according to the morpological transition model (column 1), the morphological clock model (column 2), or the tree model (column 3).

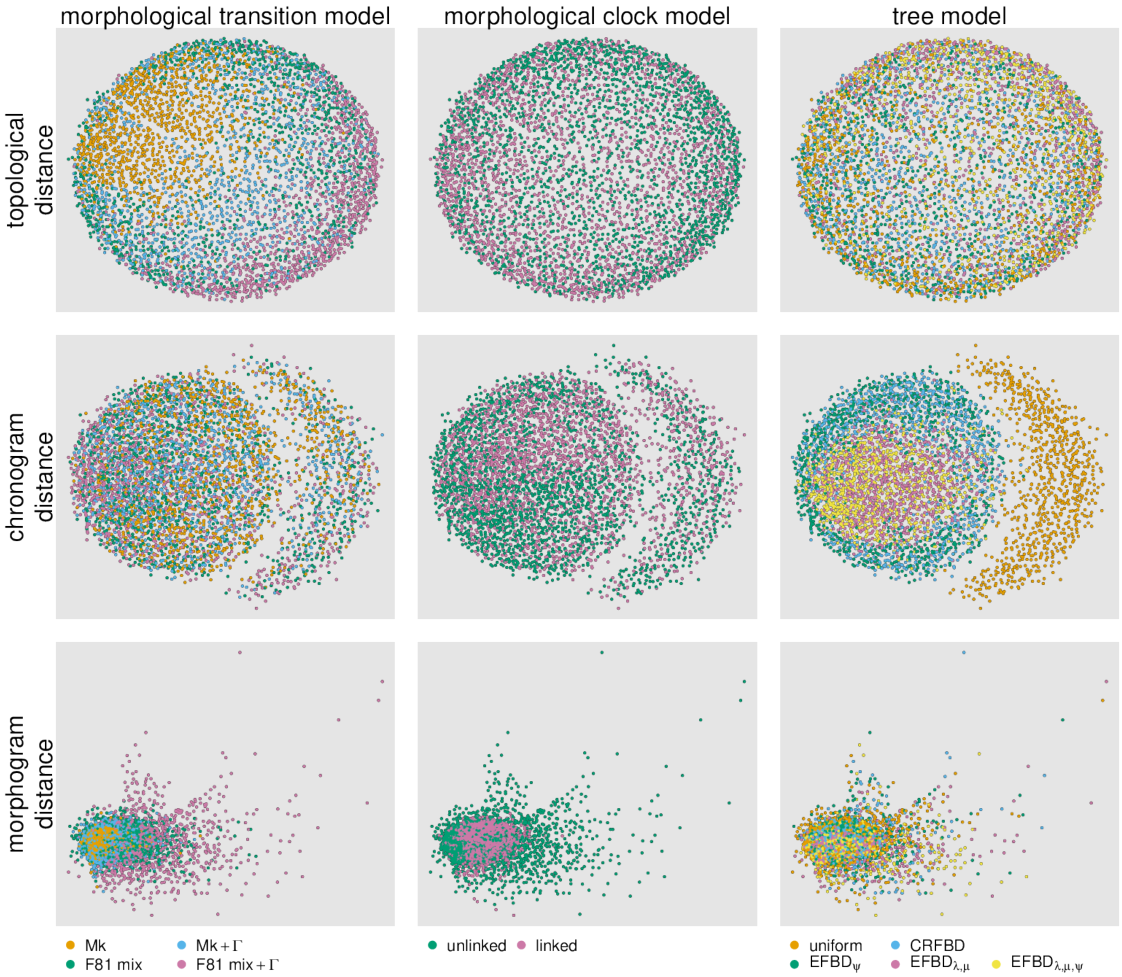


**Figure S32: Comparing distributions of extinct trees among model combinations using the overall sampling fraction.** We compute the Robinson-Foulds distance (RF, a measure of topological distance, top row), the Kühner-Felsenstein distance (KF, a distance metric that incorporates both topology and branch lengths) between morphological phylograms ("morphograms", middle row) and chronograms (bottom row). We removed extant taxa before computing distances. We then plot the (square-root transformed) distances in two-dimensional space using multi-dimensional scaling (MDS); each point represents the location of a given sampled tree in tree space according to the distance metric. We color points according to the morpological transition model (column 1), the morphological clock model (column 2), or the tree model (column 3).

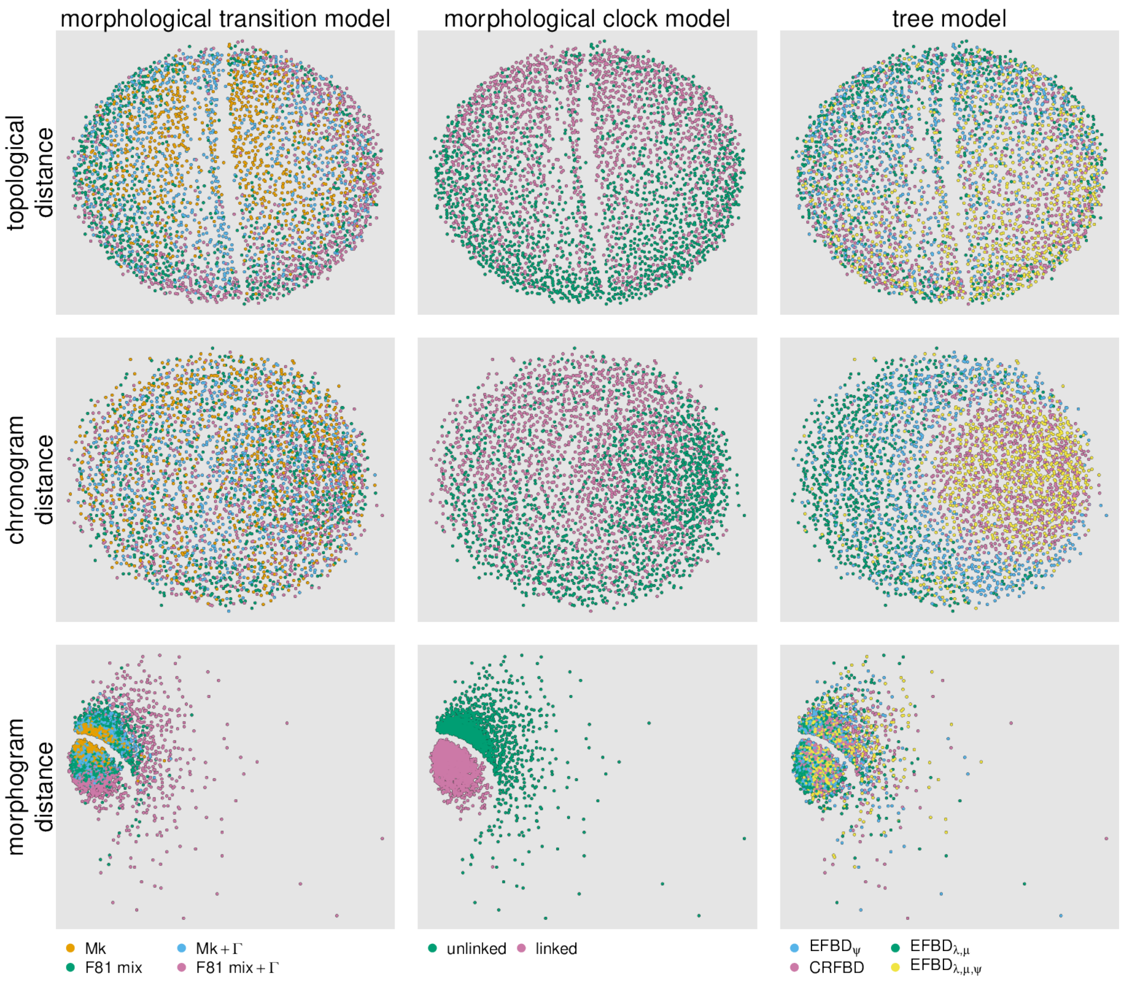


**Figure S33: Comparing distributions of trees among model combinations using the overall sampling fraction, excluding the uniform tree model.** We compute the Robinson-Foulds distance (RF, a measure of topological distance, top row), the Kühner-Felsenstein distance (KF, a distance metric that incorporates both topology and branch lengths) between morphological phylograms ("morphograms", middle row) and chronograms (bottom row). We removed extant taxa before computing distances. We then plot the (square-root transformed) distances in two-dimensional space using multi-dimensional scaling (MDS); each point represents the location of a given sampled tree in tree space according to the distance metric. We color points according to the morpological transition model (column 1), the morphological clock model (column 2), or the tree model (column 3).

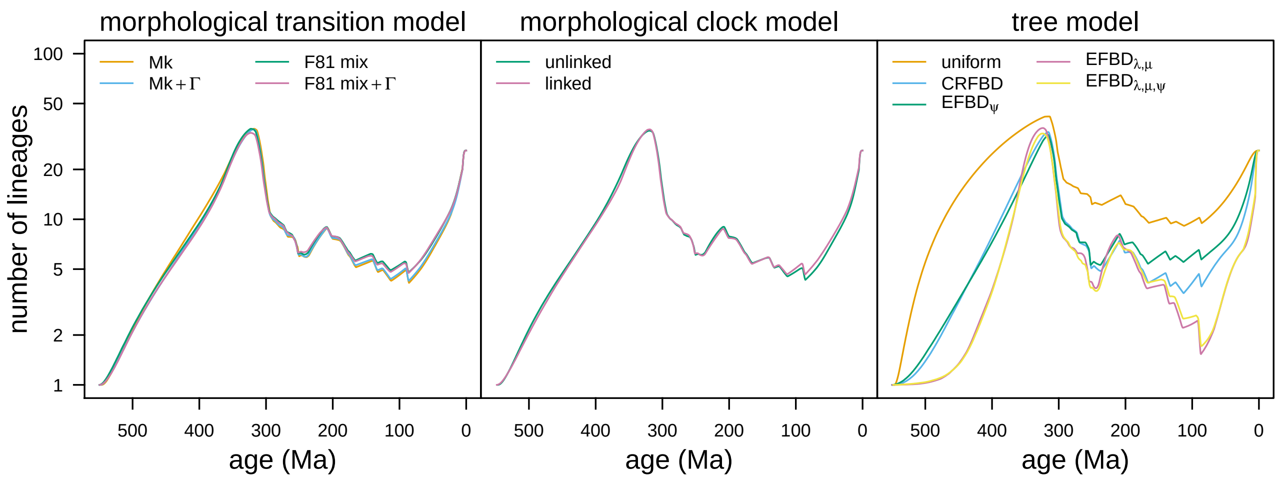


**Figure S34: Average lineage-through-time curves among model combinations using the overall sampling fraction.** For each focal model component (morphological transition model, and morphological clock model, and the the tree model, respectively), we compute the LTT curve averaged over the remaining model components, *i.e.*, the average number of branches in the phylogeny at a given time, averaged over all model combinations that share the focal model component. We removed outgroup taxa to emphasize the influence of model specification on age estimates within our ingroup. Left) We compute the average LTT for each of the 40 models (from 2000 sampled trees for each model), then compute the mean of the resulting average LTT among models with the same morphological-transition model. Middle) As in left, but we compute the mean of the average LTT among models with the same morphological clock model. Right) As in left, but we compute the mean of the average LTT among models with the same tree model.

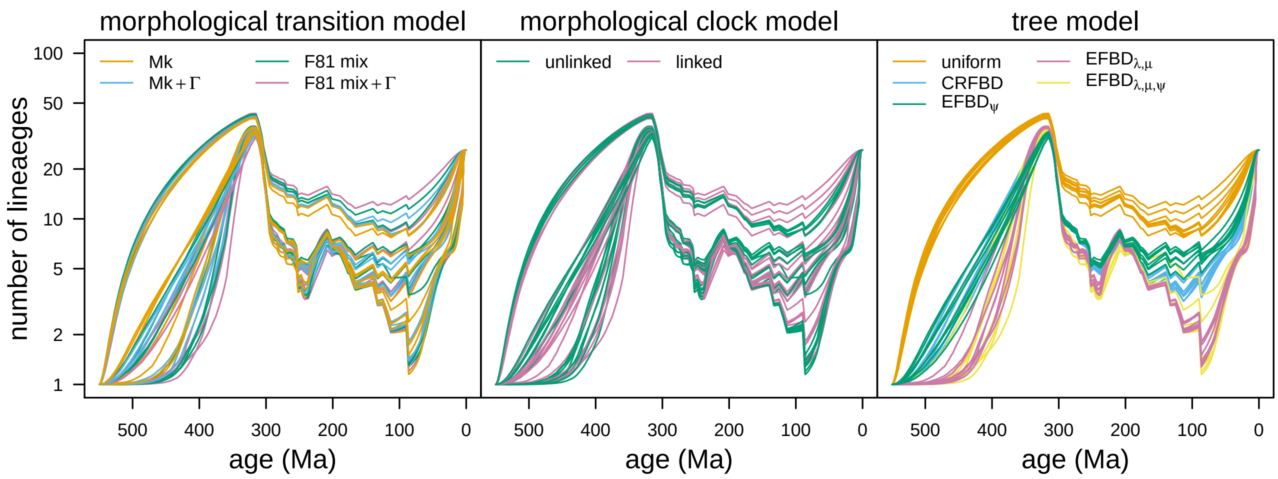


**Figure S35: Comparing lineage-through-time curves among model combinations using the overall sampling fraction.** Each curve corresponds to the lineage-through-time (LTT) curve of the extant Marattiacaea for a given model, averaged over the posterior distribution of trees from that model. The curves in each panel are color coded by morphological-transition model (left), morphological-clock model (middle), and tree model (right), respectively. We removed outgroup taxa to emphasize the influence of model specification on age estimates within our ingroup.

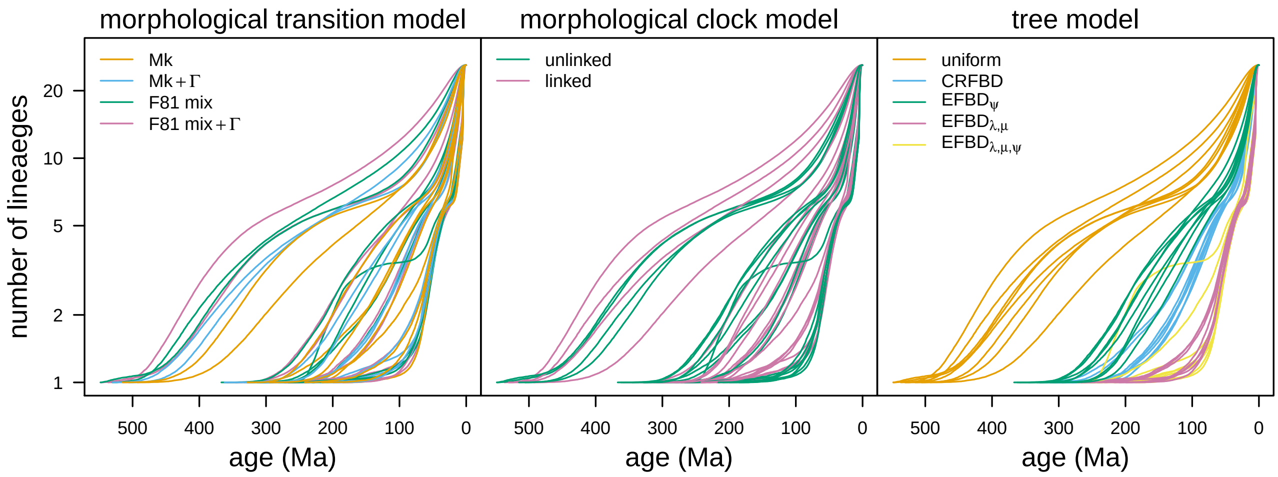


**Figure S36: Comparing lineage-through-time curves of extant taxa among model combinations using the overall sampling fraction.** Each curve corresponds to the lineage-through-time (LTT) curve of the extant Marattiacaea for a given model, averaged over the posterior distribution of trees from that model. The curves in each panel are color coded by morphological-transition model (left), morphological-clock model (middle), and tree model (right), respectively. We removed outgroup taxa to emphasize the influence of model specification on age estimates within our ingroup.

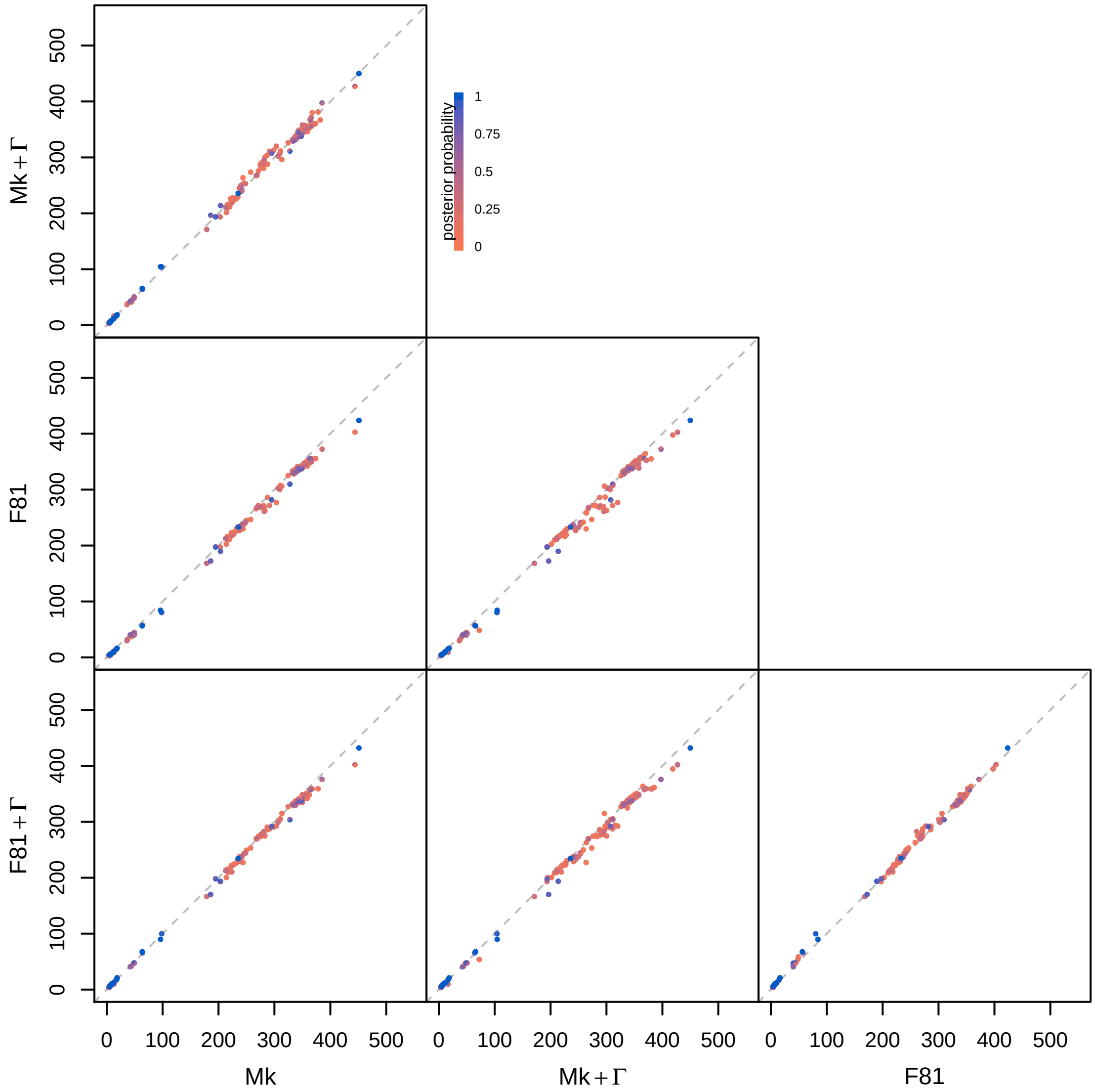


**Figure S37: Comparing divergence-time estimates for each clade by morphological transition model using the overall sampling fraction.** For each pair of models, we compare the posterior-mean age of each clade with posterior probability > 0.05 under the transition model on the x-axis against the transition model on the y-axis. Each point is divided in half, with the top-left semicircle colored by the posterior probability under the model on the y-axis and the bottom-right semicircle colored by the posterior probability under the model on the x-axis. In these comparisons, we assume the preferred morphological clock and tree models (linked and $\text{EFBD}_{\lambda,\mu}$, respectively).

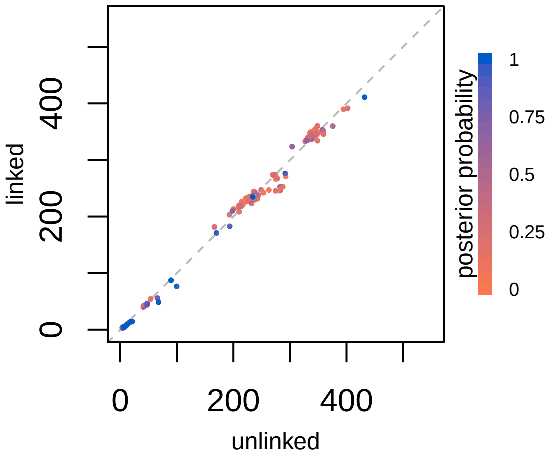


**Figure S38: Comparing divergence-time estimates for each clade by morphological clock model using the overall sampling fraction.** For each pair of models, we compare the posterior-mean age of each clade with posterior probability $> 0.05$ under the clock model on the x-axis against the clock model on the y-axis. Each point is divided in half, with the top-left semicircle colored by the posterior probability under the model on the y-axis and the bottom-right semicircle colored by the posterior probability under the model on the x-axis. In these comparisons, we assume the preferred morphological transition and tree models (F81 mixture+$\Gamma$ and EFBD$_{\lambda,\mu}$, respectively).

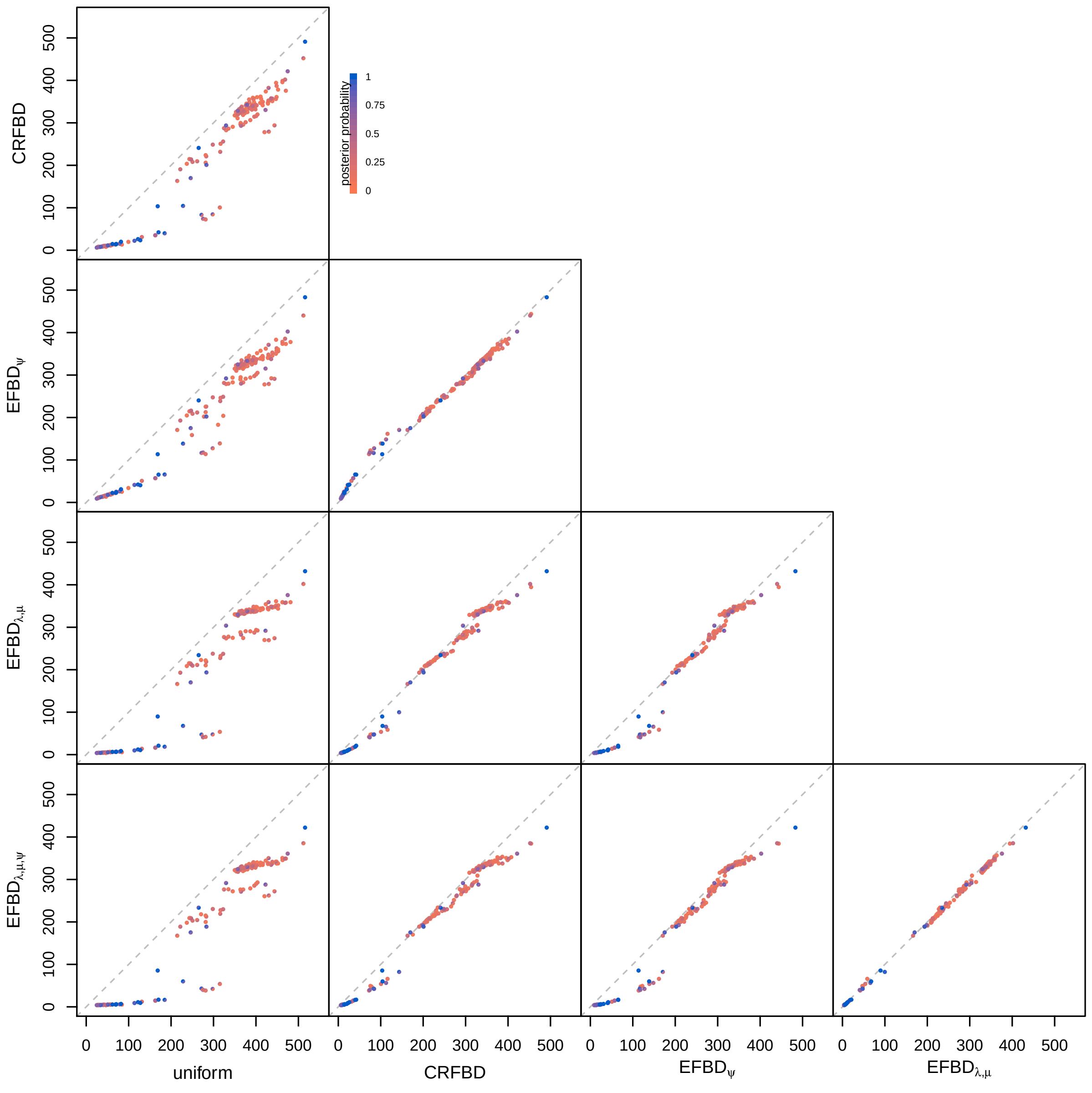


**Figure S39: Comparing divergence-time estimates for each clade by tree model using the overall sampling fraction.** For each pair of models, we compare the posterior-mean age of each clade with posterior probability $> 0.05$ under the tree model on the x-axis against the tree model on the y-axis. Each point is divided in half, with the top-left semicircle colored by the posterior probability under the model on the y-axis and the bottom-right semicircle colored by the posterior probability under the model on the x-axis. In these comparisons, we assume the preferred morphological transition and clock models (F81 mixture+$\Gamma$ and linked, respectively).

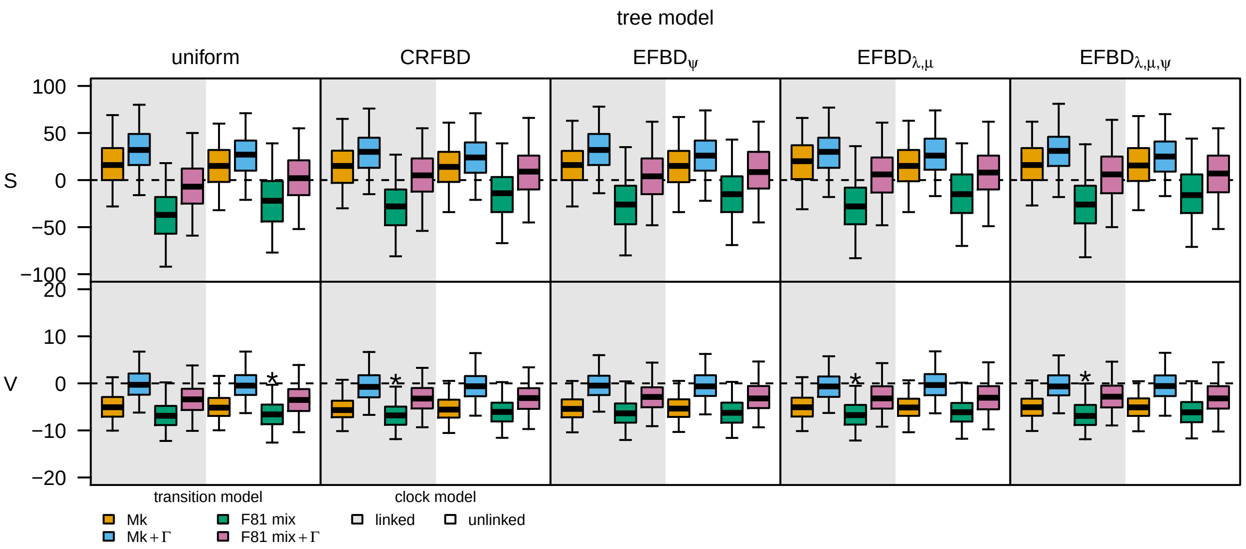


**Figure S40: Comparing model adequacy among model combinations using the overall sampling fraction.** Each boxplot represents the posterior-predictive distribution for a given model for a given statistic. The top row of panels correspond to the posterior-predictive distributions of the total parsimony score statistic, $S$; the bottom row of panels are for the variance in parsimony score statistic, $V$. Each column of panels corresponds to analyses under a given tree model. Within each panel, boxplots are colored by the morphological matrix model, and regions of the panel are colored according to the morphological clock model. Models that are inadequate at the $\alpha = 0.05$ level are indicated with an asterisk.

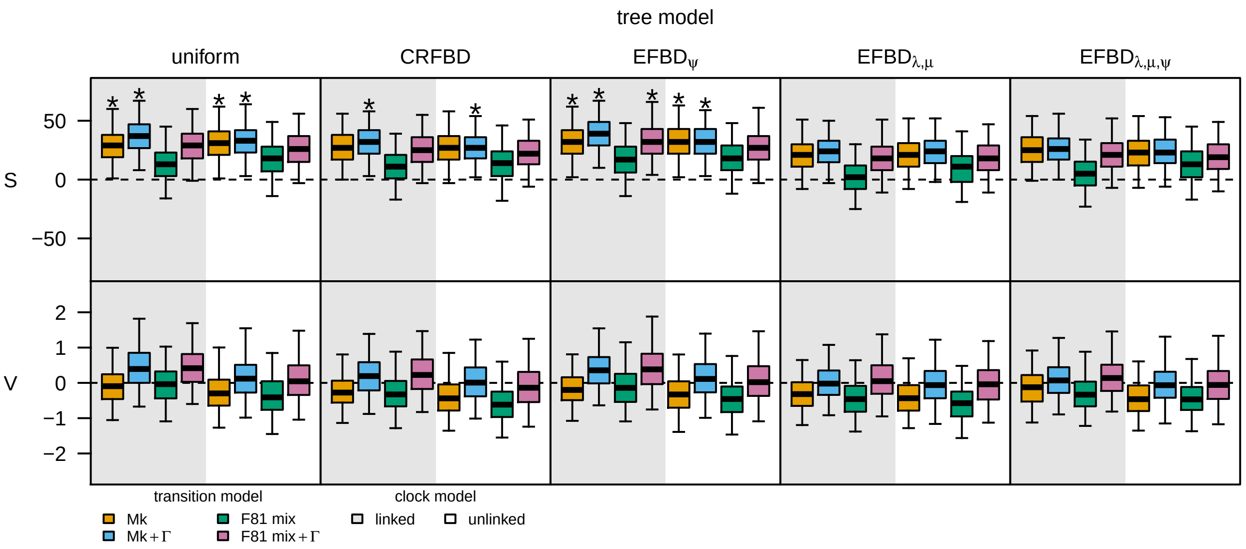


**Figure S41: Comparing model adequacy for extant taxa among model combinations using the overall sampling fraction.** Each boxplot represents the posterior-predictive distribution for a given model for a given statistic, computed after removing extinct taxa from the simulated and observed datasets. The top row of panels correspond to the posterior-predictive distributions of the total parsimony score statistic, $S$; the bottom row of panels are for the variance in parsimony score statistic, $V$. Each column of panels corresponds to analyses under a given tree model. Within each panel, boxplots are colored by the morphological transition model, and regions of the panel are shaded according to the morphological clock model. Models that are inadequate at the $\alpha = 0.05$ level are indicated with an asterisk.

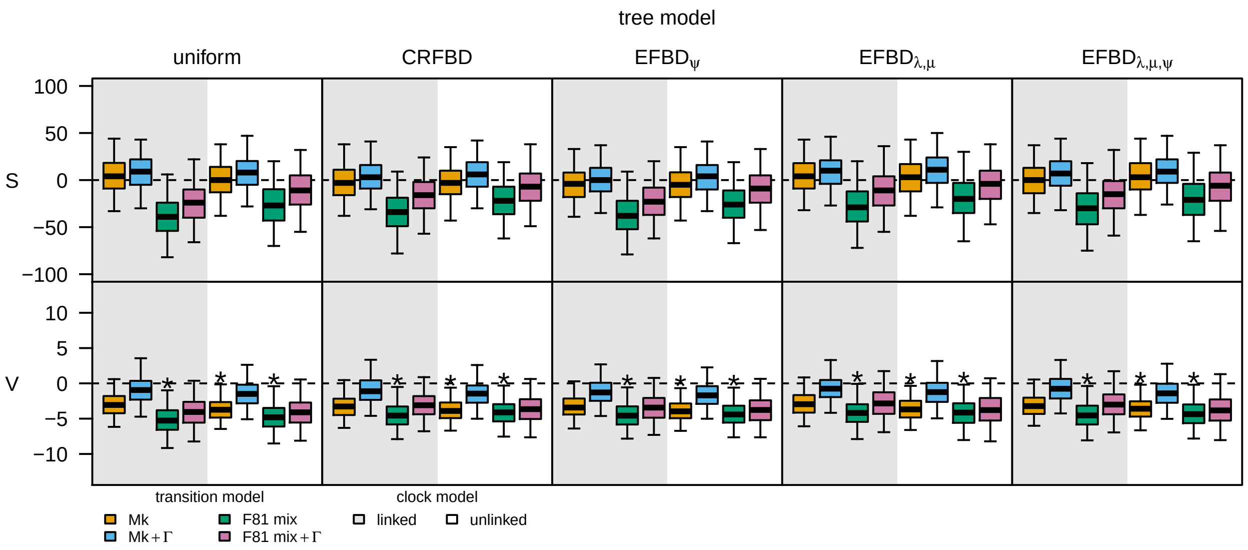


**Figure S42: Comparing model adequacy for extinct taxa among model combinations using the overall sampling fraction.** Each boxplot represents the posterior-predictive distribution for a given model for a given statistic, computed after removing extant taxa from the simulated and observed datasets. The top row of panels correspond to the posterior-predictive distributions of the total parsimony score statistic, $S$; the bottom row of panels are for the variance in parsimony score statistic, $V$. Each column of panels corresponds to analyses under a given tree model. Within each panel, boxplots are colored by the morphological transition model, and regions of the panel are shaded according to the morphological clock model. Models that are inadequate at the $\alpha = 0.05$ level are indicated with an asterisk.

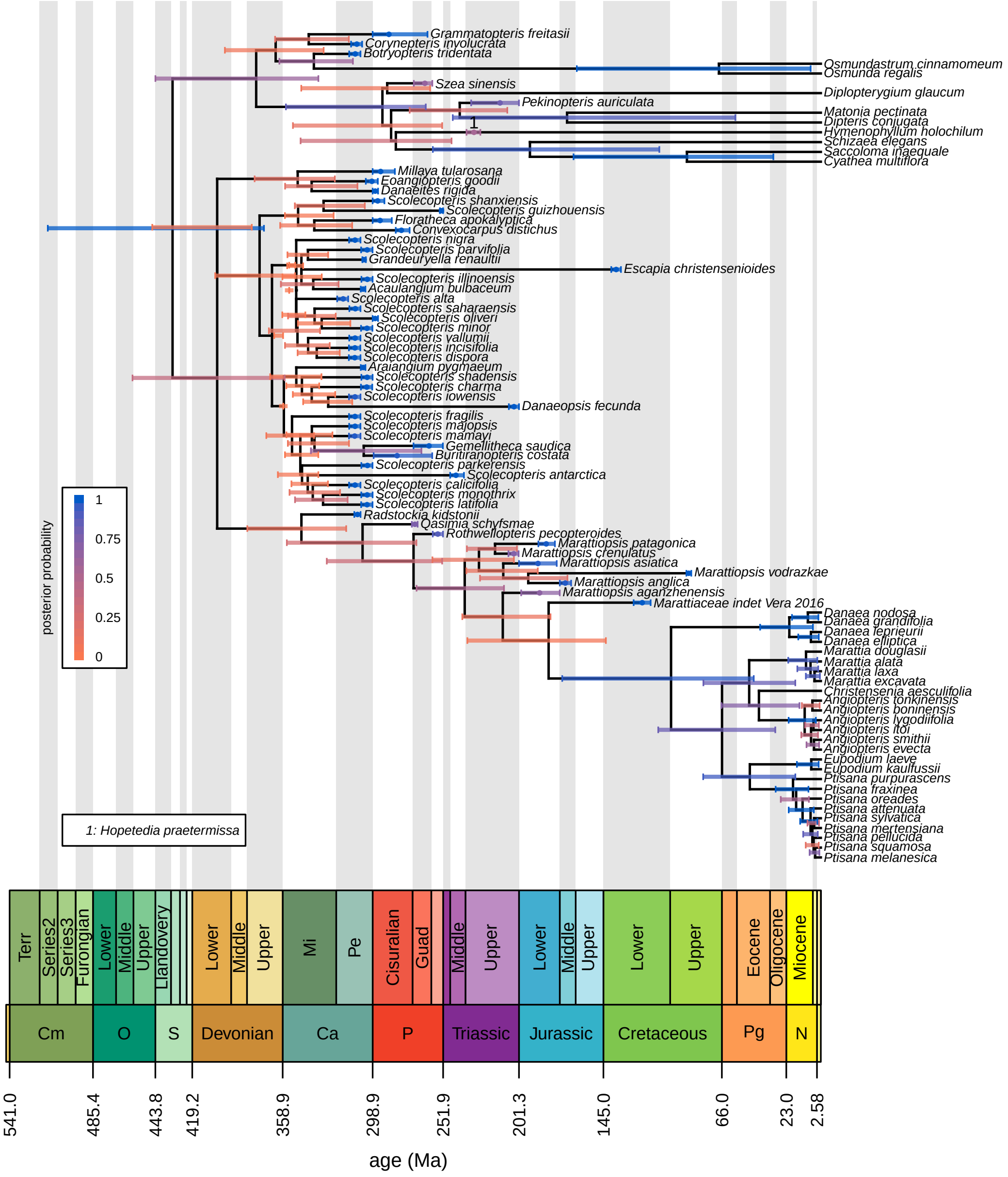


**Figure S43: The maximum clade credibility tree under the preferred model with overall sampling fraction.** Bars correspond to the 95% credible interval of clade ages (for internal nodes) and tip ages (for fossils). Numbers on braches and associated age bars correspond to sampled ancestors (key bottom left). Bars are colored in proportion to the posterior probability of the clade for internal nodes, by the probability that the specimen is not a sampled ancestor for tips, and by the probability that the specimen is a sampled ancestor for sampled ancestors (legend, left).

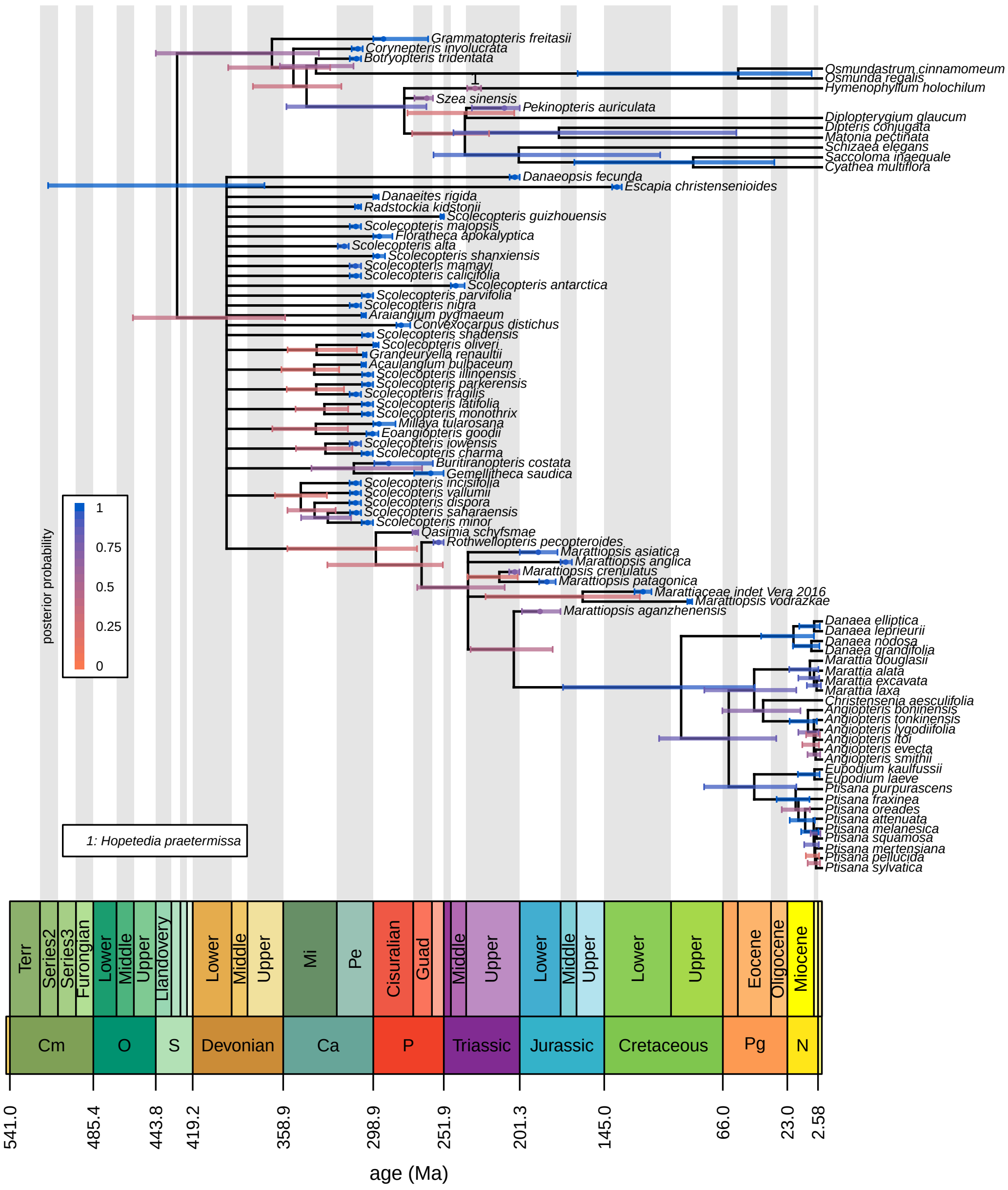


**Figure S44: The 20%-rule consensus tree under the preferred model with overall sampling fraction.** Bars correspond to the 95% credible interval of clade ages (for internal nodes) and tip ages (for fossils). Numbers on braches and associated age bars correspond to sampled ancestors (key bottom left). Bars are colored in proportion to the posterior probability of the clade for internal nodes, by the probability that the specimen is not a sampled ancestor for tips, and by the probability that the specimen is a sampled ancestor for sampled ancestors (legend, left).

#### S§6.3 Ingroup vs. Overall Sampling Fraction

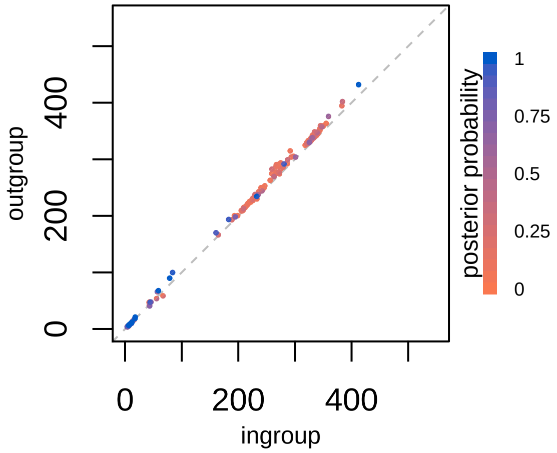


**Figure S45: Comparing divergence-time estimates for each clade by assumed sampling fraction.** For each pair of models, we compare the posterior-mean age of each clade with posterior probability $> 0.05$ under the sampling fraction on the x-axis against the sampling fraction on the y-axis. Each point is divided in half, with the top-left semicircle colored by the posterior probability under the model on the y-axis and the bottom-right semicircle colored by the posterior probability under the model on the x-axis. In these comparisons, we assume the preferred morphological transition and clock models and tree model (F81 mixture+$\Gamma$, linked, EFBD$_{\lambda,\mu}$ respectively).

#### S§6.4 Uniform Tree Model

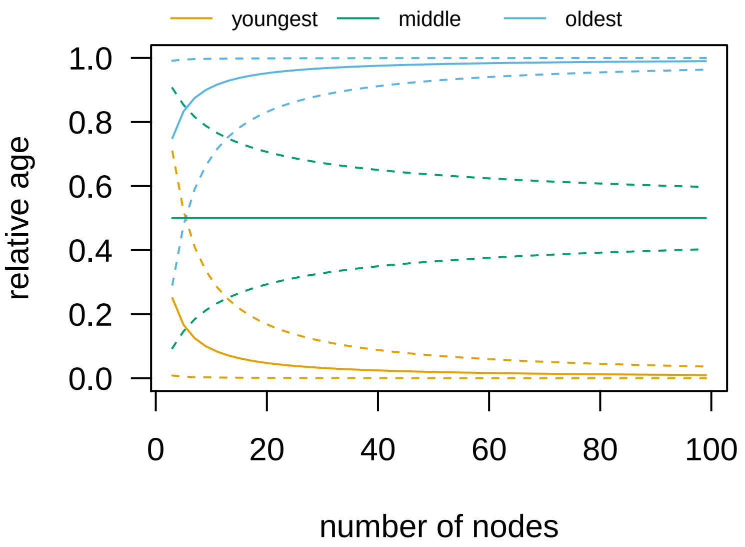


**Figure S46: The uniform tree model is an informative prior on node ages.** Here we consider a tree with all extant species for the sake of simplicity, but the logic applies equally to trees with extinct tips. Under the uniform tree model, node ages are drawn from an ordered uniform distribution. Because the ages are ordered, the age of the $i^{\text{th}}$ node in a tree with $n$ internal nodes is the $i^{\text{th}}$ order statistic a uniform distribution. Assuming the age of the tree is 1 arbitrary time unit, the $i^{\text{th}}$ node age is a Beta random variable with $\alpha = i$ and $\beta = n + 1 - i$. We show the resulting prior mean (solid lines) and 95% marginal prior interval (dashed lines) for the youngest node, the oldest node, and the middle node as a function of the total number of internal nodes. As the number of nodes increases, the first and last nodes are pushed toward the boundaries and have highly constrained (informative) priors. Likewise, the middle node is halfway between the present and the root (/stem) age with increasing prior probability as the number of nodes increases.

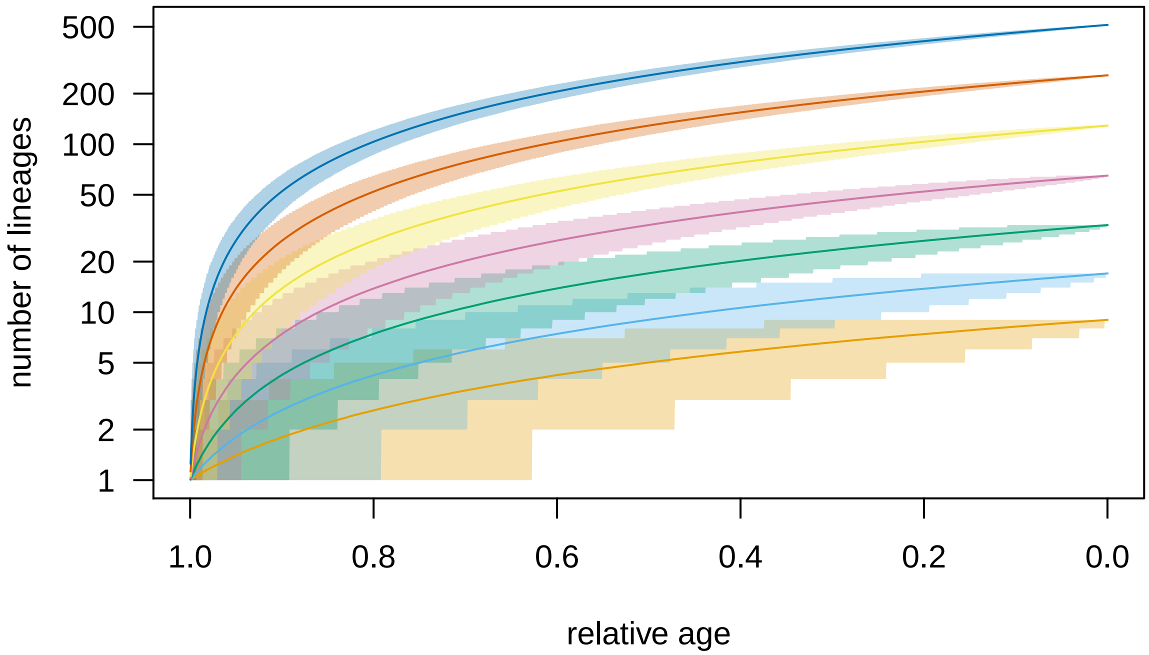


**Figure S47: The uniform model is an informative prior on lineage-through-time curves.** We simulated trees of extant species under the uniform tree model, and computed the distribution of resulting LTT curves as a function of the number of extant species (lines are prior mean number of lineages, shaded regions are the 95% prior interval). As the number of species grows, the distribution of ages becomes increasingly narrow.

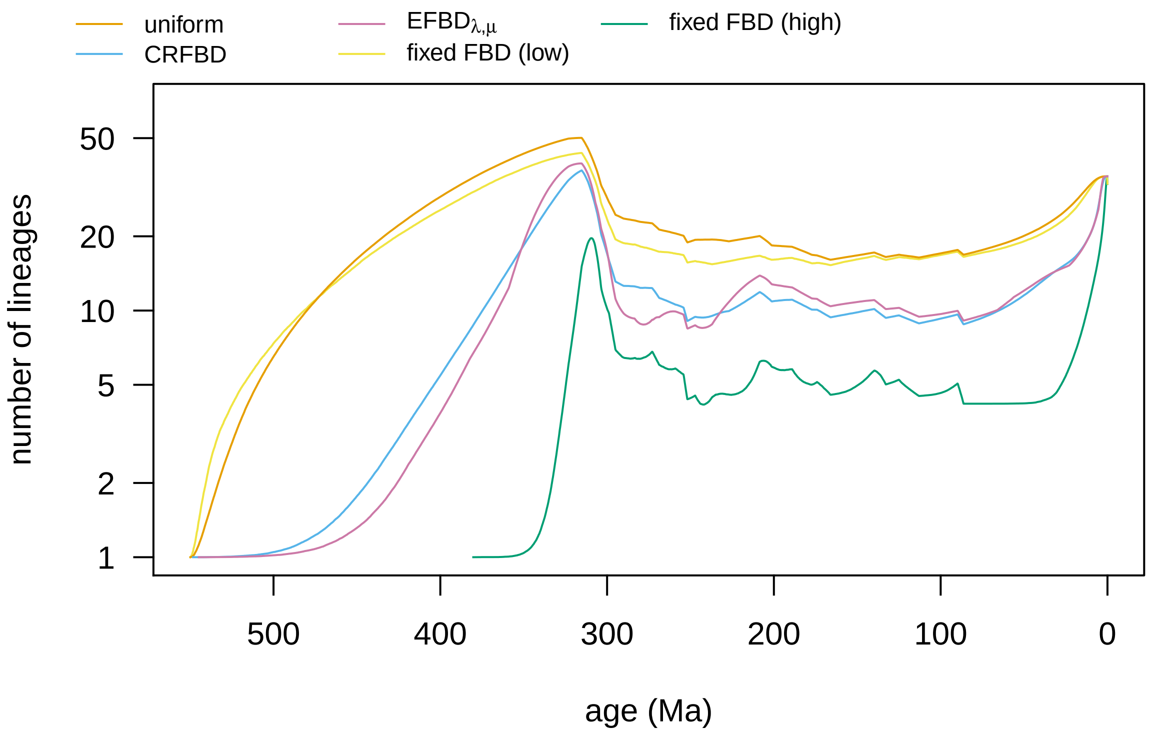


**Figure S48: The fossilized birth-death model is an informative prior on node ages when hyperparameters are fixed.** We estimated the posterior distribution of trees under two constant-rate fossilized birth-death models with fixed hyperparameters: the "low" fixed FBD ($\lambda = 0.001, \mu = 0.0009, \psi = 0.0001$, yellow LTT curve) and the "high" fixed FBD ($\lambda = 0.5, \mu = 0.45, \psi = 0.01$, green LTT curve). These parameter values were selected to be arbitrarily high or low to demonstrate the potential informativeness of the FBD with fixed hyperparameters. LTT curves under both models are significantly different from each other and every other model, whereas the LTT curves under both FBD models with estimated hyperpameters (blue and pink curves) are relatively similar, indicating that with fixed hyperparameters the fossilized birth-death model strongly informs node ages.

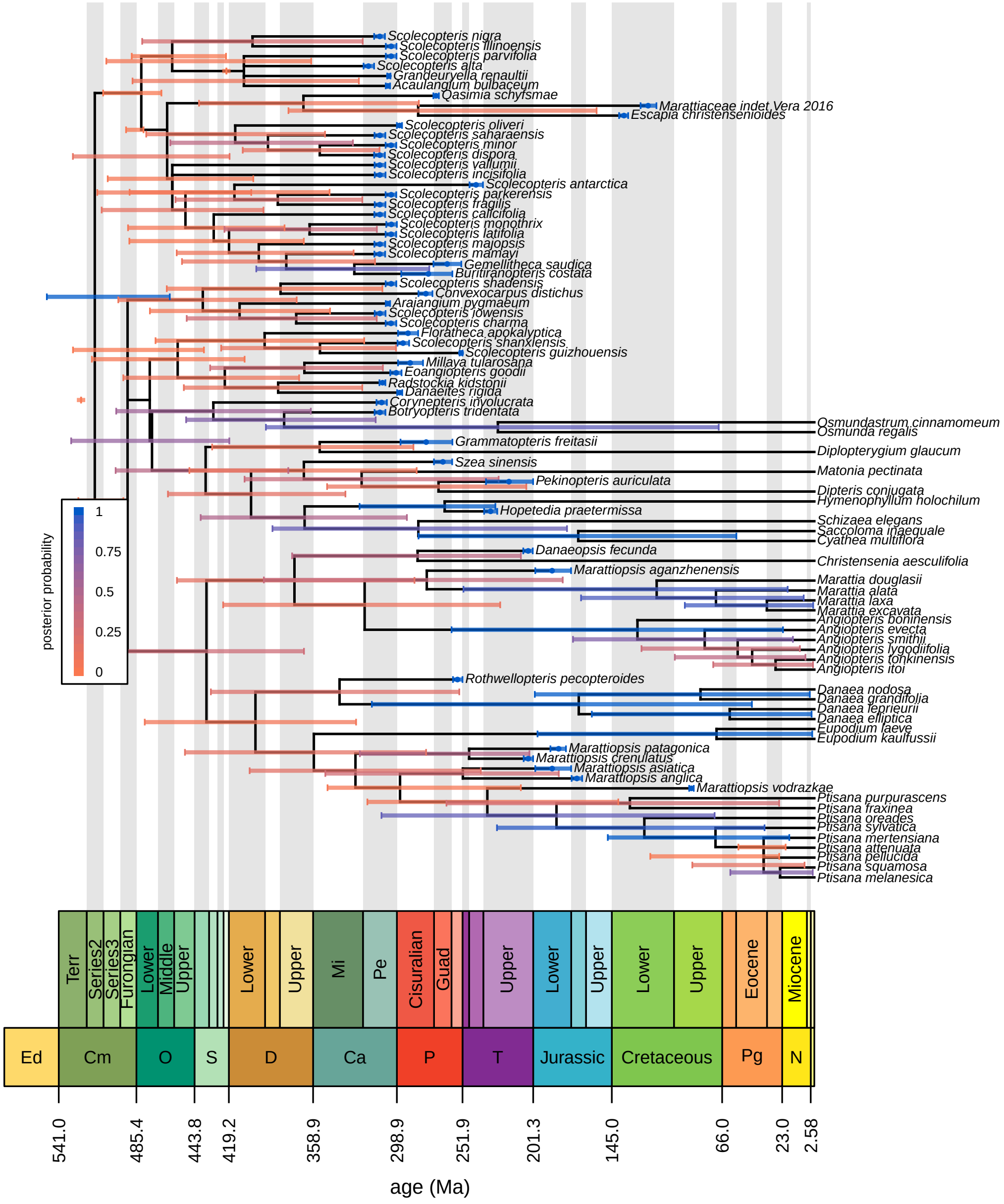

**Figure S49: The maximum clade credibility tree under the uniform tree model.** Bars correspond to the 95% credible interval of clade ages (for internal nodes) and tip ages (for fossils). Numbers on braches and associated age bars correspond to sampled ancestors (key bottom left). Bars are colored in proportion to the posterior probability of the clade for internal nodes, by the probability that the specimen is not a sampled ancestor for tips, and by the probability that the specimen is a sampled ancestor for sampled ancestors (legend, left). Here, we assume the "best" morphological transition and clock models (F81 mixture+Γ and linked, respectively).

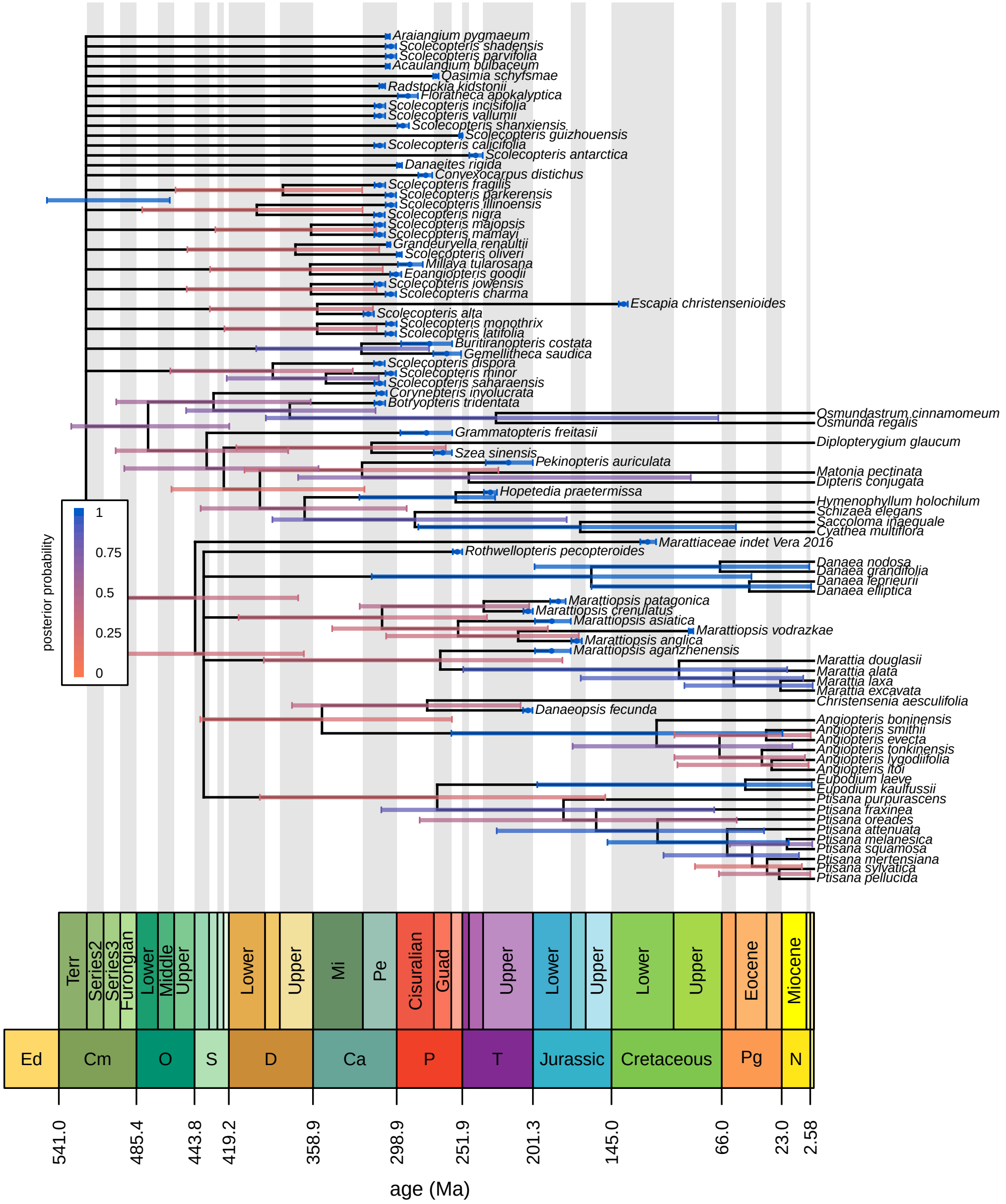


**Figure S50: The 20%-rule consensus tree under the uniform tree model.** Bars correspond to the 95% credible interval of clade ages (for internal nodes) and tip ages (for fossils). Numbers on braches and associated age bars correspond to sampled ancestors (key bottom left). Bars are colored in proportion to the posterior probability of the clade for internal nodes, by the probability that the specimen is not a sampled ancestor for tips, and by the probability that the specimen is a sampled ancestor for sampled ancestors (legend, left). Here, we assume the "best" morphological transition and clock models (F81 mixture+Γ and linked, respectively).

#### S§6.5 Polarized Analysis

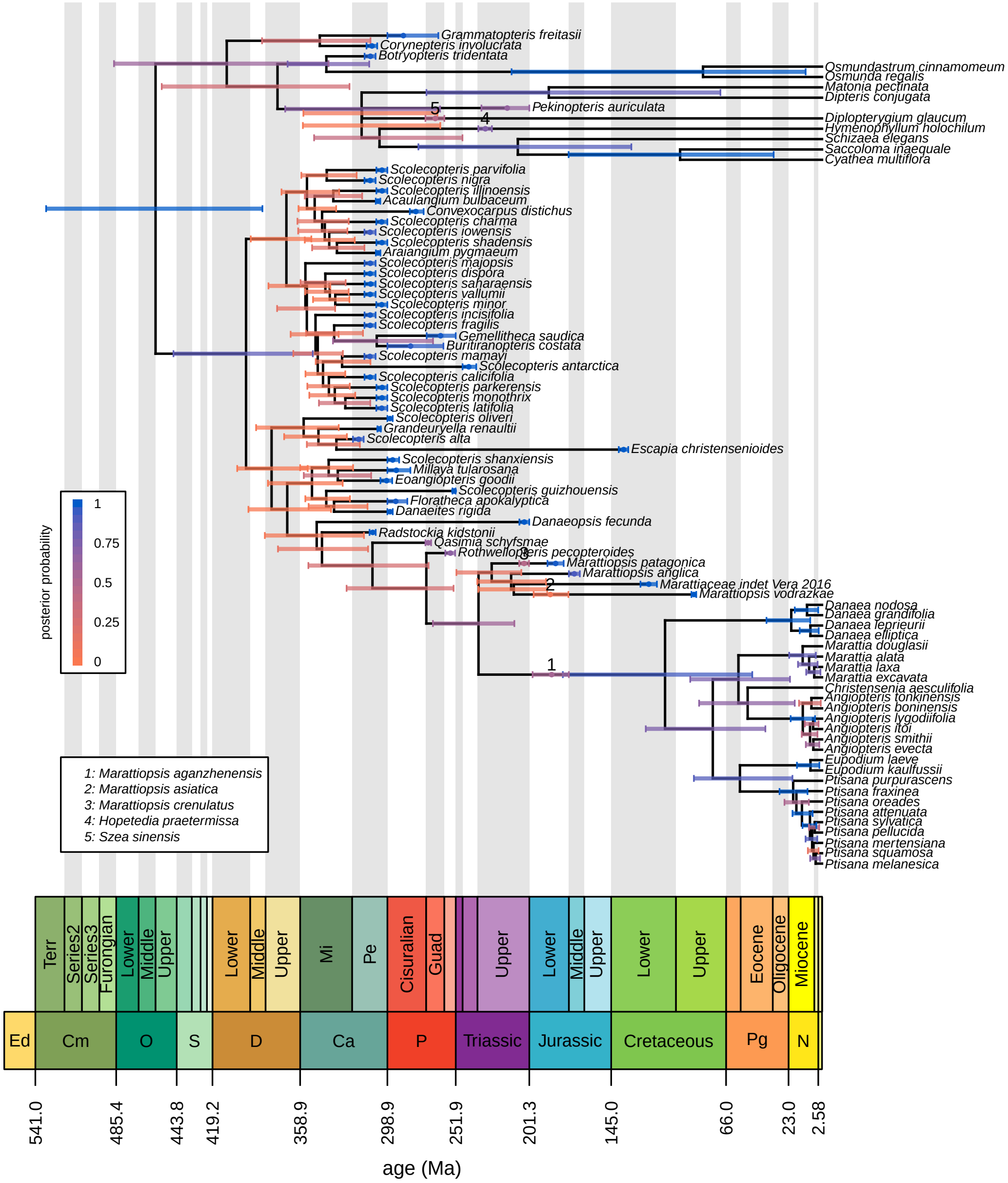


**Figure S51: The maximum clade credibility tree under the preferred model with polarized root.** Bars correspond to the 95% credible interval of clade ages (for internal nodes) and tip ages (for fossils). Numbers on braches and associated age bars correspond to sampled ancestors (key bottom left). Bars are colored in proportion to the posterior probability of the clade for internal nodes, by the probability that the specimen is not a sampled ancestor for tips, and by the probability that the specimen is a sampled ancestor for sampled ancestors (legend, left).

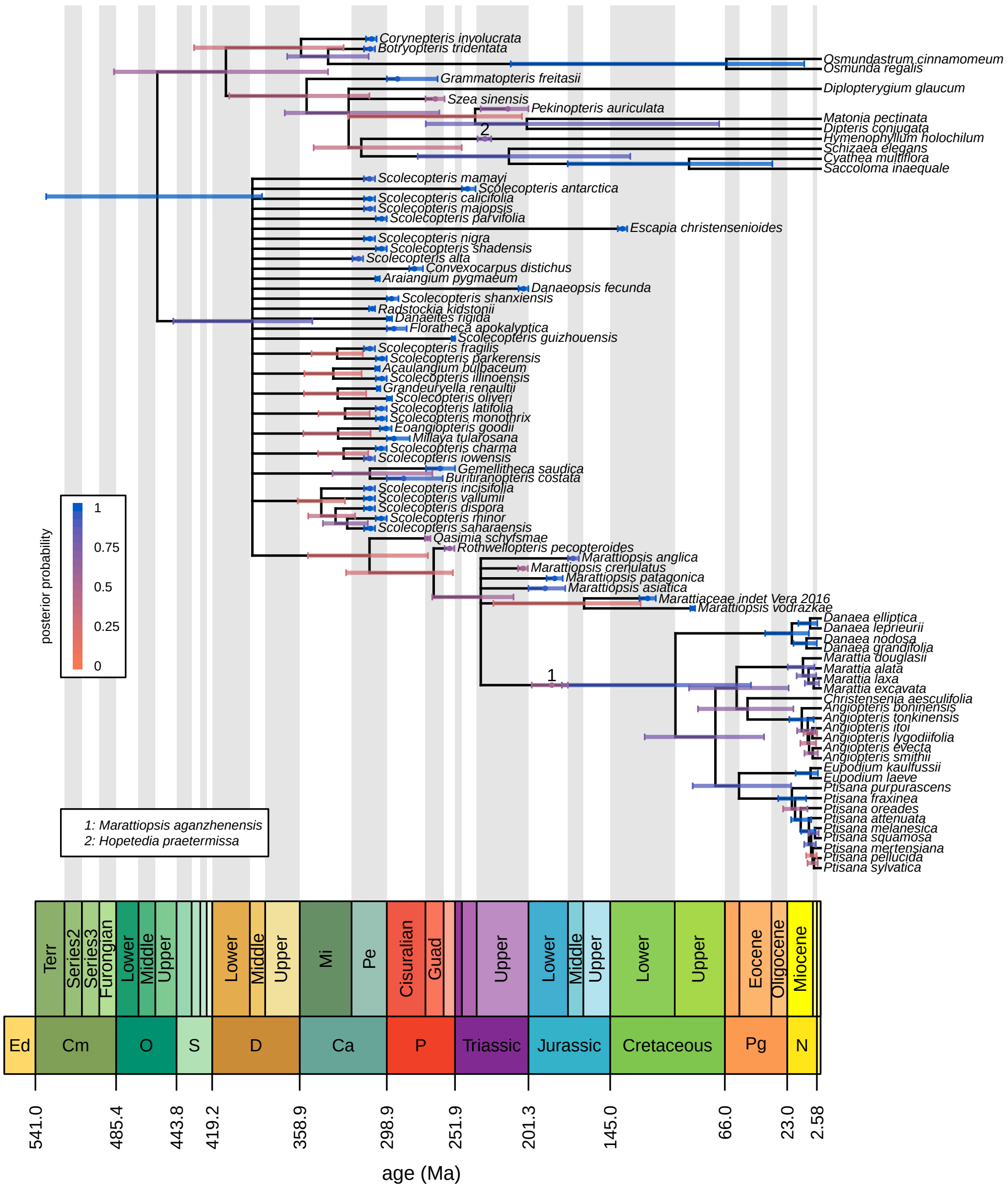


**Figure S52: The 20%-rule consensus tree under the preferred model with polarized root.** Bars correspond to the 95% credible interval of clade ages (for internal nodes) and tip ages (for fossils). Numbers on braches and associated age bars correspond to sampled ancestors (key bottom left). Bars are colored in proportion to the posterior probability of the clade for internal nodes, by the probability that the specimen is not a sampled ancestor for tips, and by the probability that the specimen is a sampled ancestor for sampled ancestors (legend, left).

#### S§6.6 Ancient Plants Analysis

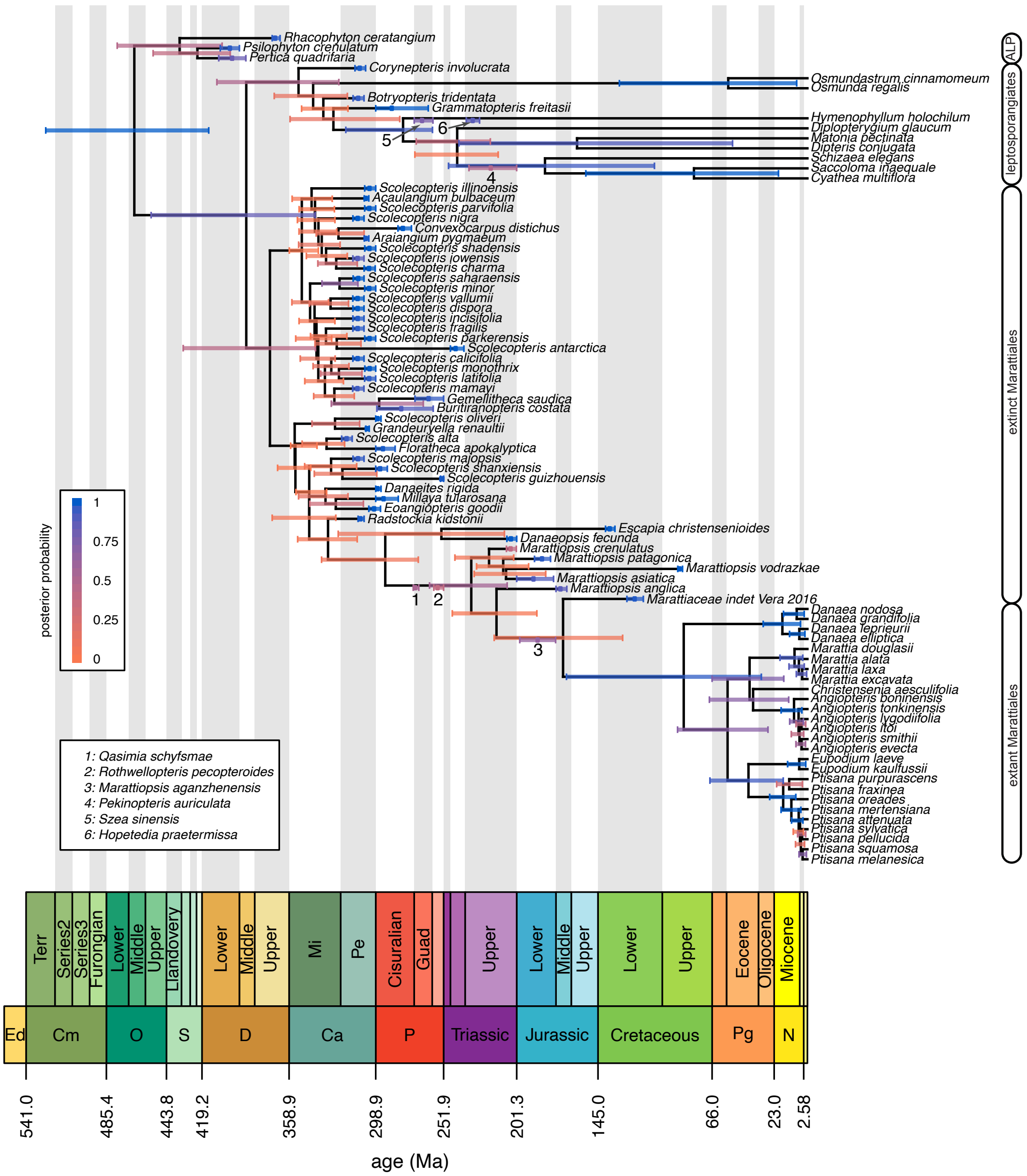


**Figure S53: The maximum clade credibility tree under the preferred model with ancient plant fossils.** Bars correspond to the 95% credible interval of clade ages (for internal nodes) and tip ages (for fossils). Numbers on braches and associated age bars correspond to sampled ancestors (key bottom left). Bars are colored in proportion to the posterior probability of the clade for internal nodes, by the probability that the specimen is not a sampled ancestor for tips, and by the probability that the specimen is a sampled ancestor for sampled ancestors (legend, left).

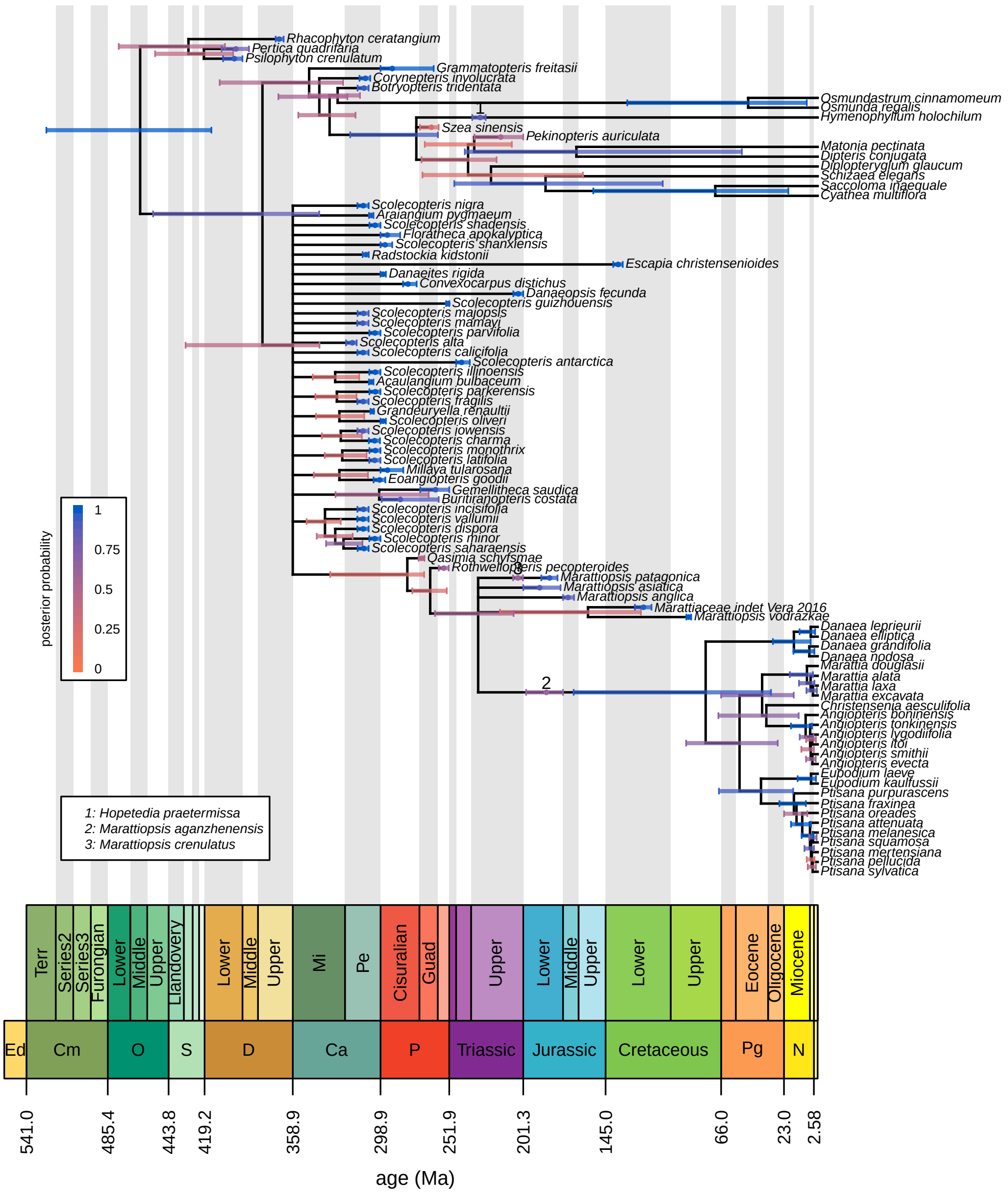


**Figure S54: The 20%-rule consensus tree under the preferred model with ancient land plant fossils.** Bars correspond to the 95% credible interval of clade ages (for internal nodes) and tip ages (for fossils). Numbers on braches and associated age bars correspond to sampled ancestors (key bottom left). Bars are colored in proportion to the posterior probability of the clade for internal nodes, by the probability that the specimen is not a sampled ancestor for tips, and by the probability that the specimen is a sampled ancestor for sampled ancestors (legend, left).

#### S§6.7 Ingroup Analysis

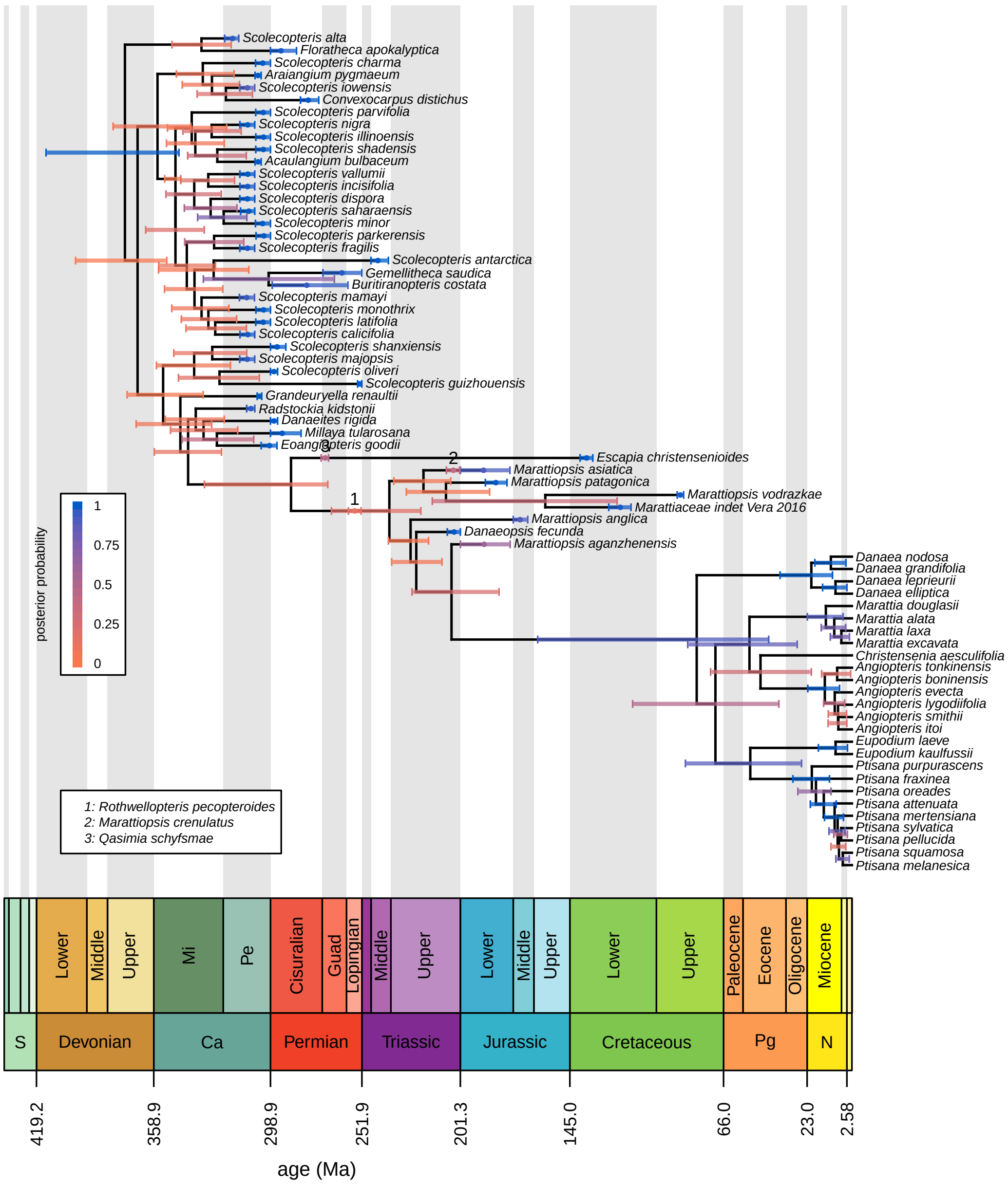


**Figure S55: The maximum clade credibility tree under the preferred model and just ingroup taxa.** Bars correspond to the 95% credible interval of clade ages (for internal nodes) and tip ages (for fossils). Numbers on branches and associated age bars correspond to sampled ancestors (key bottom left). Bars are colored in proportion to the posterior probability of the clade for internal nodes, by the probability that the specimen is not a sampled ancestor for tips, and by the probability that the specimen is a sampled ancestor for sampled ancestors (legend, left).

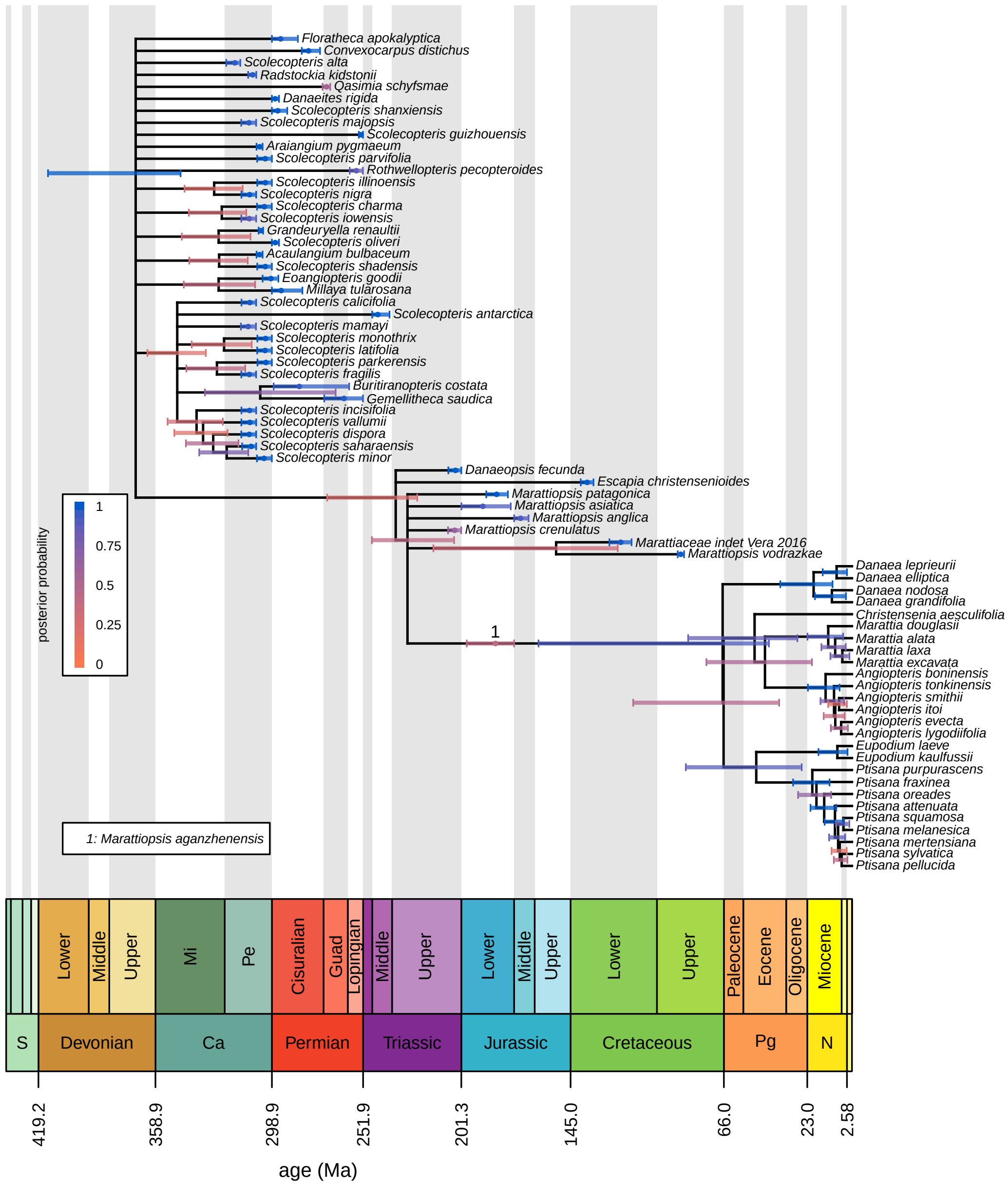


**Figure S56: The 20%-rule consensus tree under the preferred model and just ingroup taxa.** Bars correspond to the 95% credible interval of clade ages (for internal nodes) and tip ages (for fossils). Numbers on braches and associated age bars correspond to sampled ancestors (key bottom left). Bars are colored in proportion to the posterior probability of the clade for internal nodes, by the probability that the specimen is not a sampled ancestor for tips, and by the probability that the specimen is a sampled ancestor for sampled ancestors (legend, left).

#### S§6.8 Comparing Empirical Considerations

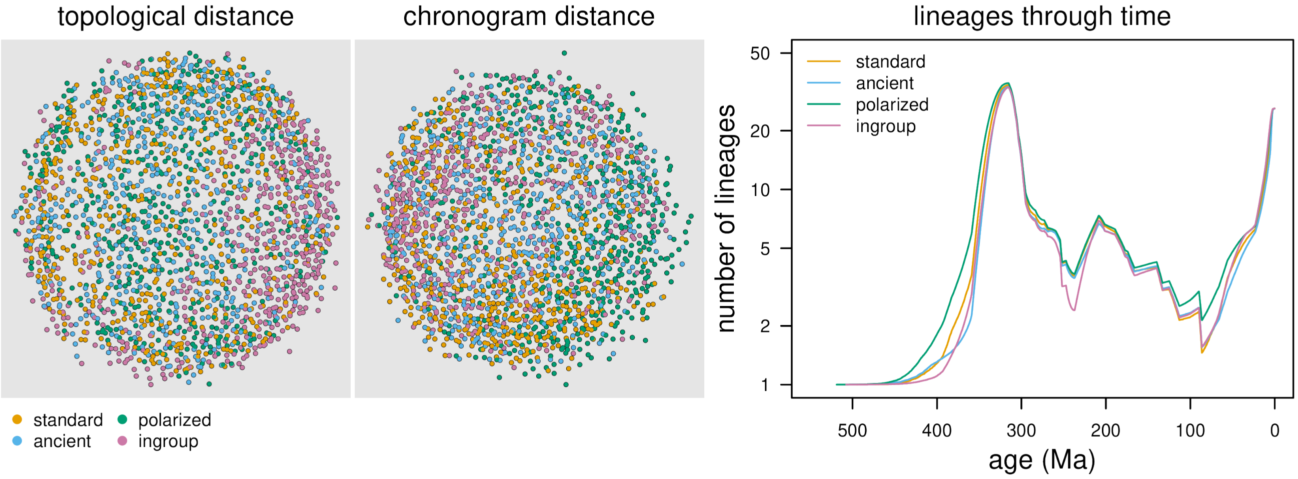


**Figure S57: The impact of empirical considerations on phylogenetic estimates.** We compute the RF distances and KF distances between chronograms among our "empirical considerations" analyses and the "standard" dataset under the preferred models (left and middle, respectively). We only compare samples where the ingroup is inferred to be monophyletic, and pruned outgroup taxa before computing pairwise distances. We also computed the LTT curves for the ingroup taxa for the same sets of analyses, again conditioning on the monophyly of the ingroup and pruning outgroup taxa (right). We compare the ages of particular clades in Fig. S59.

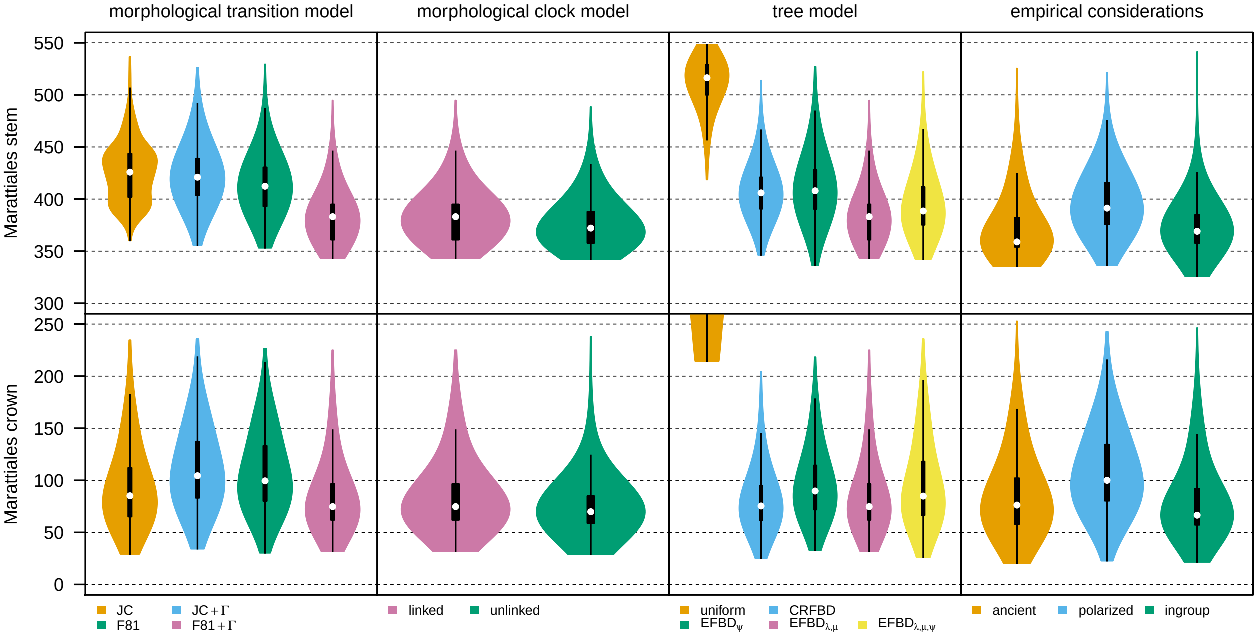


**Figure S58: The impact of modeling and empirical considerations on divergence-time estimates of major clades.** We plot the marginal posterior distribution of the age of each node (in rows) as a function of the morphological-transition model (column 1), the morphological clock model (column 2), the tree model (column 3), and the empirical considerations (column 4). White dots are the posterior median ages, and black bars correspond to the 50% (thick) and 95% (narrow) credible intervals. We report means, medians, and 95% CIs in the Supplemental Material (Tables S.3 and S.4)

**Table S.3: Age of stem Marattiales under various models and empirical consideration.**

| Dataset | Transition model | Clock model | Tree model | Mean | Median | 95% CI |
|---|---|---|---|---|---|---|
| standard | Mk | linked | $EFBD_{\lambda,\mu}$ | 455.79 | 452.88 | [392.17 – 520.74] |
| standard | Mk+Γ | linked | $EFBD_{\lambda,\mu}$ | 470.22 | 470.38 | [399.28 – 533.04] |
| standard | F81 | linked | $EFBD_{\lambda,\mu}$ | 465.34 | 465.23 | [392.78 – 529.96] |
| standard | F81+Γ | linked | $EFBD_{\lambda,\mu}$ | 428.87 | 427.26 | [366.98 – 506.83] |
| standard | F81+Γ | unlinked | $EFBD_{\lambda,\mu}$ | 409.14 | 406.43 | [355.79 – 483.87] |
| standard | F81+Γ | linked | uniform | 520.74 | 527.45 | [460.15 – 547.77] |
| standard | F81+Γ | linked | CRFBD | 441.42 | 439.69 | [391.07 – 505.03] |
| standard | F81+Γ | linked | $EFBD_{\psi}$ | 457.70 | 457.03 | [395.20 – 523.94] |
| standard | F81+Γ | linked | $EFBD_{\lambda,\mu,\psi}$ | 443.84 | 439.29 | [381.55 – 523.05] |
| ancient | F81+Γ | linked | $EFBD_{\lambda,\mu}$ | 414.84 | 413.80 | [350.90 – 505.74] |
| polarized | F81+Γ | linked | $EFBD_{\lambda,\mu}$ | 455.02 | 453.48 | [383.54 – 530.10] |

**Table S.4: Age of crown Marattiales under various models and empirical consideration.**

| Dataset | Transition model | Clock model | Tree model | Mean | Median | 95% CI |
|---|---|---|---|---|---|---|
| standard | Mk | linked | $EFBD_{\lambda,\mu}$ | 93.80 | 85.15 | [46.99 – 179.36] |
| standard | Mk+Γ | linked | $EFBD_{\lambda,\mu}$ | 110.99 | 104.39 | [54.06 – 186.06] |
| standard | F81 | linked | $EFBD_{\lambda,\mu}$ | 107.68 | 99.48 | [54.78 – 181.12] |
| standard | F81+Γ | linked | $EFBD_{\lambda,\mu}$ | 83.92 | 74.74 | [46.32 – 169.07] |
| standard | F81+Γ | unlinked | $EFBD_{\lambda,\mu}$ | 74.85 | 69.84 | [44.76 – 139.55] |
| standard | F81+Γ | linked | $EFBD_{\lambda,\mu}$ | 83.92 | 74.74 | [46.32 – 169.07] |
| standard | F81+Γ | linked | uniform | 322.65 | 313.39 | [214.69 – 426.96] |
| standard | F81+Γ | linked | CRFBD | 80.95 | 75.38 | [42.91 – 151.43] |
| standard | F81+Γ | linked | $EFBD_{\psi}$ | 95.91 | 89.73 | [49.67 – 173.27] |
| standard | F81+Γ | linked | $EFBD_{\lambda,\mu,\psi}$ | 95.79 | 84.85 | [45.51 – 185.14] |
| ancient | F81+Γ | linked | $EFBD_{\lambda,\mu}$ | 85.50 | 76.21 | [37.44 – 173.74] |
| polarized | F81+Γ | linked | $EFBD_{\lambda,\mu}$ | 107.97 | 100.00 | [52.31 – 188.56] |
| ingroup | F81+Γ | linked | $EFBD_{\lambda,\mu}$ | 79.87 | 66.51 | [45.11 – 168.21] |

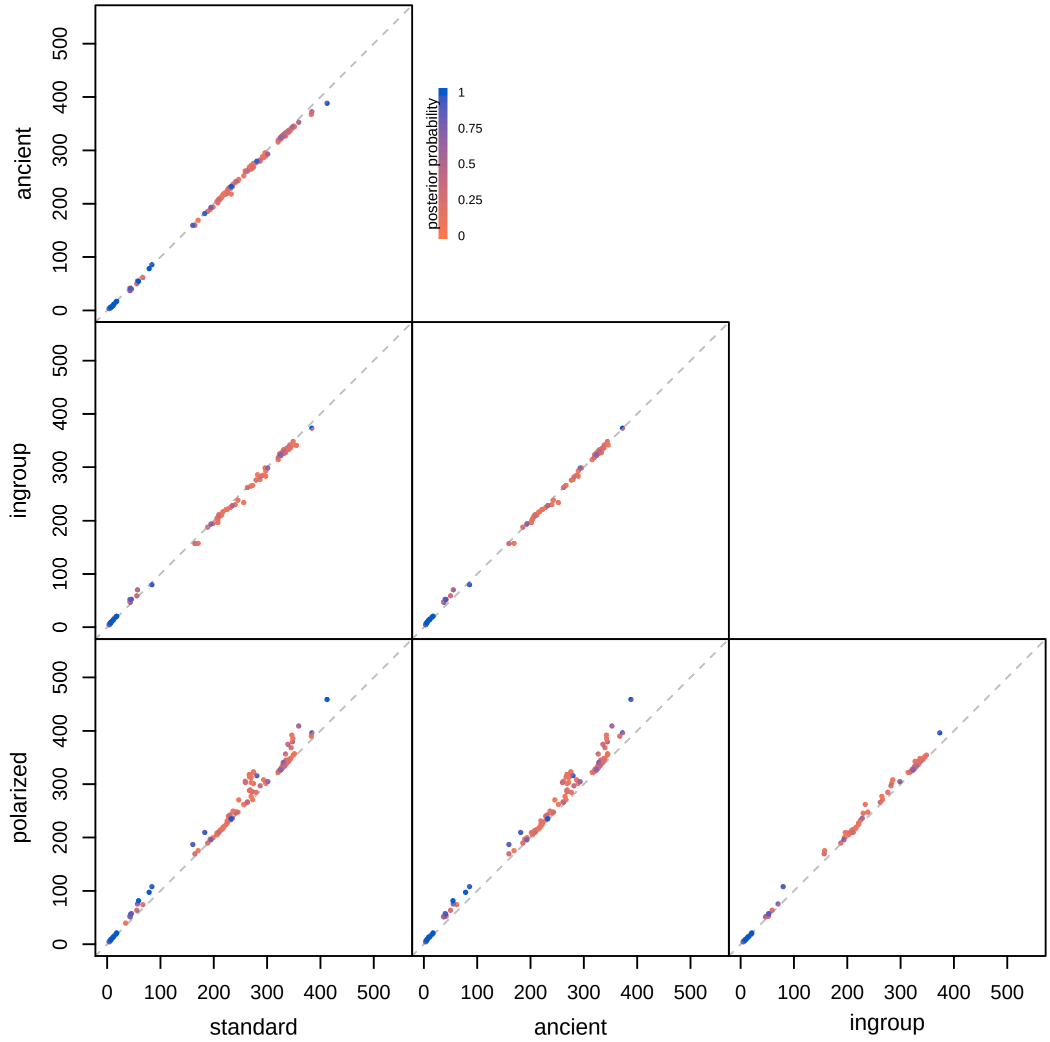


**Figure S59: Comparing divergence-time estimates for each clade with different empirical considerations.** For each pair of models, we compare the posterior-mean age of each clade with posterior probability $> 0.05$ under the analysis on the x-axis against the analysis on the y-axis. Each point is divided in half, with the top-left semicircle colored by the posterior probability under the model on the y-axis and the bottom-right semicircle colored by the posterior probability under the model on the x-axis. In these comparisons, we assume the preferred morphological transition and clock models and tree model (F81 mixture+$\Gamma$, linked, $\text{EFBD}_{\lambda,\mu}$ respectively).

**Table S.5: Delay before appearance of crown Marattiales under various models and empirical considerations.** Specifically, we calculate the difference between the age of the MRCA of extant Marattiales (conditional on it being monophyletic, *i.e.*,, it does not include extinct taxa) and the age of the node from which it descends (the most recent common ancestor with another sample). We do not report this difference under the uniform model because the crown Marattiales never exclude fossils.

| Dataset | Transition model | Clock model | Tree model | Mean | Median | 95% CI |
|---|---|---|---|---|---|---|
| standard | Mk | linked | $EFBD_{\lambda,\mu}$ | 100.36 | 106.44 | [12.66 – 162.64] |
| standard | Mk+$\Gamma$ | linked | $EFBD_{\lambda,\mu}$ | 81.35 | 85.14 | [5.75 – 148.87] |
| standard | F81 | linked | $EFBD_{\lambda,\mu}$ | 86.72 | 91.56 | [10.17 – 151.81] |
| standard | F81+$\Gamma$ | linked | $EFBD_{\lambda,\mu}$ | 111.12 | 116.35 | [18.29 – 169.06] |
| standard | F81+$\Gamma$ | unlinked | $EFBD_{\lambda,\mu}$ | 130.47 | 134.53 | [45.82 – 187.93] |
| standard | F81+$\Gamma$ | unlinked | $EFBD_{\lambda,\mu}$ | 130.47 | 134.53 | [45.82 – 187.93] |
| standard | F81+$\Gamma$ | linked | $EFBD_{\psi}$ | 92.81 | 96.99 | [13.93 – 156.50] |
| standard | F81+$\Gamma$ | linked | CRFBD | 105.22 | 110.37 | [27.89 – 159.10] |
| standard | F81+$\Gamma$ | linked | $EFBD_{\lambda,\mu,\psi}$ | 98.02 | 106.45 | [7.38 – 165.26] |
| ancient | F81+$\Gamma$ | linked | $EFBD_{\lambda,\mu}$ | 107.42 | 114.57 | [12.54 – 172.42] |
| polarized | F81+$\Gamma$ | linked | $EFBD_{\lambda,\mu}$ | 87.77 | 91.49 | [8.04 – 157.08] |

#### S§6.9 Extant Phylogenies

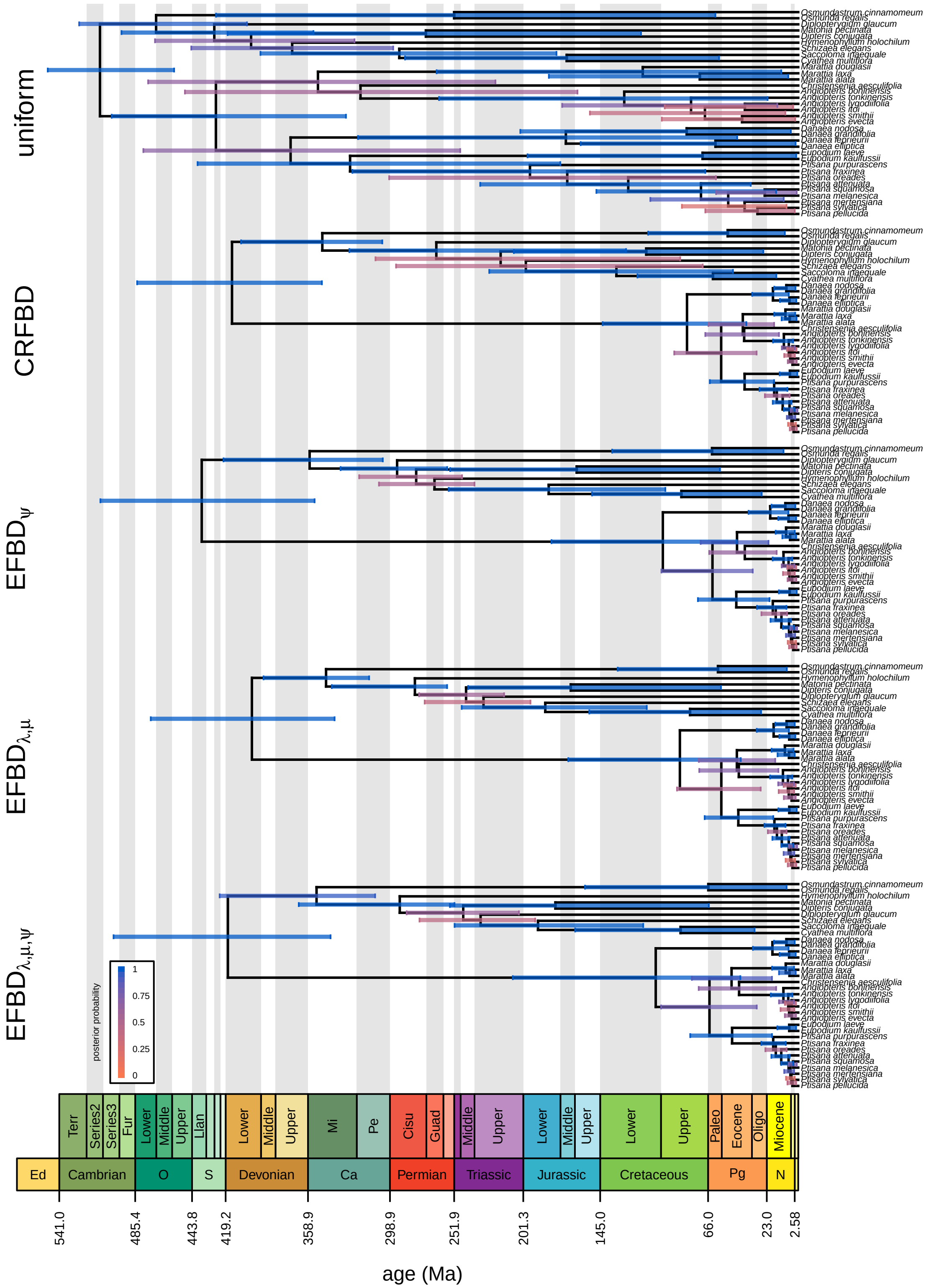


**Figure S60: The maximum clade credibility trees for extant taxa among tree models.** Bars correspond to the 95% credible interval of clade ages. Bars are colored in proportion to the posterior probability of the clade (legend, bottom left). In these comparisons, we assume the preferred morphological transition and clock models (F81+$\Gamma$ and linked, respectively).

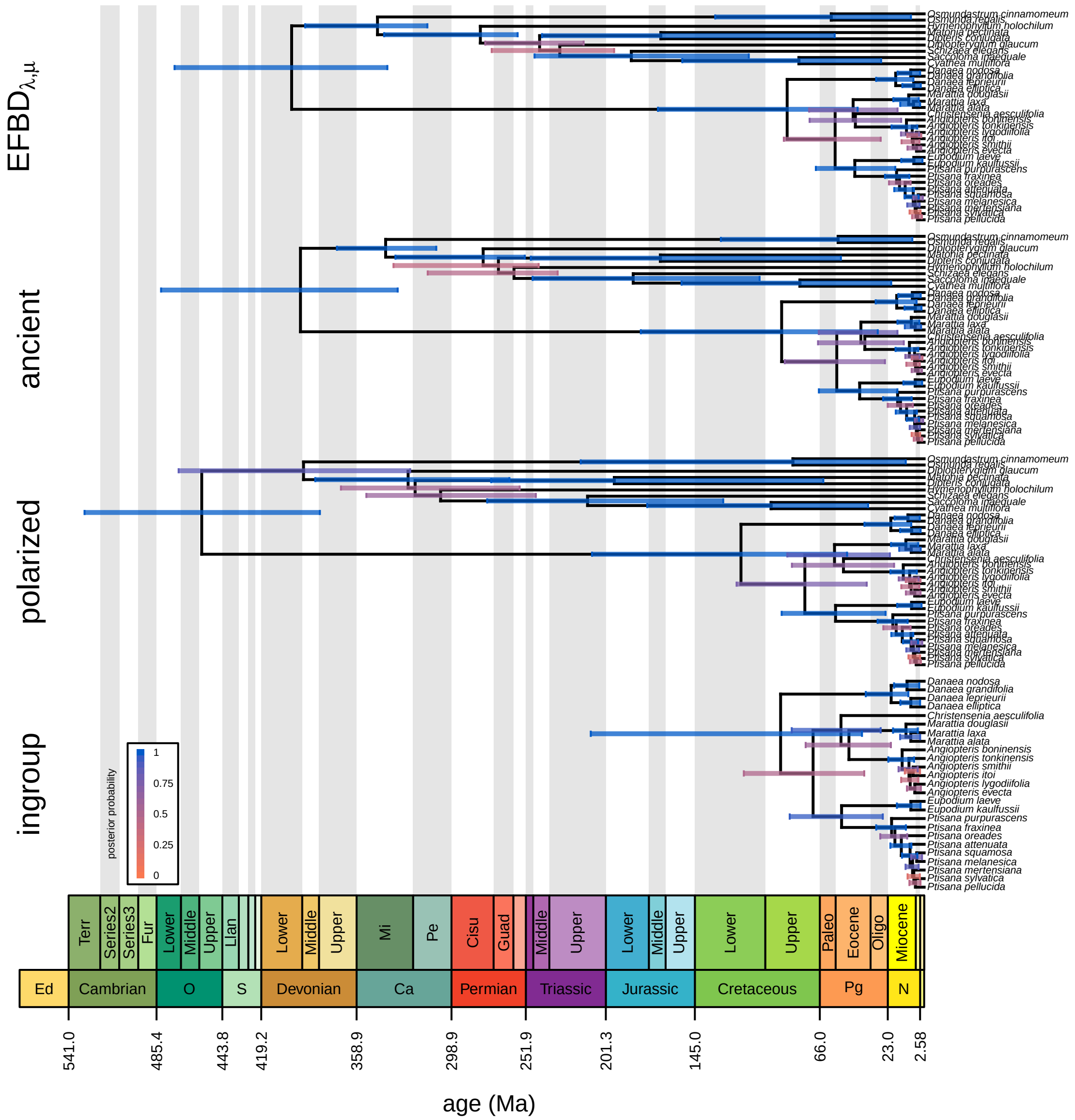


**Figure S61: The maximum clade credibility trees for extant taxa among empirical datasets.** Bars correspond to the 95% credible interval of clade ages. Bars are colored in proportion to the posterior probability of the clade (legend, bottom left). In these comparisons, we assume the preferred morphological transition and clock models and tree model (F81 mixture+Γ, linked, EFBD$_{\lambda,\mu}$ respectively).

#### S§6.10 Stochastic Maps

We simulated stochastic maps on maximum clade credibility trees using posterior-mean estimates for all relevant parameters under the preferred models (the F81 mixture+$\Gamma$ morphological-transition model, the linked morphological-clock model, and the $\text{EFBD}_{\lambda,\mu}$ tree model). Here, we report stochastic maps for characters of interest (*i.e.*, those that we interpret biologically in the main text) under the ancient plants analyses. We provide the stochastic maps for the remaining characters and for the standard dataset under the preferred model in the Supplemental Archive (DRYAD XXXX).

In each of the following stochastic maps, we assign a unique color to each character state. The color at a given time on a given branch is the weighted average of the state-specific colors, weighted by the posterior probability that the character is in each state. Pie charts at the tips of the tree represent the raw data, not the posterior estimate of the state under the model (*i.e.*, as implied by the stochastic map); pies with multiple colors therefore reflect partial or complete missing data. The legend indicates the labels for each state and their associated color, and the axis is the age in millions of years (Ma).

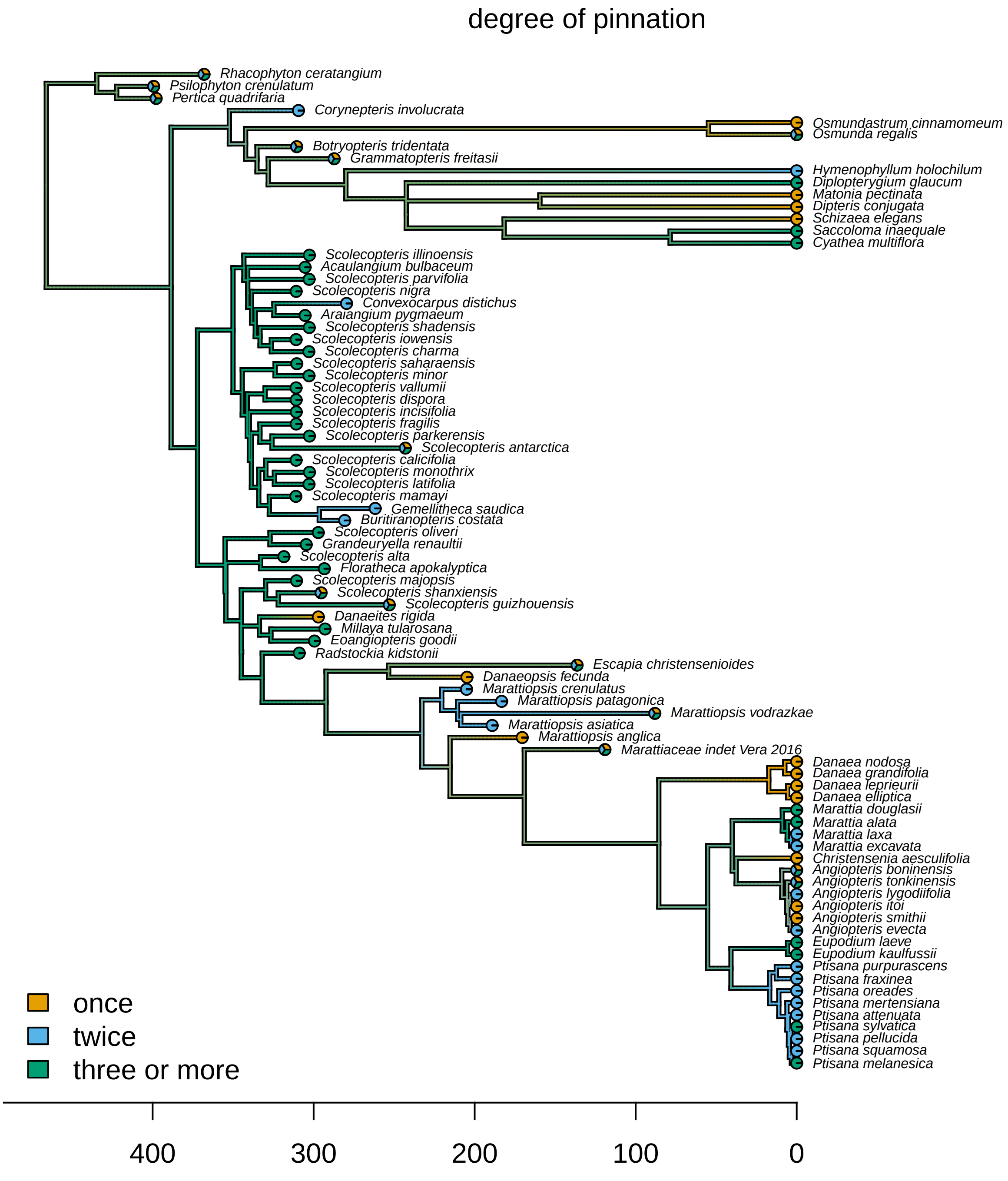


**Figure S62: Stochastic map for character 23: degree of pinnation.**

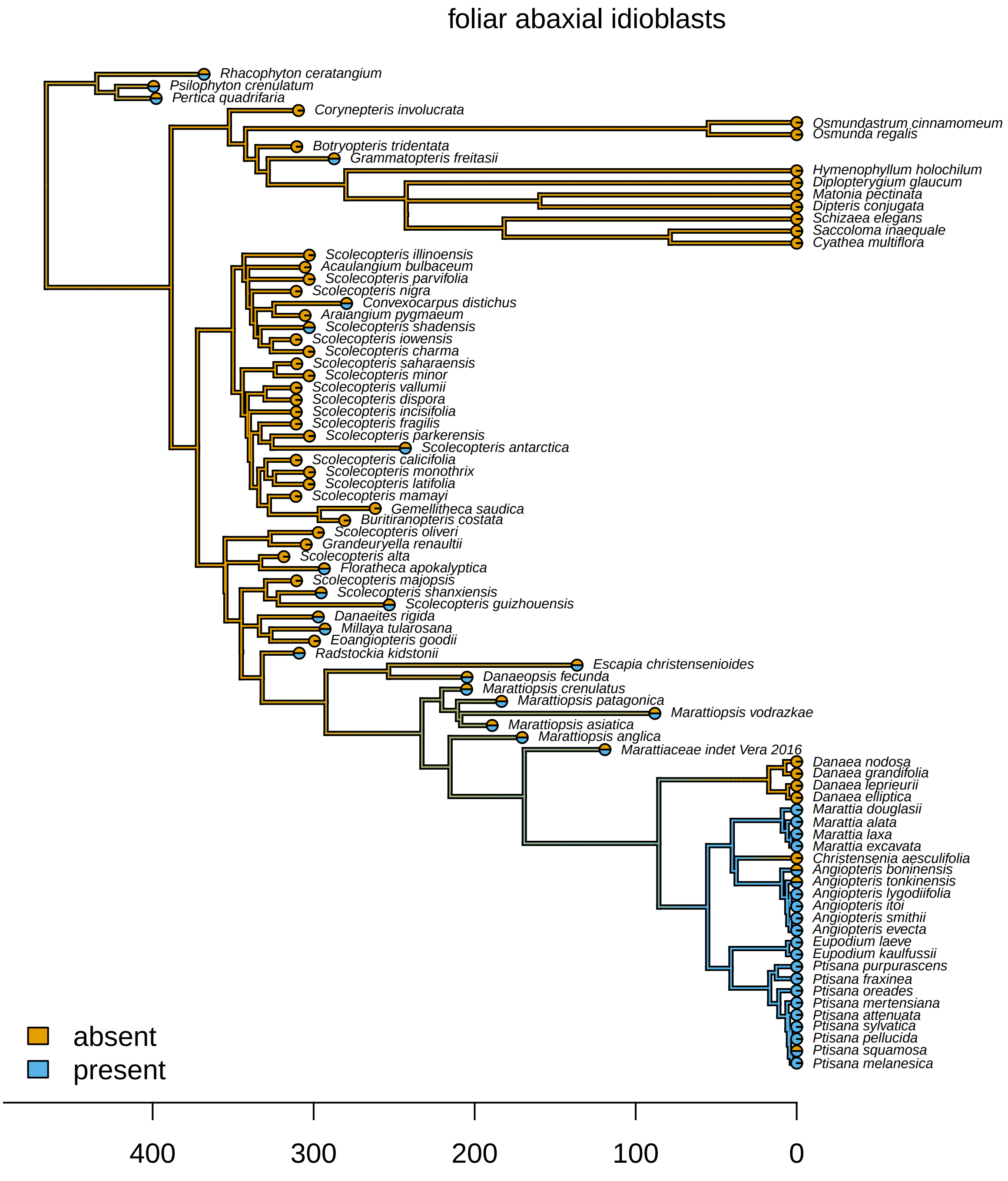


**Figure S63: Stochastic map for character 28: foliar abaxial idioblasts.**

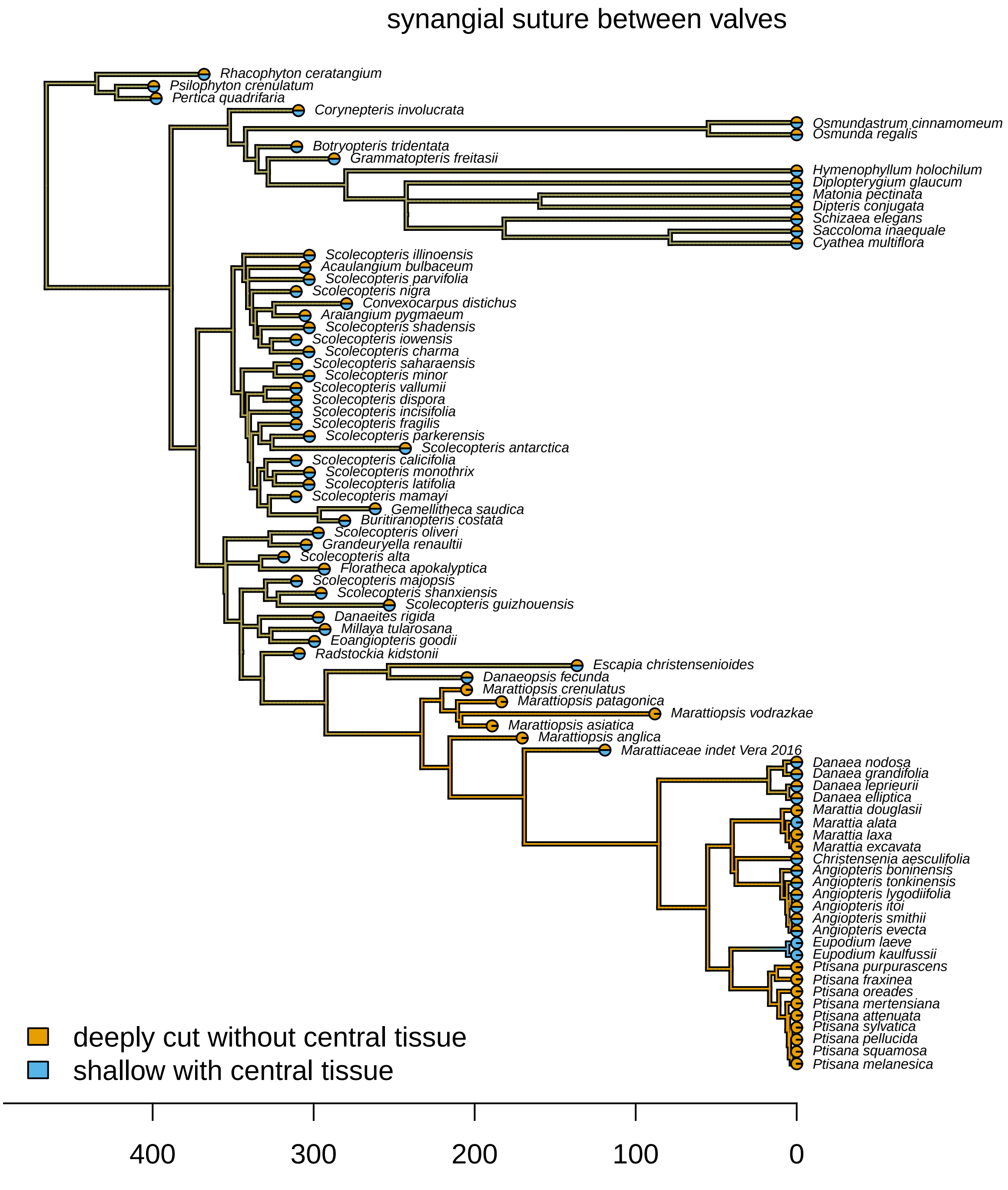


**Figure S64: Stochastic map for character 52: synangial suture between valves.**

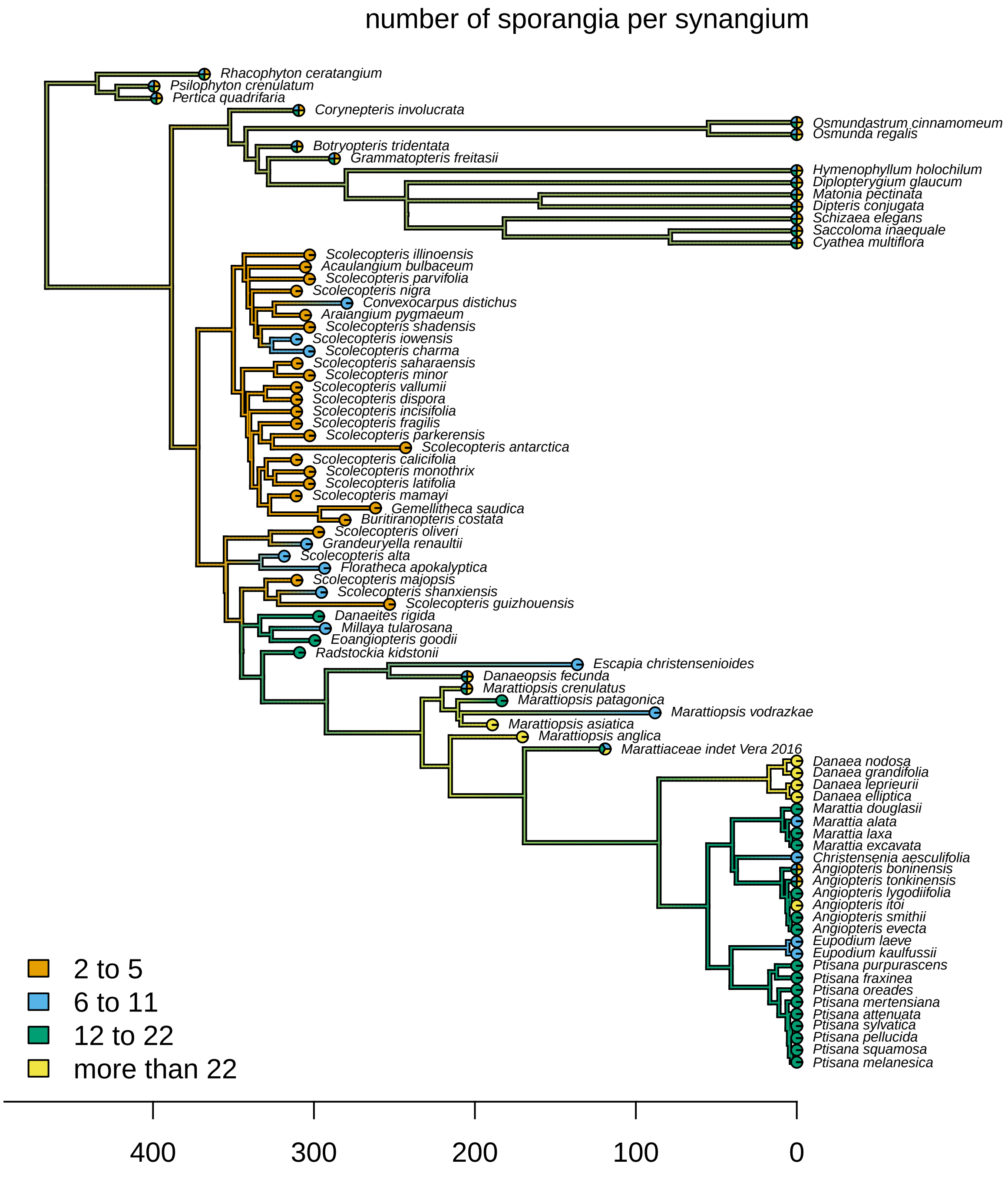


**Figure S65: Stochastic map for character 54: number of sporangia per synangium.**

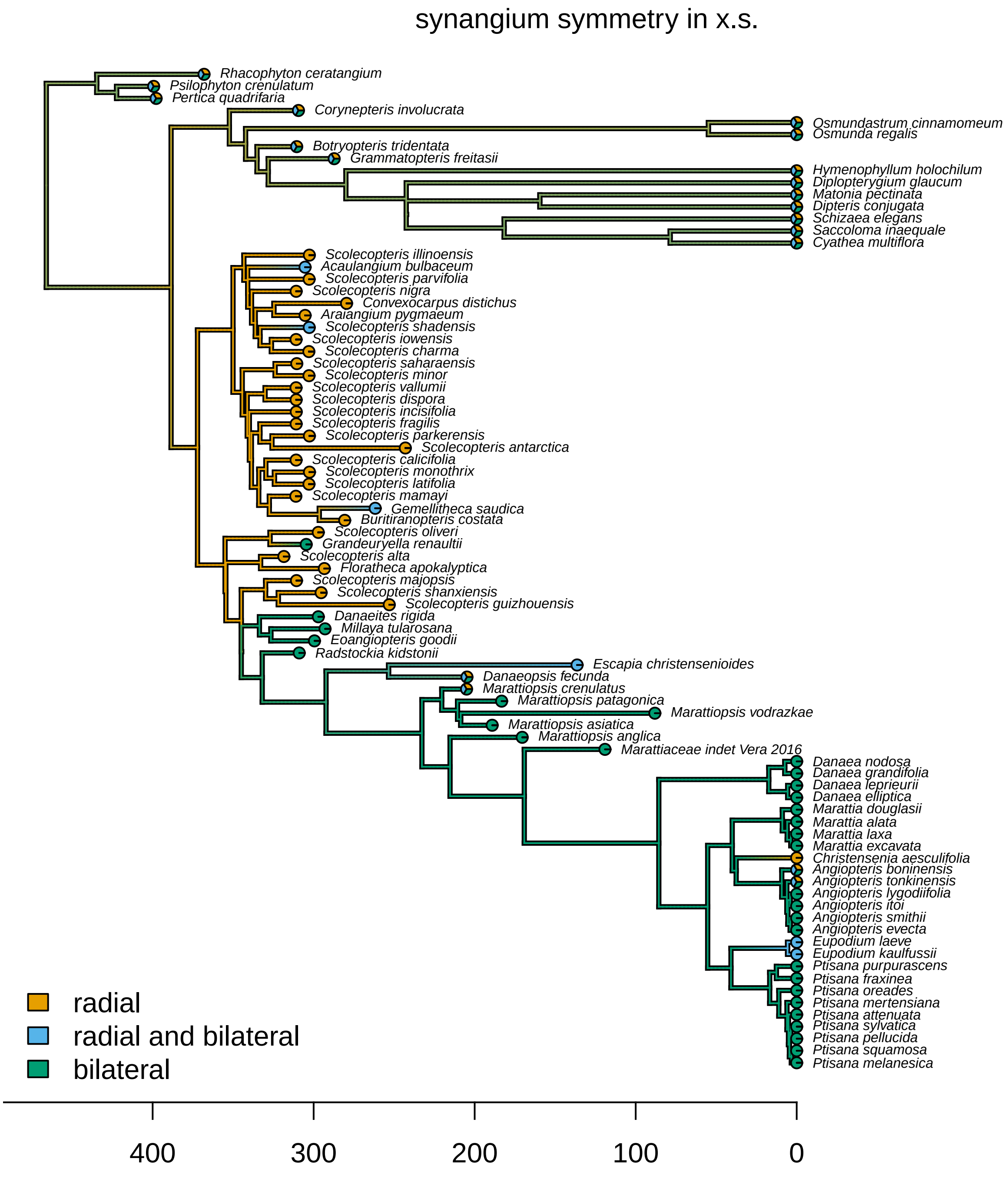


**Figure S66: Stochastic map for character 55: synangium symmetry in x.s.**

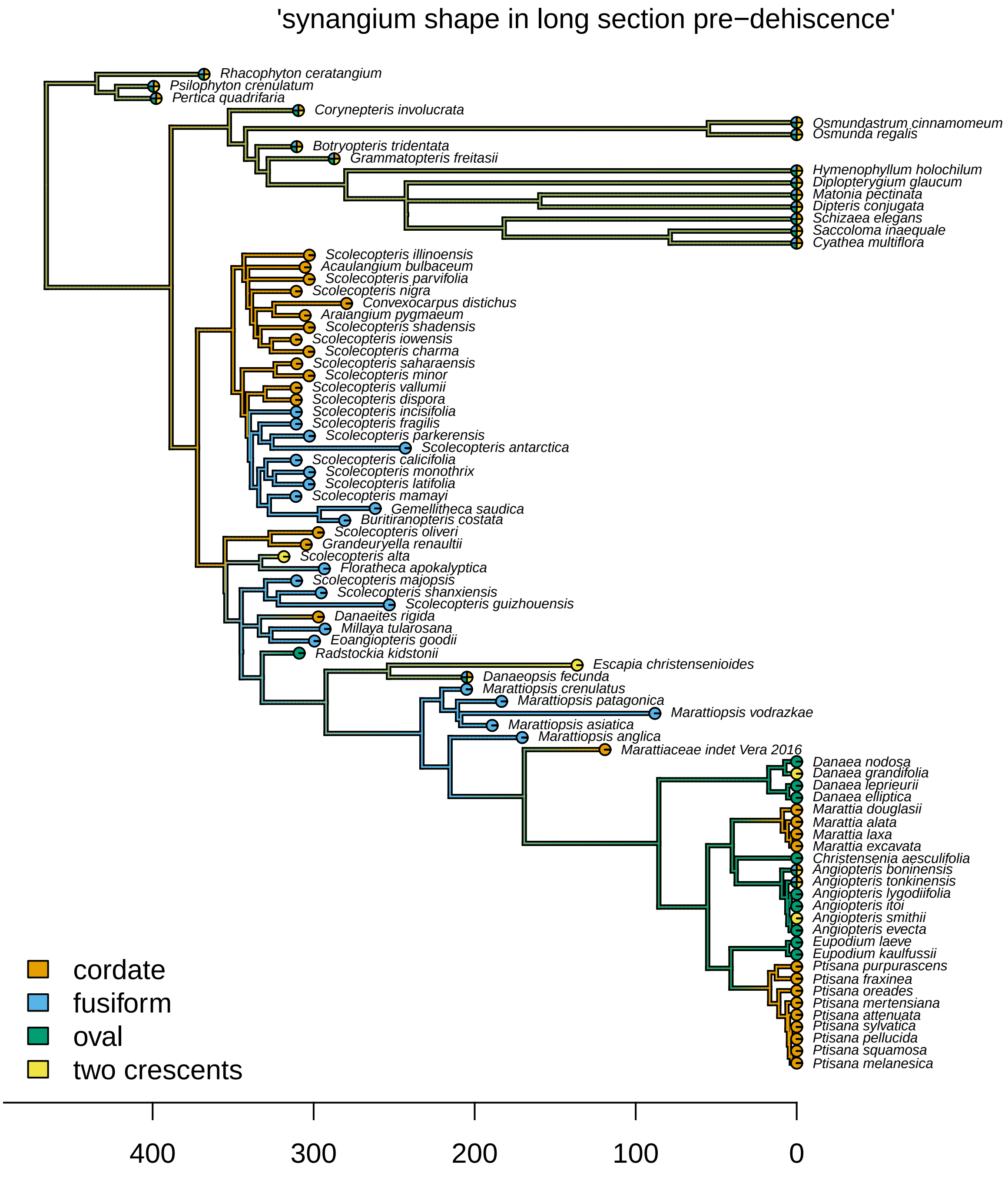


**Figure S67: Stochastic map for character 57: synangium shape in long section pre-dehiscence.**

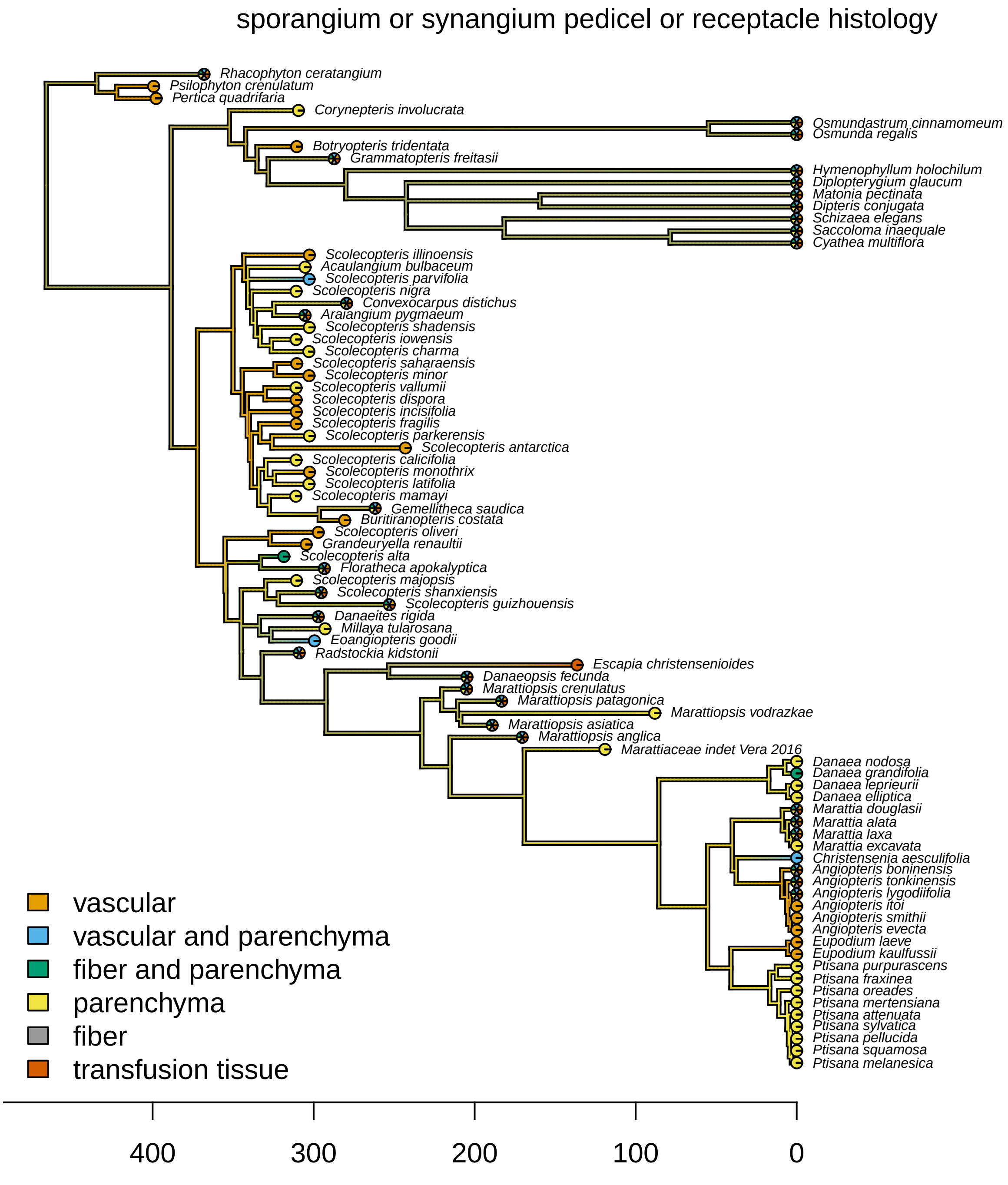


**Figure S68: Stochastic map for character 58: sporangium or synangium pedicel or receptacle histology.**

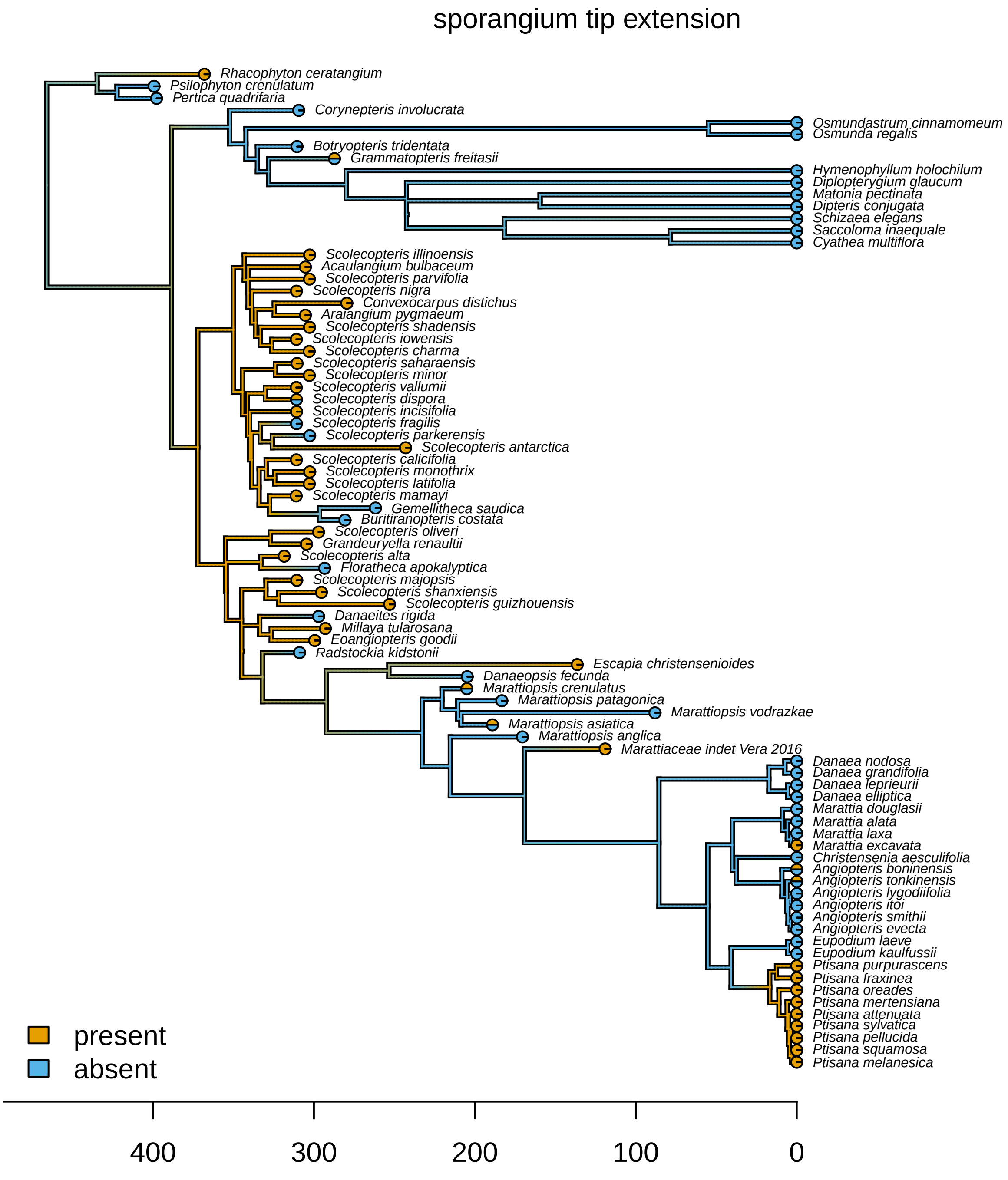


**Figure S69: Stochastic map for character 59: sporangium tip extension.**

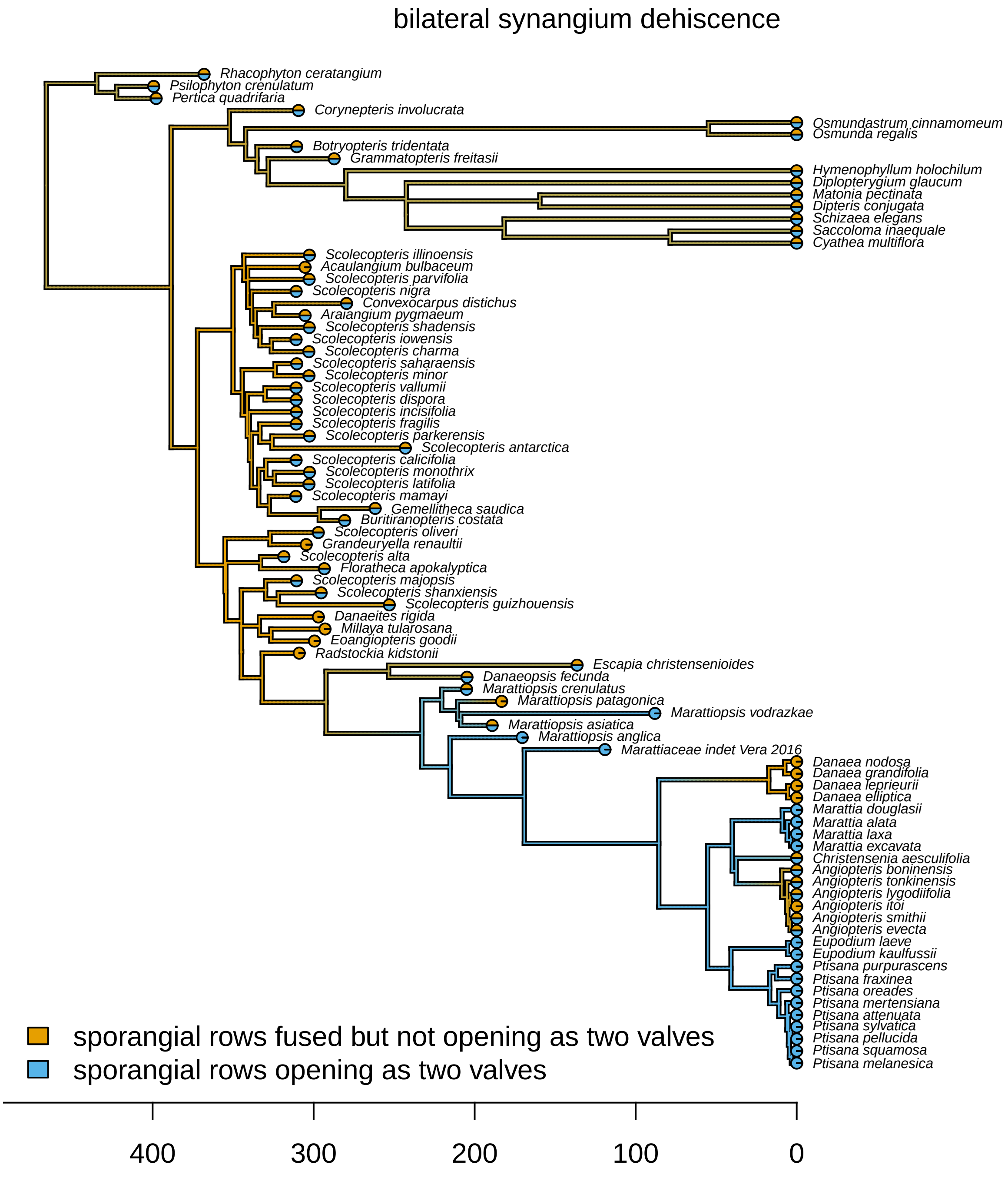


**Figure S70: Stochastic map for character 62: bilateral synangium dehiscence.**

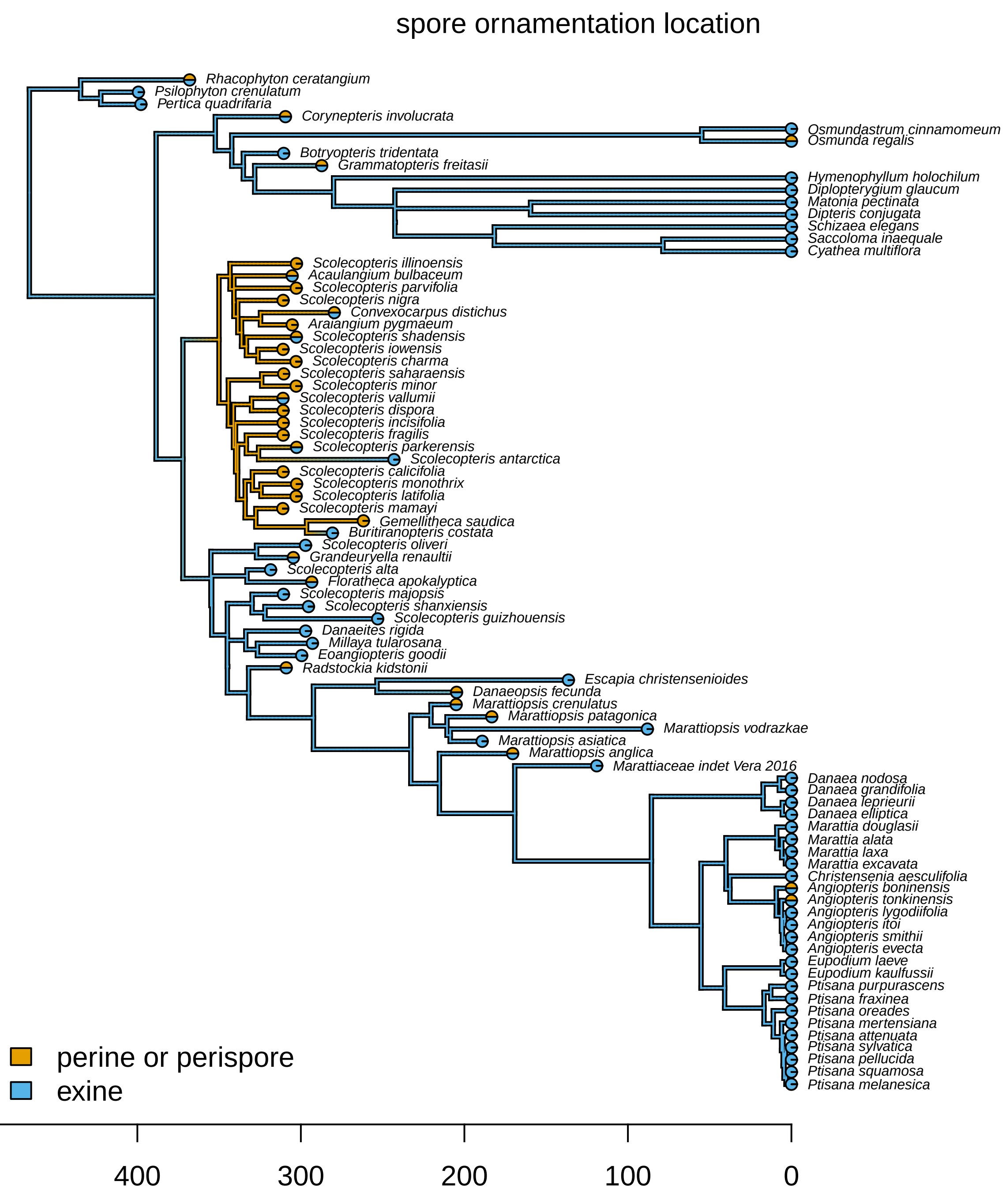


**Figure S71: Stochastic map for character 70: spore ornamentation location.**

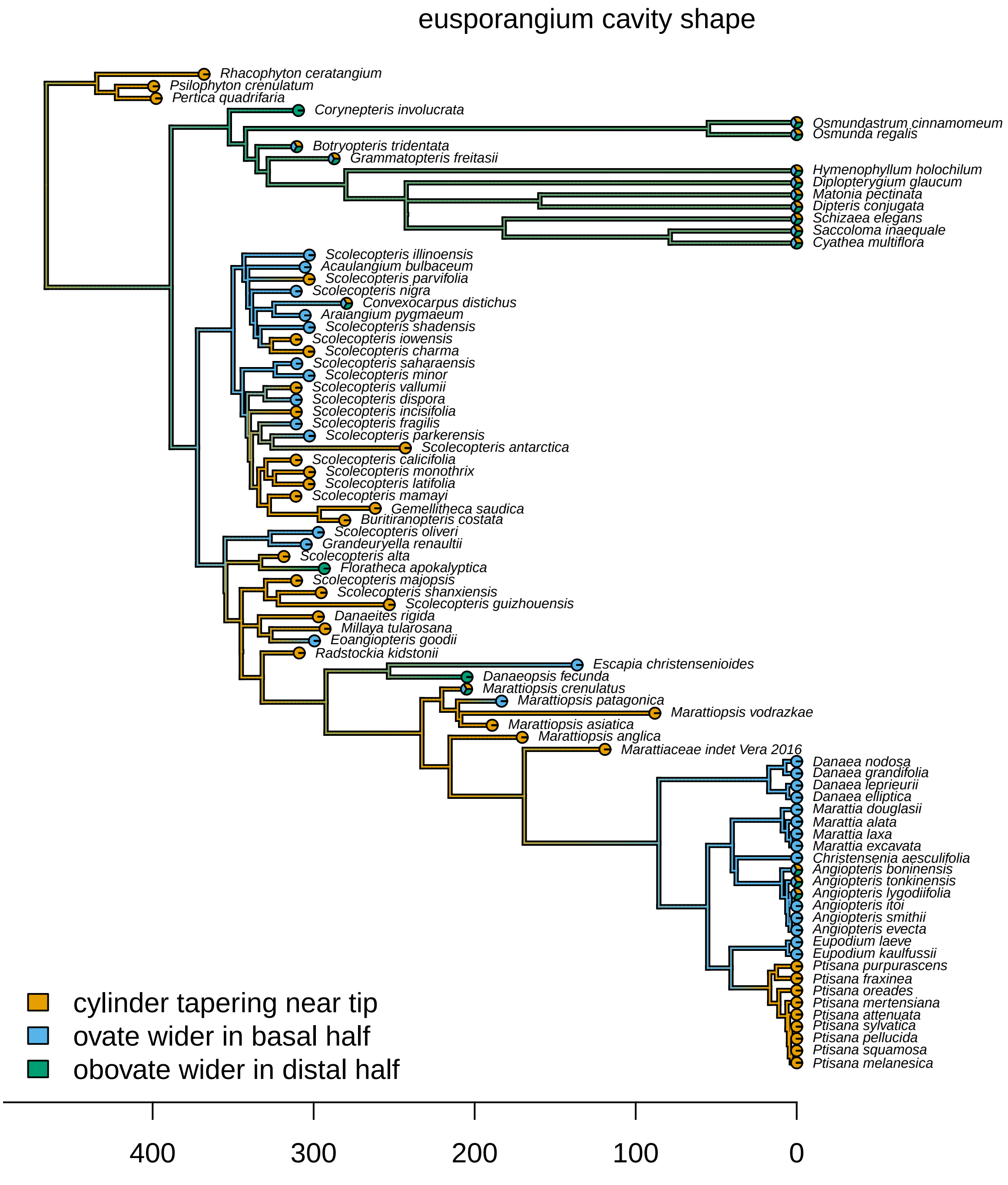


**Figure S72: Stochastic map for character 75: eusporangium cavity shape.**

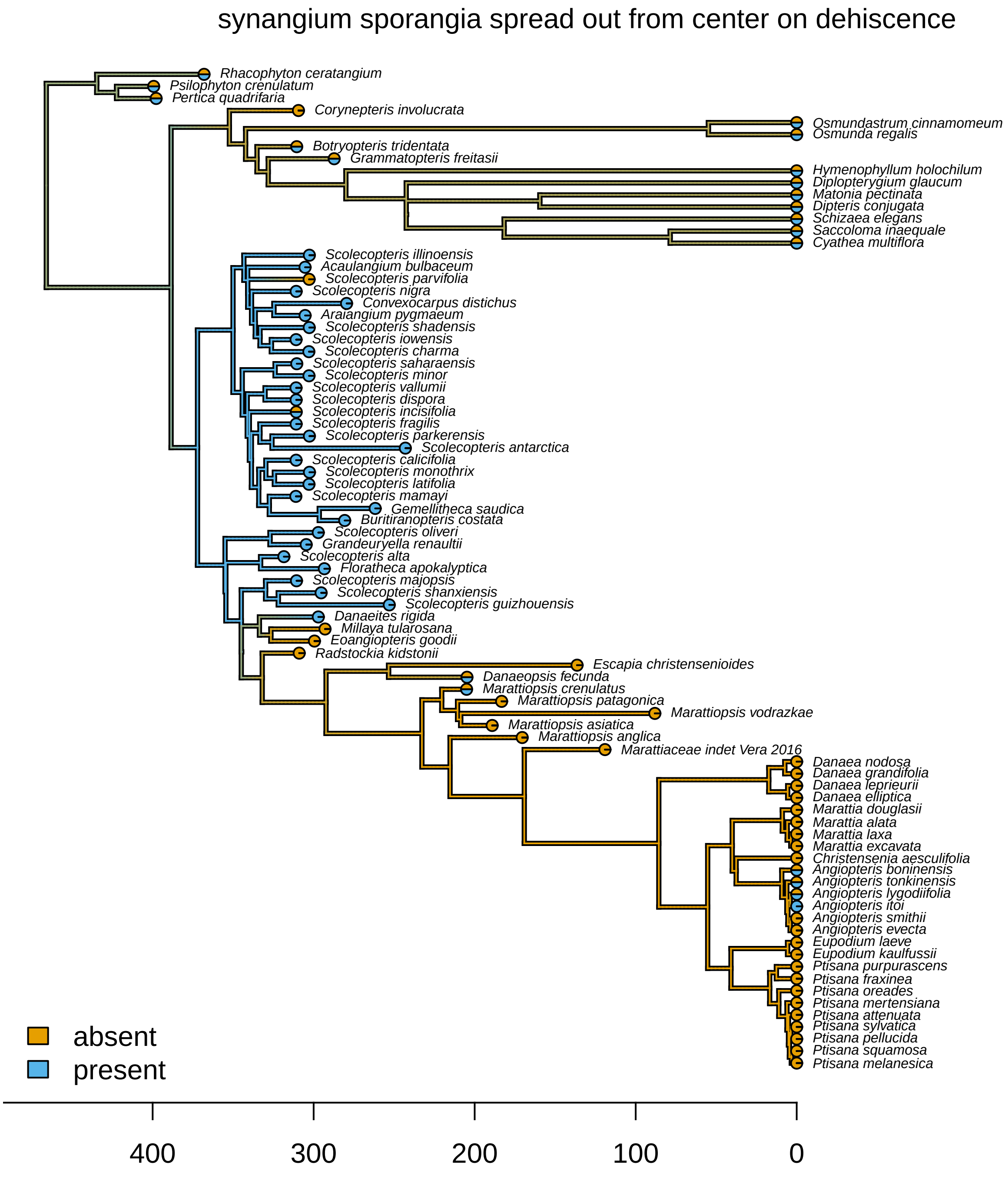


**Figure S73: Stochastic map for character 77: synangium sporangia spread out from center on dehiscence.**

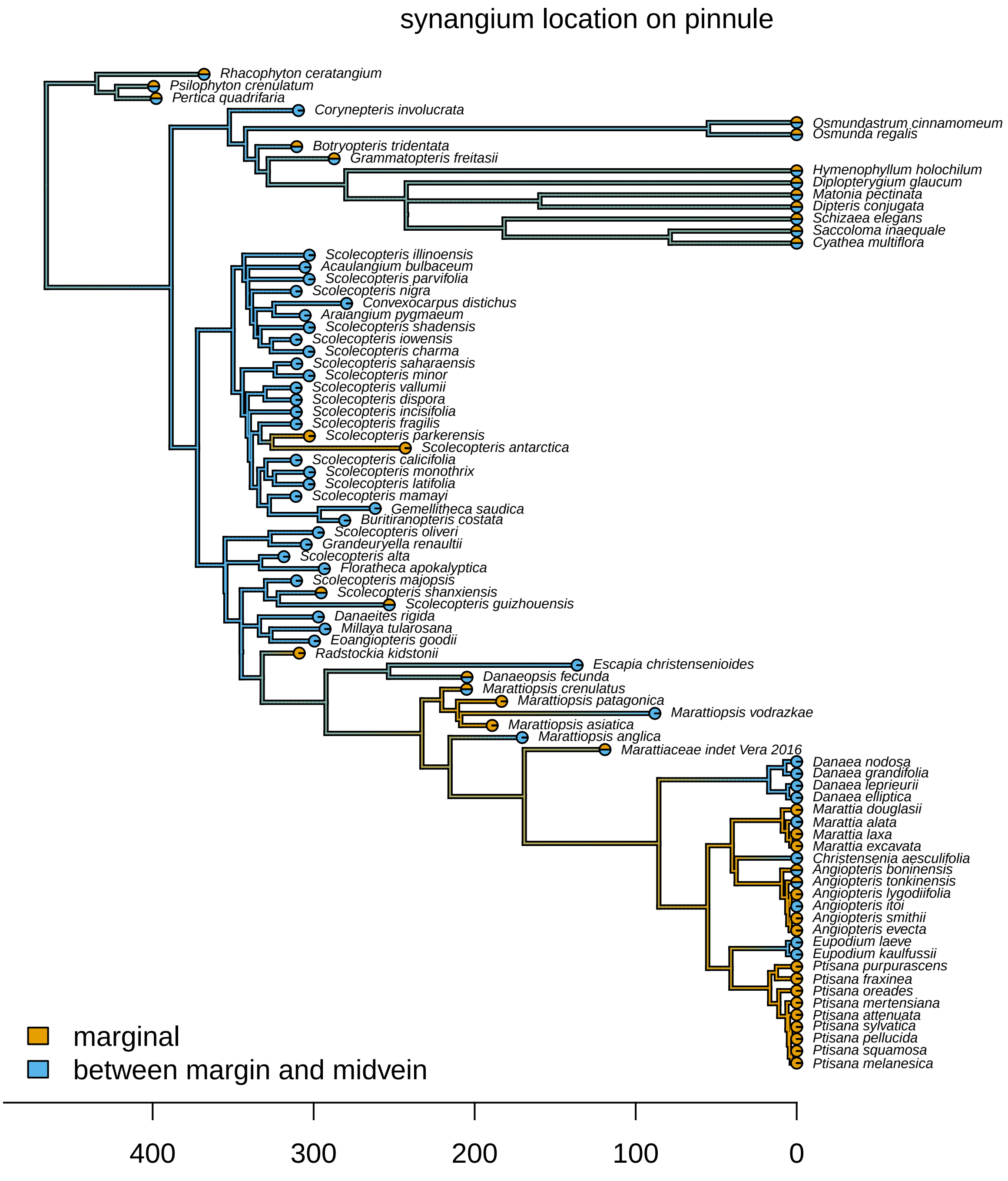


**Figure S74: Stochastic map for character 78: synangium location on pinnule.**

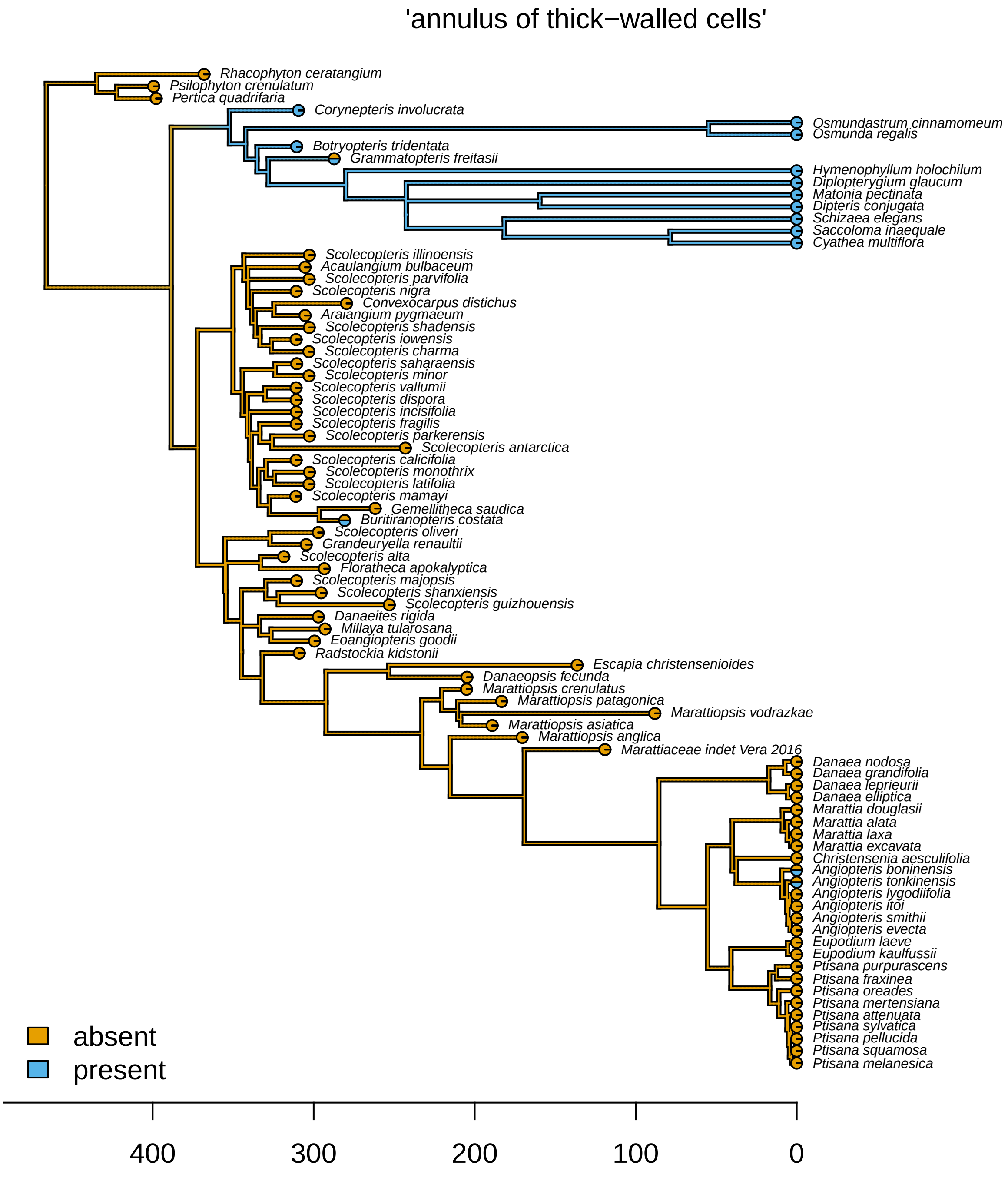


**Figure S75: Stochastic map for character 87: annulus of thick-walled cells.**

#### S§6.11 Dated vs. Non-Dated Topologies

To understand the discrepancies between the topology we inferred compared to those from previous work, we performed a non-dated phylogenetic analysis using the ancient plants dataset. For these analyses, we assumed the F81 mixture+$\Gamma$ morphological transition model, but used a non-dated tree model where where the tree topology was drawn from a uniform prior distribution and molecular branch lengths were drawn from an exponential distribution. To mimic the assumption that rates of morphological and molecular evolution are proportional (as in the ancient plants analysis), we included an additional rate multiplier, $\beta_m$, that scaled the morphological branch lengths relative to the molecular branch lengths. We then compared the phylogeny tree estimated under the ancient plant analysis to that estimated under this non-dated analysis (Figs. S77, S78) using `cophylo` from the `R` package `phytools` (Revell 2012).

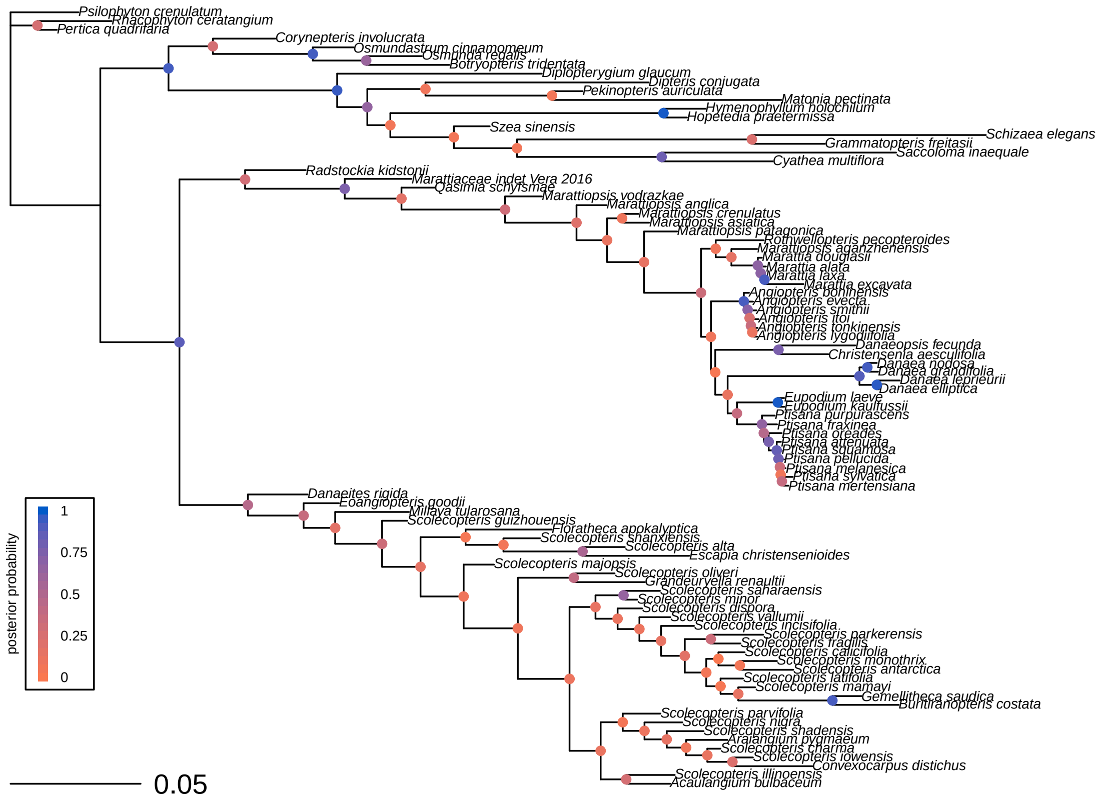


**Figure S76: The maximum clade credibility tree under the non-dated (non-clock) model.** Nodes are colored in proportion to their posterior probability (legend, left).

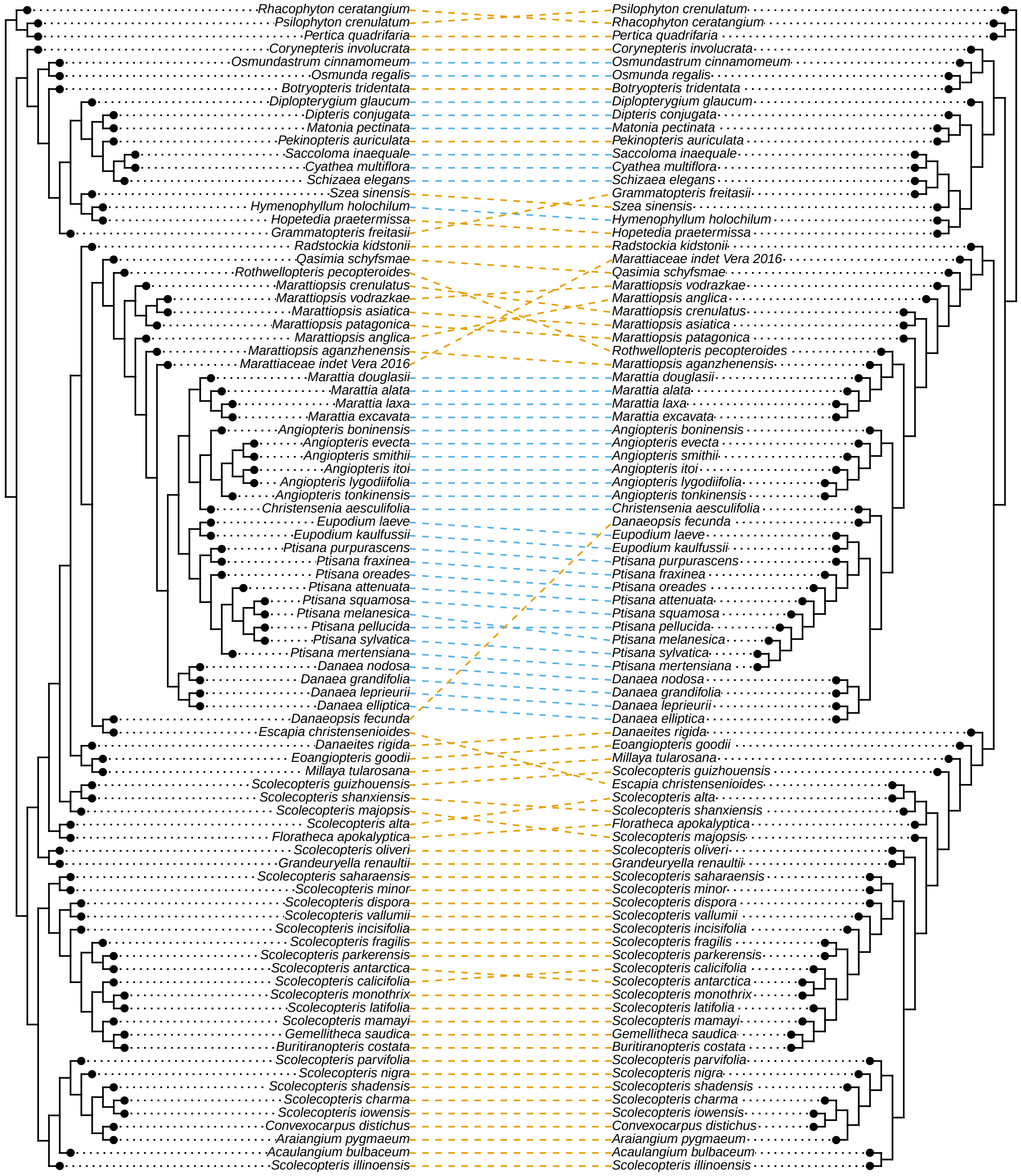


**Figure S77: Topological conflict beween dated and non-dated MCC trees.** We compared the maximum-clade-credibility (MCC) tree under the ancient plant analysis (left) against the MCC tree from the non-dated analysis (right), with branch lengths set arbitrarily to 1. Blue and orange links correspond to extinct and extant taxa, respectively.

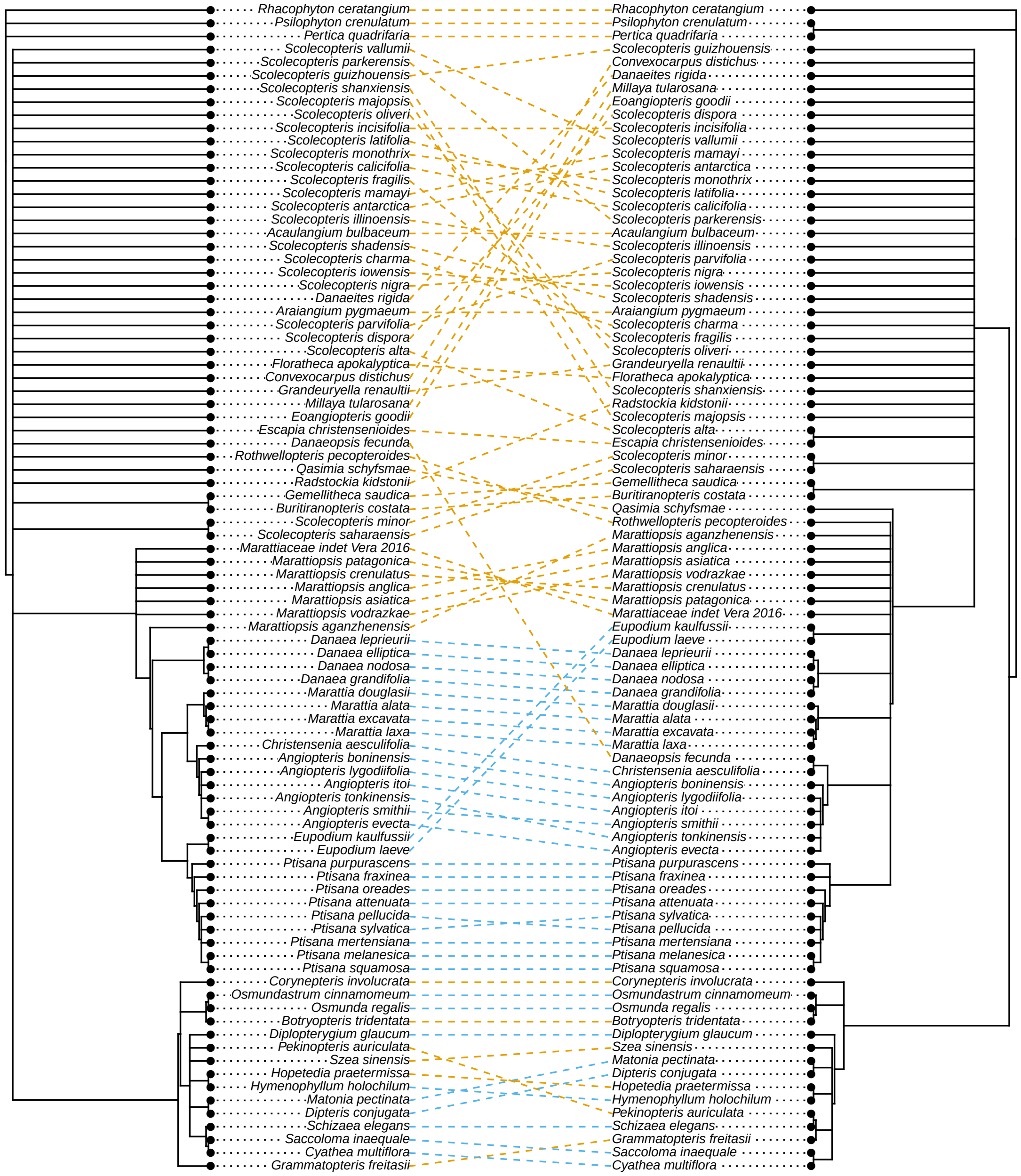


**Figure S78: Topological conflict beween dated and non-dated MRC trees.** We compared the majority-rule consensus (MRC) tree under the ancient plant analysis (left) against the MRC tree from the non-dated analysis (right), with branch lengths set arbitrarily to 1. Blue and orange links correspond to extinct and extant taxa, respectively.
